# Persistent chromatin loops shape gene expression plasticity upon stimulation and restimulation of human neurons

**DOI:** 10.1101/2025.09.30.679661

**Authors:** Abraham J. Waldman, Kenneth Pham, Katelyn R. Titus, Alexandria Nikish, Constin Liu, Han-Seul Ryu, Srikar S. Muppidi, Keerthivasan Raanin Chandradoss, Peibo Xu, Rohan Patel, Ravi Boya, Jennifer E. Phillips-Cremins

**Author notes:** These authors contributed equally to this work (Abraham J. Waldman, Kenneth Pham).

## Abstract

Persistent molecular correlates of long-term memory storage remain an open question. Here, we stimulate and re-stimulate human neurons and use multi-modal single-nucleus technologies to query DNA methylation, higher-order chromatin folding, and gene expression. We find enduring traces of activity-gained and activity-lost chromatin loops. Genes anchoring persistent activity-gained loops exhibit activity-upregulated expression, whereas persistent activity-lost loops anchor activity-downregulated genes that remain repressed five days post-stimulation. CTCF-bound looped enhancers and promoters are refractory to activity-dynamic DNA methylation. Looped enhancers bound by CTCF can exhibit memory of activity-induced histone modifications and persistent expression of activity-upregulated genes. Upon second stimulation, activity-upregulated genes are robustly re-induced when unlooped but remain nonresponsive at persistent loops akin to habituation. Activity-independent gene expression can be downregulated when unlooped but protected from homeostatic downscaling when anchored in persistent loops. Our data reveal long-term genome folding persistence linked to plasticity of activity-dependent gene expression during recall in human neurons.

**Structured Abstract:** *Introduction:* A long-standing question in neuroscience is how memories of previous experiences are stored in the mammalian brain to facilitate recall over the lifetime of an individual. Classic models of memory posit that both synapse-specific events and cell-wide transcriptional programs are required for encoding, consolidation, and long-term storage of memory (*1, 2*). Learning involves synapse strengthening, and synapse weakening has been linked to memory loss (*3, 4*). Nascent transcription and protein synthesis also occur in response to neural activation *in vitro* and *in vivo* (*5-7*). A leading hypothesis, the synaptic tagging and capture model, asserts that specific synapses are biochemically marked during encoding and newly made cell-wide RNA/proteins act specifically on tagged synapses to maintain long-term memory (*8*). Multiple historic studies have pursued the identification of proteins linked to potentiated synapses (*9-13*), but few explore the possibility of persistent, activity-dependent patterns of DNA, chromatin, higher-order chromatin folding, or RNA in human models of long-term memory. Over the last decade, the molecular technique of Chromatin-Conformation-Capture has been employed to discover that the mammalian genome folds into thousands loops of (*5, 14-28*). Loops bring distal non-coding cis regulatory elements into contact with their target genes to influence gene expression (*29*). They form via the processive motion of the cohesin ring along chromatin until it stalls at the architectural protein CTCF, thus extruding out the intervening DNA (*22, 30-32*). Loops can markedly reconfigure during lineage commitment in development and in response to genetic perturbations, and their formation is critical for spatiotemporal regulation of expression *(22, 30, 33-41)*.

*Rationale:* Multiple recent works suggest that loops connecting activity-dependent enhancers to distal genes and can be induced during neural stimulation *in vitro* and behavior paradigms *in vivo (5, 27, 42, 43)*. Genetic elimination of the architectural proteins CTCF and cohesin prior to learning substantially impairs memory encoding across multiple behavior tasks *in vivo* (*42, 44, 45*). Cohesin-mediated loops are necessary for the establishment of new gene expression programs in post-mitotic neurons, including the upregulation of genes encoding axon guidance, dendritic spine morphology, and synaptic plasticity during neuron maturation *in vivo* and activity-dependent gene expression during neural stimulation *in vitro (27).* Fear conditioning recruits epigenetically plastic neurons to form the memory engram (*46*) and induces chromatin accessibility changes at enhancers that might persist at least five days after fear conditioning (*43*). Together, these data provide the rationale for our hypothesis that key structural features of higher-order chromatin folding could be persistent long after the exposure and removal of a pharmacological, environmental, or behavioral stimulus that causes neurons to fire action potentials.

*Results:* Here, we apply cutting-edge single-nucleus multi-modal technologies toward the goal of ascertaining the extent to which there are enduring traces of higher-order chromatin folding, DNA methylation, histone modifications at activity-dependent enhancers, and/or mRNA levels in single human neurons upon exposure to a repetitive stimulation and re-stimulation paradigm. We find that pharmacological stimulation of human induced pluripotent stem cell (iPSC)-derived neurons induces chromatin loop plasticity, including persistent activity-gained (PG) and persistent activity-lost (PL) loops that are dependent on CREB phosphorylation. We also query activity-dependent patterns of DNA methylation genome-wide and demonstrate that the majority of looped enhancers and promoters are negligibly differentially methylated in human neurons in response to pharmacological stimulation. CTCF binding to looped enhancers and promoters further protects such regulatory elements from activity-stimulated changes in DNA methylation. What are the potential functional effects of persistent chromatin loops? We find that activity-lost persistent loops anchor activity-downregulated genes that remain repressed five days post-stimulation. By contrast, activity-gained persistent loops are enriched for activity-induced gene expression at 2-5 hours post-stimulation, but mRNA levels resume baseline levels by five days after removal of the stimulus. A subset of promoter-enhancer loops bound by CTCF can exhibit enduring traces of the histone modification H3K27ac. Promoters persistently looped to CTCF-bound persistent activity-induced enhancers can exhibit memory of activity-upregulated gene expression for at least five days after stimulation. Finally, we re-stimulated our human neurons five days after the original stimulation event and assessed gene expression with single-nucleus RNA-seq. Upon second stimulation, we unexpectedly find that activity-upregulated genes can be robustly re-induced when unlooped but remain nonresponsive at persistent loops akin to habituation. We also unexpectedly find that activity-independent, invariant gene expression can be downregulated at second stimulation when unlooped but protected from homeostatic downscaling when anchored in persistent loops.

*Conclusion:* Taken together, our data reveal enduring traces of higher-order chromatin loops in human neurons after activity-simulation and their link to gene expression plasticity and habituation during re-stimulation and re-stimulation of human neurons. Our work sheds light on the role for chromatin and gene expression in an important unsolved paradox in neuroscience: How is information encoded in neural circuits on the timescale of years despite the rapid turnover of synaptic proteins/RNAs in hours to weeks? Persistent chromatin loops shape plasticity in cell-wide RNA levels, thus laying the foundation for future studies linking genes identified here to their subcellular localization and possible functional role at the synapse.

## Main Text

A central mystery in neuroscience is how neurons store memories at the cellular, subcellular, and molecular level. Classic theories posit that memories are stored in the synaptic connections between neurons (*8, 47*). Moreover, activity-dependent synapse-to-nucleus signaling cascades drive nascent transcription and protein synthesis in response to neural activation (*5-7*). However, nascently formed synaptic connections retract over time (*48*), synapse strengthening has not yet been reported past several weeks *in vivo* (*49*), and the half-life of most RNAs and proteins is only days to weeks (*50, 51*). Therefore, we and others have long hypothesized that enduring molecular traces in either the nucleus or synapse or both may mechanistically contribute to long-term memory storage (*52-54*).

Chromatin provides a candidate substrate that could encode such an enduring trace. Neuronal activation through *in vitro* pharmacological stimulation and *in vivo* behavioral paradigms drives widespread changes in long-range chromatin looping interactions (*5, 27, 43, 55, 56*). Degradation of the architectural proteins CTCF and cohesin prior to and during training substantially impairs memory encoding across multiple behavior tasks *in vivo* (*42, 44, 45*). Cohesin-mediated loops are necessary for the establishment of new gene expression programs in post-mitotic neurons, including the upregulation of genes encoding axon guidance, dendritic spine morphology, and synaptic plasticity during neuron maturation *in vivo* and activity-dependent gene expression during neural stimulation *in vitro (27).* However, the persistence of activity-induced changes in chromatin folding and the link to gene expression during memory recall has remained unclear.

Here, we apply single-nucleus multi-modal technologies to ascertain the extent to which there are enduring traces of higher-order chromatin folding, DNA methylation, or histone modifications at activity-dependent enhancers, or mRNA levels in single human neurons upon exposure to a repetitive stimulation and re-stimulation paradigm.

## Results

### Human neurons spontaneously fire action potentials and express activity-dependent gene expression and protein markers five weeks into their differentiation from human IPSCs

To study the relationship between neuronal stimulation, higher-order chromatin folding, DNA methylation, and gene expression in human neurons, we used the induced pluripotent stem cell (iPSC) line i^3^N with a stably inserted, doxycycline (Dox) inducible Neurogenin2 (*Ngn2*) transgene at the adeno-associated virus integration site 1 (AAVS1) safe harbor locus of the human WTC11 iPSC line (*57, 58*) (**Fig. 1A, Fig. S1A)**.

**Fig. 1.**
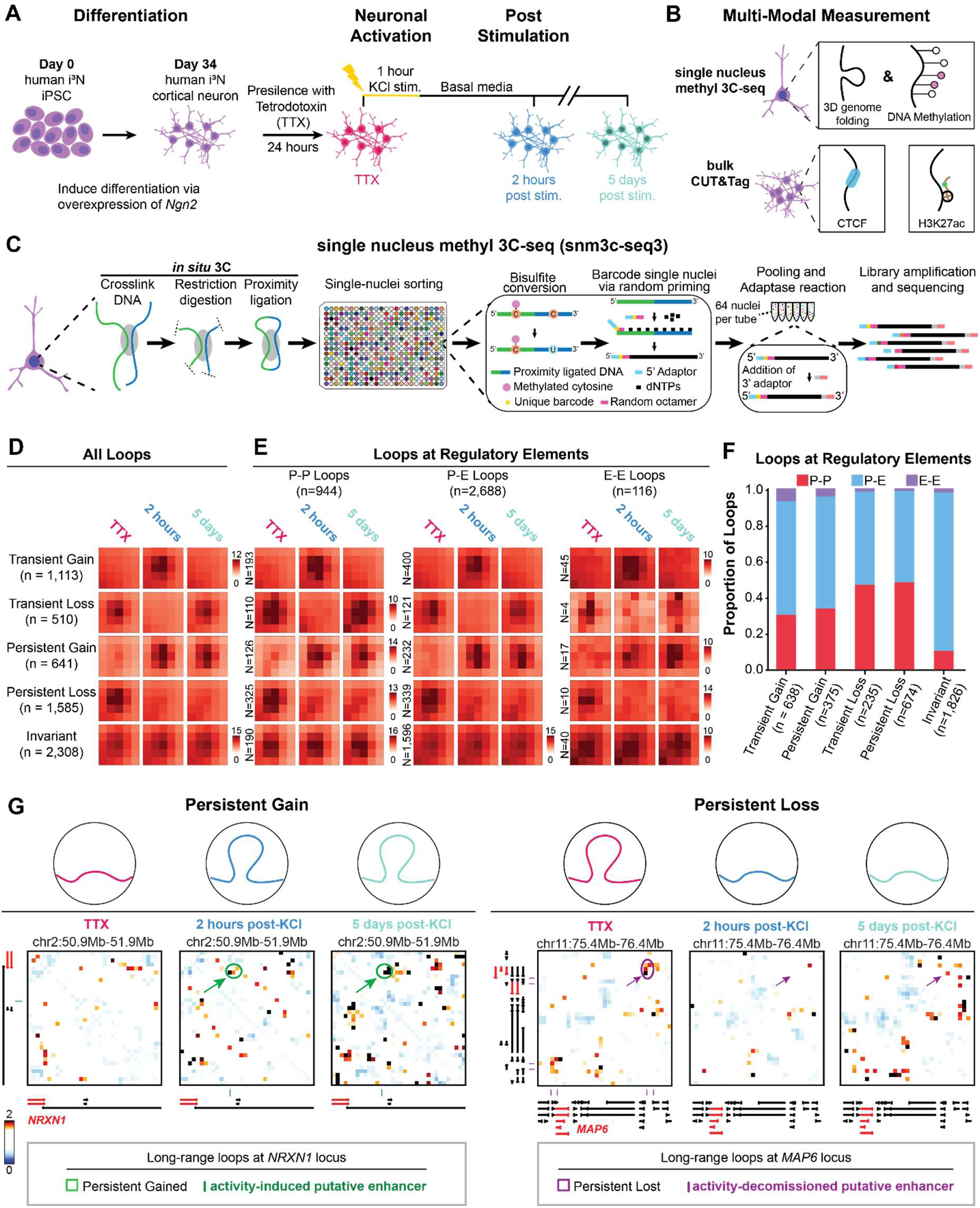
Structural persistence of a subset of activity-induced and activity-decommissioned higher-order chromatin loops at five days post-stimulation of human neurons. **(A-B**) Schematic of the differentiation and stimulation of human neurons along with collection timepoints and assays. **(C)** Single nucleus methyl 3C-seq (snm3c-seq) assays allows for multi-modal inquiry of DNA methylation and higher-order chromatin folding in single neuronal nuclei genome-wide (*64-66*). (**D**) Five loop classes identified in a timecourse of TTX pre-silencing, two hours post-stimulation, and 5 days post-stimulation. **(E-F)** Loop classes can be stratified into promoter-promoter (P-P), promoter-enhancer (P-E), and Enhancer-Enhancer (E-E) loops. Persistently and Transiently Gained PE Loops are anchored by activity-induced putative noncoding enhancers and Persistently and Transiently Lost PE Loops are anchored by activity-decommissioned putative noncoding enhancers. **(G)** Locus specific examples of snm3C-seq heatmaps displaying Observed/Expected interaction frequency of the *NRXN1* promoter connected in a persistently gained loop to a activity-induced enhancer (left) and the *MAP6* promoter connected in a persistently lost loop to an activity-decommissioned enhancer (right) across the timecourse of human neuron pre-silencing, pharmacological stimulation, and persistence upon removal of the stimulus.

To validate our i^3^N model system, we used IF staining and phase contrast imaging. We confirmed that colonies homogeneously express the pluripotency markers POU5F1 and NANOG (**Fig. S1B)**, and exhibit the expected rounded, compact morphologies of hiPSCs (**Fig. S1C).** We next induced differentiation by adding Dox (1µg/mL) to culture media for 3 days and maintained cultures for five weeks post-induction. Using phase contrast microscopy, we observed definitive morphological changes as cells differentiated from hiPSCs to post-mitotic neurons (**Fig. S1C)**. Moreover, homogenous IF staining for the mature layer 2/3 cortical marker CUX1 (*57*), the pan-neuronal marker NEUN, and the microtubule associated protein MAP2 at DIV 35 confirmed that i^3^N homogeneously differentiate into post-mitotic neurons (**Fig. S2A).** We found that the presynaptic marker SYNAPSIN1 and the postsynaptic marker HOMER1 co-localize to MAP2-stained dendrites (**Fig. S2B)** via IF staining. Furthermore, we observed spontaneous action potentials fired by our DIV35 neurons as measured by MicroElectrode Array (MEA) recordings. DIV35 neurons fired synchronously in network bursts, indicating strong interconnectivity among the cultured neurons (**Fig. S2C, Table S3)**. We further confirmed that DIV35 i^3^Ns exhibited the expected activity-dependent changes in protein expression, protein phosphorylation, and gene expression markers previously seen in literature (*5, 27, 56, 59-62*) (**Supplementary Text, Fig. 1A, Fig. S1-S3, Table S3-S4)**. Our results demonstrate that inducible *Ngn2* overexpression via Dox treatment results in the rapid differentiation of human i^3^N iPSCs into a near homogenous population of post-mitotic neurons capable of spontaneously firing action potentials (*63*).

### A subset of activity-induced and activity-decommissioned chromatin loops in human neurons persist up to 5 days after stimulation

Upon validation of our human model system, we next set out to test how neuronal stimulation via membrane depolarization leads to persistent changes in chromatin folding and DNA methylation. We pre-silenced DIV34 neurons with 24 hours of treatment with tetrodotoxin (TTX) (*5*), washed out TTX, stimulated neurons for 1 hour with 50 mM KCl, and returned samples to basal media (**Fig. 1A, Supplementary Text, Supplementary Methods**). To measure chromatin folding and DNA methylation in the same single neurons we conducted fluorescence activated nuclei sorting and multi-modal single-nucleus methyl 3C sequencing (snm3c-seq) (*64-66*) (detailed in **Fig. S4**, **Supplementary Methods, Table S1-S2**). We measured chromatin occupancy of CTCF and the post-translational histone modification H3K27ac using bulk ChIP-seq (*67*) and bulk CUT&Tag (*68, 69*), respectively, and measured mRNA levels with bulk nuclear RNA-seq (**Supplementary Methods, Table S4-7**). We collected samples prior to stimulation with KCl (i.e. pre-silenced with TTX), 2 hours after KCl stimulation, and five days after KCl stimulation (**Fig. 1A-C**, **Supplementary Methods**).

We sequenced >1,000 neurons per condition at a read depth of ∼1 Million reads per neuron and conducted extensive steps to filter only the highest quality 750 cells per condition (**Fig. S5**) and normalize the DNA methylation data at single-cell resolution with gold-standard methods (**Fig. S6**) (*64-66*). We clustered neurons as previously published on genome-wide DNA methylation signal in genome-wide 100 kb bins per cell (*64-66*) and demonstrated homogeneity in excitatory post-mitotic neuron cell state as suggested by our protein and gene expression characterization (**Fig. S7**) and documented homogeneity of the *Ngn2*-differentiated neurons (*63*). After demonstrating neural subtype homogeneity, we normalized single-neuron Chromosome-Conformation-Capture data for sequencing depth differences by scaling counts by genomic distance as previously reported (**Fig. S8A**) (*5, 16, 18, 25, 26, 28, 70-72*). We pooled sparse single-neuron matrices into Hi-C pseudobulk matrices from 750 neurons per condition, Knight-Ruiz matrix balanced, and called loops as clusters of pixels with significantly higher interaction frequency compared to local chromatin domain background with using extensively validated computational steps (*5, 16, 18, 25, 26, 28, 29, 70-72*) (**Fig. S8-S10, Supplementary Methods**).

We next set out to classify long-range chromatin loops by their plasticity across the timecourse of human neuron stimulation (**Fig. 1D-H**). We identified five loop classes: (1) Loops that are transiently gained 2 hours after KCl stimulation but that do not persist five days later (Transient Gained Loops; n=1,113), (2) Loops that are transiently lost 2 hours after KCl stimulation but which return to their basal state by five days later (Transiently Lost Loops; n=510), (3) Persistently gained loops which are activity-induced and enduring at five days post-stimulation (Persistently Gained Loops; n=641), (4) loops that are activity-decommissioned and remain lost at five days post-stimulation (Persistent Lost Loops; n=1,585), and (5) loops exhibiting invariantly high interaction frequency across all three timepoints (Activity-invariant Loops; n=2,308) (**Fig. 1D, Table S8-S9)**. We next integrated loop classes with promoters, putative activity-induced non-coding enhancers, and putative activity-decommissioned non-coding enhancers and observed that the majority of activity-dynamic loops connect promoters to activity-dependent enhancers (**Fig. 1E-F, Supplementary Methods, Table S5-6**). Together, our data demonstrate widespread plasticity and persistent traces of chromatin loops genome-wide upon pharmacological stimulation of human neurons.

We next explored whether loci anchoring activity-induced looping changes were associated with physiologically relevant genes. We highlight a persistently gained PE loop connecting the *NRXN1* gene to a distal, noncoding, activity-induced putative enhancer (**Fig. 1G)**. *NRXN1* is necessary for learning and memory in mice (*73*) and in which variants have been associated with autism spectrum disorder (*74*). Similarly, we highlight a persistently lost loop connecting the *MAP6* gene to a distal, noncoding, activity-decommissioned putative enhancer (**Fig. 1G)**, which codes for a protein involved in calmodulin binding and dendritic maintenance (*75*). We additionally observe transiently gained loops at *HMBOX1*, *EMILIN2*, *FAM162A*, and *NAF1* (**Fig. S11**). Our locus-specific examples illustrate that activity-dynamic loops physically connect crucial neuronal and synaptic genes to long-range, distal activity-dependent cis regulatory enhancer elements.

We next sought to assess the signaling pathway through which KCl stimulation might propagate from the synapse to the nucleus to drive activity-dependent plasticity in chromatin looping. Neural stimulation with KCl drives increases in intracellular calcium through L-type calcium channels, leading to a signaling cascade from the synapse to the nucleus that persistently phosphorylates the cAMP-response element binding protein (CREB) (*59, 76, 77*). Increases in intracellular calcium can be blocked by dihydropyridines such as nimodipine (Nim) to prevent CREB phosphorylation (pCREB) and its associated effect on gene expression (**Fig. S3)** (*59, 76*). To assess which activity-induced changes in chromatin looping occur through L-type calcium channels and lead to pCREB, we stimulated neurons at DIV35 with KCl in the presence of Nim and assessed whether looping changes were blocked (pCREB/Nim-dependent) or unaffected (pCREB/Nim-independent) **(Fig. S12A,B)**. We found that the majority of transient and persistently gained loops were pCREB-dependent (**Fig. S12C-D, Table S10)**. A substantial proportion of activity-decommissioned transient and persistently lost loops were also pCREB-dependent (**Fig. S12E-F, Table S11**). Our data indicate that plasticity in chromatin looping upon KCl-mediated stimulation of human neurons can occur through signaling pathways involving CREB phosphorylation activated at the synapse.

### Looped promoters and enhancers bound by CTCF are refractory to activity-dependent changes in methylation of CpG dinucleotides

Perturbations of DNA methyltransferases inhibit memory formation and consolidation in murine models (*78, 79*). Moreover, recent studies have identified activity-induced changes in DNA methylation upon neural stimulation using *in vitro* and *in vivo* model systems (*78-84*), suggesting activity-dependent differentially methylated loci (DML) might interplay with activity-dynamic loops in human neurons.

To assess activity-dependent DMLs in our model system, we analyzed changes in methylation at single CpG resolution 2 hours post-KCl stimulation and 5 days post-KCl stimulation in comparison to the TTX pre-silencing condition (**Supplementary Methods, Fig. S13-S16**). We identified 157,940 activity-induced DMLs genome-wide in the 2 hours post-KCl stimulation condition, including 60,998 transiently hypermethylated DMLs, 52,466 persistently hypermethylated DMLs, 41,134 transiently hypomethylated DMLs, and 3,342 persistently hypomethylated DMLs genome-wide (**Fig. S17**). Our data confirm the existence of activity-dependent DMLs in human neurons and that snm3C-seq in our hands and our computational methodologies are sensitive and specific to detect them.

CTCF has been hypothesized to block DNA methylation at developmental and imprinted promoters (*85-87*). Here, in a postmitotic, activity-induced paradigm, we sought to investigate whether DMLs at promoters and enhancers preferentially occurred at sites with or without CTCF. Specifically, we focused our analysis on promoters with CpG islands, promoters without CpG islands, activity-decommissioned putative enhancers, and activity-induced putative enhancers (**Supplemental Methods, Fig. S18, Fig. 2A**). We strikingly observed that CpG dinucleotides at CTCF-bound promoters and enhancers anchoring loops were refractory to activity-dependent DMLs (**Fig. 2B top, Fig. S18B top**). Looped enhancers and promoters with no CTCF occupancy showed a substantial increase in DML signal compared to CTCF-bound, looped regulatory elements (**Fig. 2B bottom, Fig. S18B bottom**). The fraction of reads with CpG methylation at a particular regulatory element did not clearly trend with the activity-dependent DML patterns (**Fig. 2C, Fig. S18C**). CTCF-bound, looped promoters are unmethylated at baseline and do not gain DNA methylation upon neural stimulation (**Fig. 2B-C, Fig. S18B-C**). Taken together, our data indicate that the large majority of activity-dependent DMLs do not occur and looped promoters and enhancers and further suggests that CTCF occupancy protects looped regulatory elements from dynamic DNA methylation upon human neuron stimulation.

**Fig. 2.**
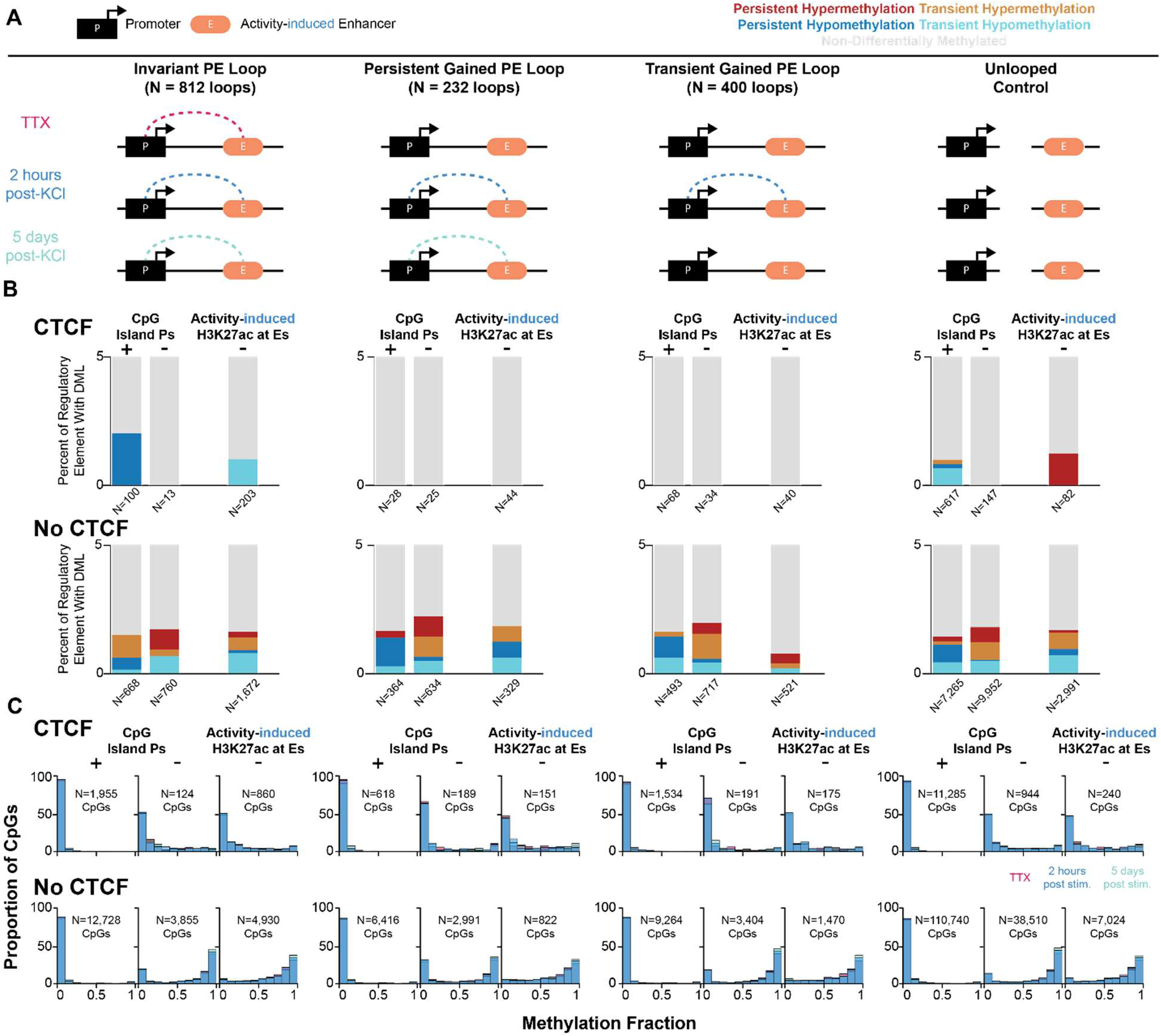
CTCF-bound enhancers and promoters connected in activity-dynamic loops are resistant to activity-dependent DNA methylation. **(A**) From left to right: schematics of the invariant PE loops, persistently gained PE loops, transiently gained PE loops, unlooped controls for putative enhancers that gain H3K27ac upon KCl stimulation (activity-induced enhancers). **(B)** Percentage of regulatory elements with each type of DML for the four looping classes defined in **(A)**. From left to right within a given class: promoters with CpG islands, promoters without CpG islands, and enhancers that gain H3K27ac upon KCl stimulation. Upper row represents regulatory elements bound by CTCF, and lower row represents regulatory elements not bound by CTCF. **(C)** Histograms of methylation fraction for each group defined in **(B)**.

### Persistently lost PE loops associated with activity-downregulated mRNA levels that remain persistently low at five days post-stimulation

We next investigated the link between persistently lost loops and gene expression (*27, 88*). A recent study reported that KCl-stimulation drives both the up and downregulation of thousands of genes in glutamatergic neurons (*56*), but the link to activity-decommissioned loops and the persistence of the loss of the higher-order structural contacts is unknown.

Using bulk nuclear RNA-seq, we identified genes whose expression was activity-downregulated in comparison to TTX either 2 hours (**Fig. 3A)** or 5 hours post KCl stimulation (**Fig. 3B**) (**Supplementary Methods, Table S4, S12**). We identified 2,522 activity-downregulated genes upon 2 or 5 hours KCl treatment compared to TTX pre-silencing with a 5% false discovery rate (FDR) (**Fig. 3A,B**). We further stratified the activity-downregulated genes (so-called blue genes) by their expression post-stimulation (**Fig. 3C, Supplementary Methods**). We identified 1,327 activity-downregulated genes that remained persistently downregulated (blue➔blue) after 5 days post-KCl treatment as defined by a 1.2-fold change (FC) decrease from counts in the TTX condition (**Fig. 3C-D, Table S12**). Our data indicate that a substantial subset of activity-downregulated genes remain persistently low in mRNA levels for at least five days after KCl stimulation.

**Fig. 3:**
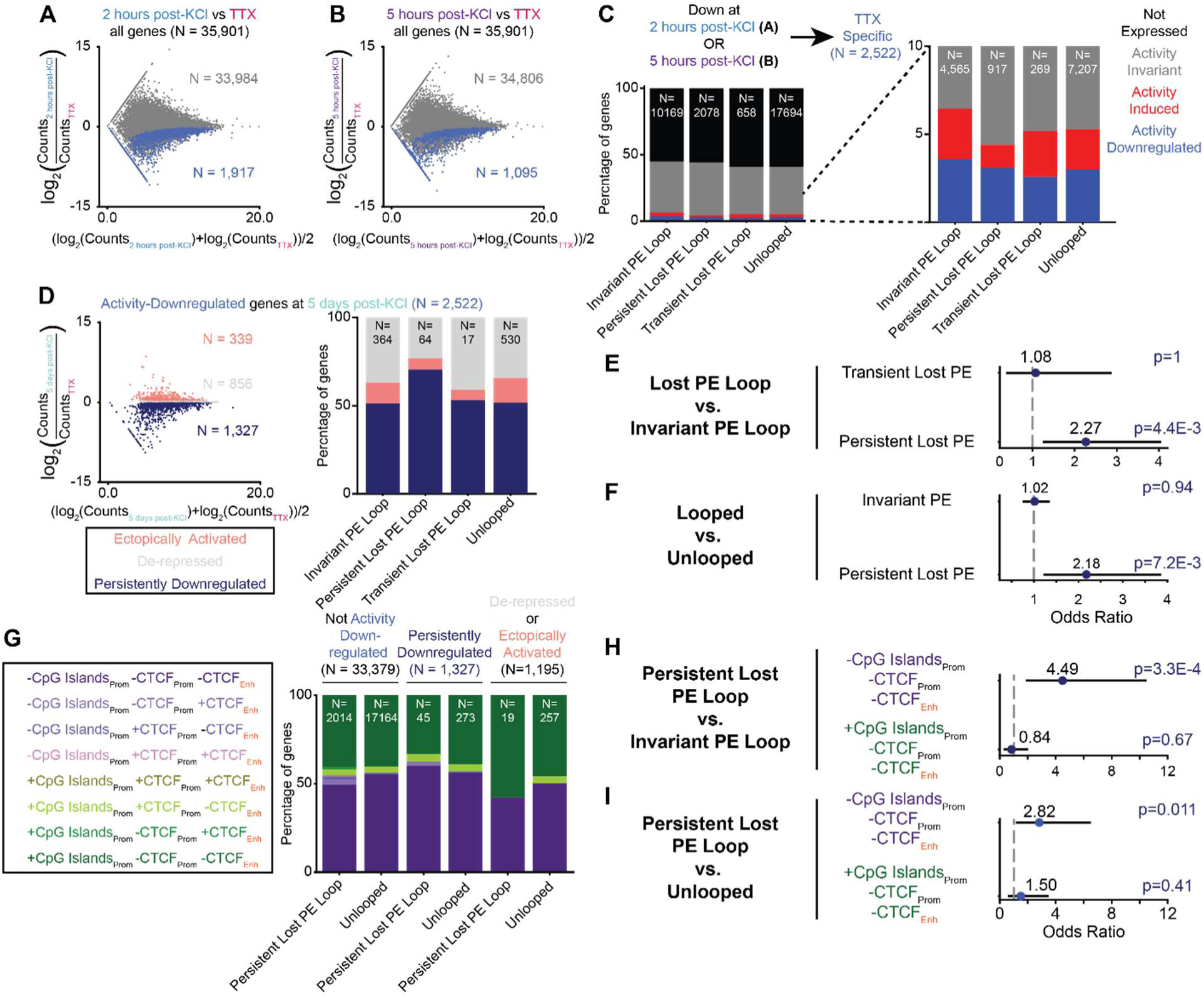
Persistently lost loops connect activity-decommissioned enhancers to promoters exhibiting persistent activity-downregulated gene expression. **(A-B)** M-A plot comparing nuclear RNA-seq at **(A)** 2 hours and **(B)** 5 hours post-KCl to TTX. Genes colored blue represent activity-downregulated genes (q < 0.05, Benjamini-Hochberg multiple-testing correction). **(C)** Proportion of genes not expressed, activity-independent invariantly expressed, activity-upregulated, or activity-downregulated for promoters connected to activity-decommissioned enhancers in invariant PE loops, persistently lost PE loops, transiently lost PE loops, or an unlooped control. **(D)** M-A plot comparing nuclear RNA-seq counts 5 days post-KCl to TTX for all genes identified as activity-downregulated at 2 or 5 hours post-stimulation in **(A)**. Light red represents activity-downregulated genes that are ectopically upregulated at 5 days post-KCl (FC TTX / 5 days post-KCl < 1/1.2), grey represents activity-downregulated genes that are de-repressed at 5 days post-KCl (1/1.2 < FC < 1.2), and dark blue represents activity-downregulated genes that are persistently downregulated (FC > 1.2). **(E-F)** Odds ratio and Pvalue (Fisher’s Exact Test) for persistent downregulation of activity-decommissioned gene expression at **(E)** persistently lost PE loops vs. invariant PE loops or **(F)** persistently lost PE loops vs. unlooped. **(G)** Proportion of loops with CTCF occupancy at enhancers and promoters with and without CpG islands. **(H-I)** Odds ratio and Pvalue (Fisher’s Exact Test) for persistent downregulation of activity-decommissioned gene expression at **(H)** persistently lost PE loops vs. invariant PE loops with and without CpG islands or **(I)** persistently lost PE loops vs. unlooped with and without CpG islands.

We next integrated promoters anchoring loop classes and unlooped with gene expression. To avoid conflating phenomena, we removed all promoters coinciding with DMLs from our analysis. We found a striking enrichment between persistently lost PE loops and activity-downregulated genes that remain persistently downregulated post-stimulation compared to transiently lost PE loops and unlooped controls (**Fig. 3D-F**). We further parsed the promoters by the presence or absence of CTCF or CpG islands and enhancers by the presence or absence of CTCF. We found that promoters and enhancers devoid of CTCF occupancy and without CpG islands made up the large majority of regulatory elements connected in persistently lost loops linked to persistently downregulated gene expression **(Fig. 3G-I)**. Thus, we uncover a strong correlation between persistence in gene expression downregulation and persistent loss in structural contact loss between enhancers and promoters, with the strongest effect at regulatory elements without CTCF or CpG islands.

### Persistently gained loops are anchored by activity-induced and activity-independent invariantly expressed genes

We next set out to assess the gene expression trends that occur at persistently gained PE loops. Recent murine studies have established that activity-induced chromatin looping at distal non-coding enhancers governs gene expression levels of the secondary response gene *BDNF* at six hours post-stimulation (*5, 27*). The link between persistent changes in activity-induced chromatin looping, and persistent, activity-induced changes in gene expression remains unanswered.

Using bulk nuclear RNA-seq, we first identified genes that exhibited increased mRNA levels (activity-upregulated) in comparison to TTX either 2 or 5hours post KCl stimulation (**Fig. 4A-B**). We identified 2,203 activity-upregulated genes compared to TTX pre-silencing (**Fig. 4A-B, Table S12**). We find that persistently gained PE loops anchored by activity-induced enhancers were enriched for activity-induced genes in comparison to unlooped controls (**Fig. 4C,D**). We next identified cohorts of activity-induced genes that were still persistently on (Red➔Red) compared to deactivated (Red➔Grey) or ectopically repressed (Red➔Blue) at 5 days post-KCl stimulation (**Fig. 4E, Table S13**). We found that more than half of activity-induced genes maintained persistent upregulation 5 days post-KCl in comparison to TTX (**Fig. 4E**). However, persistently gained PE loops did not show an enrichment for persistently upregulated (Red➔Red) genes expression against invariant PE loops, transiently gained PE loops, or unlooped controls (**Fig. 4E**). Thus, while activity-induced genes are enriched at persistently gained loops, they do not show sustained high expression 5 days post-KCl stimulation.

**Figure 4:**
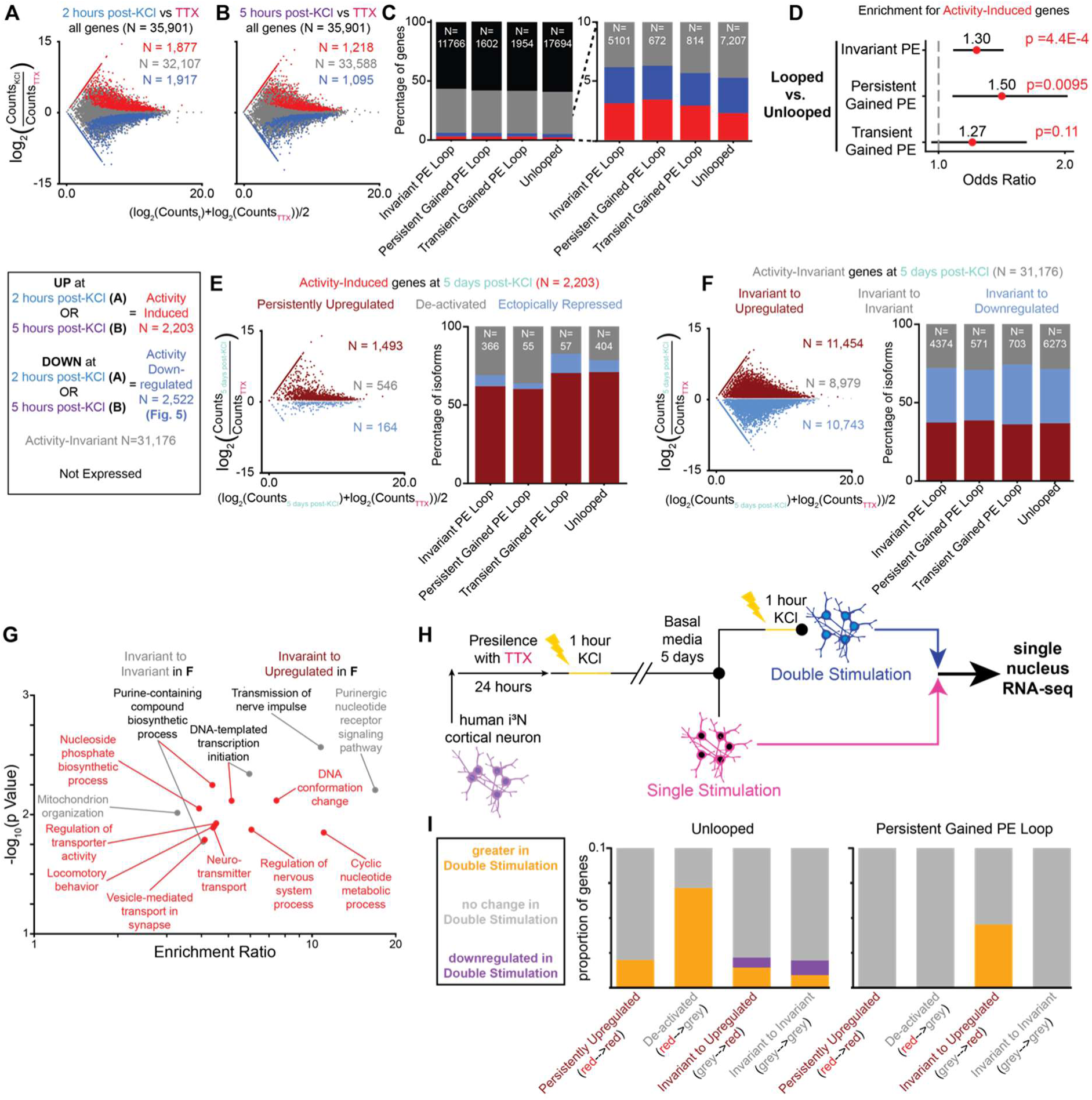
Persistently gained promoter-enhancer chromatin loops shape habituation of activity-induced and activity-independent gene expression upon restimulation. **(A-B)** M-A plot comparing nuclear RNA-seq at **(A)** 2 hours and **(B)** 5 hours post-KCl to TTX. Genes colored red and grey represent activity-upregulated and activity-independent invariantly expressed genes (q < 0.05, Benjamini-Hochberg multiple-testing correction), respectively. **(C)** Proportion of genes not expressed, activity-independent invariantly expressed, activity-upregulated, or activity-downregulated for promoters connected to activity-induced enhancers in invariant PE loops, persistently gained PE loops, transiently gained PE loops, or an unlooped control. **(D)** Odds ratio and Pvalue (Fisher’s Exact Test) for activity-induced gene expression at persistently gained PE loops vs. unlooped. **(E-F)** M-A plot comparing nuclear RNA-seq counts 5 days post-KCl to TTX for all genes identified as **(E)** activity-upregulated or **(F)** activity-independent at 2 or 5 hours post-stimulation in **(C)**. **(E)** Dark red represents activity-upregulated genes that are persistently upregulated at 5 days post-KCl (FC TTX / 5 days post-KCl < 1/1.2) (Red➔Red), grey represents activity-upregulated genes that are de-activated at 5 days post-KCl (1/1.2 < FC < 1.2) (Red➔Grey), and blue represents activity-upregulated genes that are ectopically downregulated (FC > 1.2) at 5 days post-KCl (1/1.2 < FC < 1.2) (Red➔Blue). **(F)** Dark red represents activity-invariant genes that are persistently upregulated at 5 days post-KCl (FC TTX / 5 days post-KCl < 1/1.2) (Grey➔Red), grey represents activity-invariant genes that remain invariant at 5 days post-KCl (1/1.2 < FC < 1.2) (Grey➔Grey), and blue represents activity-invariant genes that are ectopically downregulated (FC > 1.2) at 5 days post-KCl (1/1.2 < FC < 1.2) (Grey➔Blue). **(G)** Schematic of the human neuron re-stimulation experimental paradigm. **(H)** Gene Ontology analysis of genes in Grey➔Grey and Grey➔Red gene classes at persistently gained loops. (**I)** Proportion of unlooped (left) and persistent gained PE loop (right) genes stratified by difference in proportion of nuclei expressing gene. Yellow represents genes in which a higher percentage of nuclei express the gene in the Double vs. Single Stimulation condition. Grey represents genes in which the same percentage of nuclei express the gene in the Double vs. Single Stimulation condition and the Single Stimulation condition. Purple represents genes in which a lower percentage of nuclei express the gene in the Double vs. Single Stimulation condition. Unlooped genes: red-->red (N=100), red-->grey (N=29), grey-->red (N=624), grey-->grey (N=333). Persistently gained PE loops genes: red-->red (N=4), red-->grey (N=1), grey-->red (N=33), grey-->grey (N=12).

We next considered genes with activity-independent, invariant gene expression (activity-invariant expression) across all key TTX and KCl stimulation timepoints. We found a roughly equal distribution of activity-invariant to upregulated (Grey➔Red), activity-invariant to invariant (Grey➔Grey), and activity-invariant to downregulated (Grey➔Blue) gene expression patterns by five days post-stimulation across looping classes and unlooped control genes (**Fig. 4F, Table S13**). Ontology of activity-invariant genes at persistently gained PE loops that either stay invariantly expressed (Grey➔Grey) or are upregulated (Grey➔Red) five days post-stimulation revealed strong enrichment for pathways involved in synaptic signaling and neuronal function (**Fig. 4G**). Our results uncover the unexpected observation that persistent PE loops anchor activity-independent, invariantly expressed genes with ontology suggesting importance for neurophysiological pathways.

### Activity-invariant genes connected to activity-induced enhancers via persistent loops exhibit chromatin habituation upon second restimulation of human neurons

Repetitive exposure to a common experience or stimulus contributes to the encoding and storage of memory. To simulate repetitive exposure *in vitro*, we re-stimulated our human neurons a second time with KCl five days after the original treatment (**Fig. 4H**). We used single-nucleus RNA-seq (snRNA-seq) to compare the proportion of cells expressing mRNA in single-stimulated versus double-stimulated neurons. To ensure the quality, rigor, and reproducibility of our snRNA-seq analysis, we filtered out low signal genes across condition, poor quality nuclei, and we normalized data across cells and conditions to account for technical artifacts (**Fig. S19, S20, Supplementary Methods**).

We examined multiple gene expression classes for their single cell expression phenomena after second stimulation, including those that are (1) activity-invariant upon first KCl stimulation and become upregulated at five days post-single stimulation (Grey➔Red), (2) activity-invariant upon first KCl stimulation and remain invariant at five days post-single stimulation (Grey➔Grey), (3) activity-induced upon first KCl stimulation and remain persistently upregulated at five days post-single stimulation (Red➔Red), and (4) activity-induced upon first KCl stimulation but then return to baseline expression levels at five days post-single stimulation (Red➔Grey) (**Table S13**).

Two major observations emerged from our analysis of single-cell gene expression phenomena upon second KCl stimulation (**Fig. 4I**). Consistent with expectation, activity-upregulated genes (Red➔Grey or Grey➔Grey from first stimulation) can be re-induced after second stimulation when unlooped. However, when connected in persistently gained loops to activity-induced enhancers, we unexpectedly observe that the same classes of Red➔Red or Red➔Grey activity-upregulated genes exhibit a pattern of habituation in which they are not re-induced during second stimulation. We also unexpectedly observe that activity-independent, invariantly expressed synaptic genes (Grey➔Grey or Grey➔Red from first stimulation) can be downregulated only upon second stimulation when unlooped. By contrast, when anchoring persistently gained loops, such Grey➔Grey or Grey➔Red genes unexpectedly escape from downscaling and can even become activity-upregulated at second stimulation. Our data reveal that persistent activity-gained loops exhibit two types of compensatory gene expression phenomena, both protection from downscaling and habituation, after repeated pharmacological stimulation.

### CTCF binding is linked to persistent H3K27ac at activity-induced enhancers anchoring persistently gained loops at five days post-stimulation in human neurons

A notable observation from exploring the functional effects of persistently lost and persistently gained loops on gene expression is that in large part the loops were devoid of CTCF occupancy (**Fig. 3G, Fig. S21**). Having demonstrated the functional impact of non-CTCF loops, we also investigated the functional role for CTCF-mediated loops on activity-dependent gene expression in human neurons. We stratified putative activity-induced enhancers at our various PE loop classes into those with and without CTCF and assessed the H3K27ac signal changes over time, as histone modifications have been hypothesized as a substrate for cellular memory (*89, 90*). In support of this hypothesis, murine studies have demonstrated that histone lysine acetylation specifically contributes to memory formation in learning paradigms (*46, 91-94*). In a study using fear conditioning, mouse engram cells exhibited a persistent increase in H3K27ac at four enhancers tested, even in the absence of continued stimulus (*43*). We therefore assessed whether activity-induced changes in H3K27 acetylation persisted five days after stimulation genome-wide (**Fig. 5**).

**Fig. 5:**
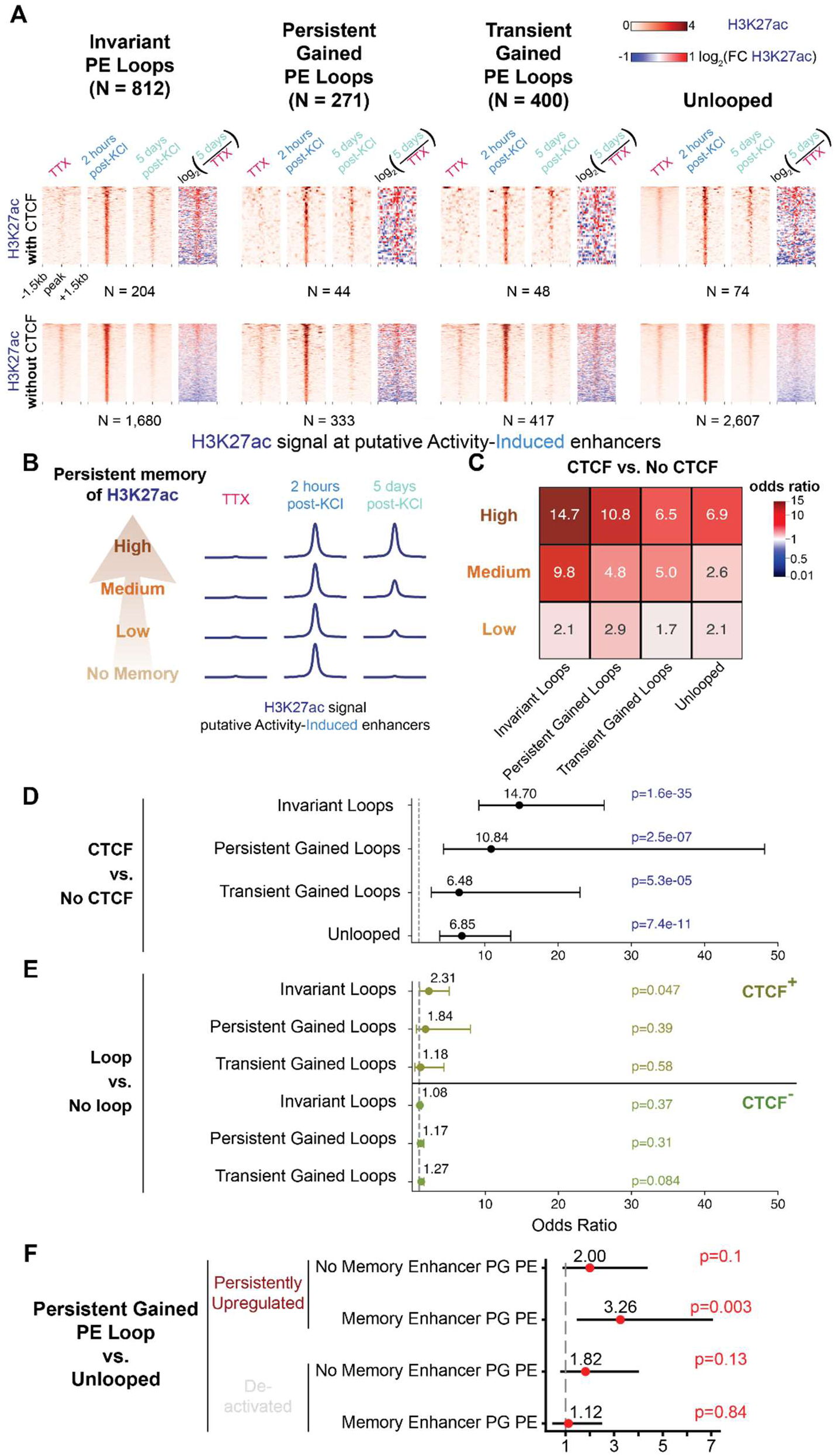
Persistent histone modification H3K27ac at putative non-coding activity-induced enhancers bound by CTCF loop to promoters with persistent activity-upregulated gene expression. **(A)** Heatmaps of H3K27ac CUT&Tag at non-coding activity-induced enhancers, comparing TTX, 2 hours and 5 days post-stimulation, and log2(fold change) of 5 days/TTX. **(B)** Schematic representation of four strata of persistent memory of activity-induced H3K27ac peaks. **(C-D)** CTCF-bound versus non-CTCF-bound activity-induced enhancers tested for association with persistent traces of H3K27ac versus no H3K27ac 5 days later. Odds ratios and Pvalues (Fisher’s Exact Test). **(C)** Odds ratios across a titration of the degree of H3K27ac persistence at CTCF and no CTCF bound enhancers for each loop class and unlooped enhancers are shown. **(D)** Odds ratios, bootstrapped 95% confidence intervals, and Pvalues for tests for each loop class and unlooped enhancers shown. **(E)** Looped versus unlooped activity-induced enhancers tested for association with persistent traces of H3K27ac versus no H3K27ac 5 days later. Odds ratios, bootstrapped 95% confidence intervals, and Pvalues (Fisher’s Exact Test) for tests for each loop class vs. unlooped enhancers after stratifying by CTCF occupancy by enhancers are shown. **(F)** Persistent gain looped versus unlooped activity-induced genes tested for association with persistent upregulation 5 days later (top). Persistent gain looped versus unlooped activity-induced genes tested for association with de-activation 5 days later (bottom). Odds ratios, bootstrapped 95% confidence intervals, and Pvalues (Fisher’s Exact Test) for tests for genes looped to memory and non-memory enhancers are shown.

We focused our analysis on putative activity-induced enhancers, defined as non-coding H3K27ac peaks gained de novo at 2 hours after KCl stimulation and not present in neurons only treated with TTX (**Fig. 5A**). We unexpectedly observed that a subset of activity-induced enhancers binding retained an enduring trace of H3K27ac signal at 5 days after KCl stimulation (**Fig. 5A**). We noticed that those enhancers with possible memory of H3K27ac were also strongly bound by CTCF. By contrast, activity-induced enhancers not bound by CTCF largely showed H3K27ac signal that returned to baseline levels at 5 days post-stimulation (**Fig. 5A**). Our data suggest that putative activity-induced enhancers with CTCF binding are more likely to retain persistent traces of activity-dependent H3K27ac compared to those without CTCF.

To test the relationship between CTCF binding and persistent H3K27ac memory, we stratified the putative activity-induced enhancers into three levels of persistence (high, medium, and low) and no persistence (no memory) based on the H3K27ac signal in neurons 5 days post KCl stimulation (**Fig. 5B, Supplementary Methods**). We then evaluated the association between CTCF binding at the putative activity-induced enhancers for each class of loop and persistent versus non-persistent H3K27ac (**Supplementary Methods**) (**Fig. 5C**). We demonstrate a statistically significant association between CTCF occupancy and H3K27ac persistence at activity-induced enhancers (**Fig. 5C-D**). The Odds Ratio effect size titrates by the strength of the H3K27ac persistence (**Fig. 5C**). Our analyses confirm our qualitative observations that CTCF binding is associated with a higher probability of enduring traces of H3K27ac at activity-induced enhancers at 5 days post-stimulation.

We assessed whether looping alone was associated with persistent H3K27ac at activity-induced enhancers. Using the most stringent persistent H3K27ac strata, we compared the looped enhancers from each looping class against unlooped enhancers matched by CTCF occupancy status. We did not find consistent, significant associations with persistent H3K27ac at activity-induced enhancers and looping (**Fig. 5E**). Our data suggest that CTCF occupancy at activity-induced enhancers regardless of looping or unlooped status contributes to persistence in H3K27ac at activity-induced enhancers.

Finally, we asked if CTCF-bound activity-induced enhancers with enduring traces of H3K27ac have an effect on mRNA expression levels of the promoters connected in loops. We observed a significant enrichment between persistently gained loops with enduring traces of H3K27ac at activity-induced enhancers and activity-induced genes upon first KCl stimulation that remain persistently upregulated at five days post-single stimulation (Red➔Red) (**Fig. 5F**). No other gene expression class (**Figs. 3-4**) was notably enriched at persistently gained loops with persistent activity-induced enhancers. Taken together, our data reveal a link among persistent loops, enduring traces of H3K27ac at activity-induced enhancers, and persistent upregulation of activity induced gene expression.

## Discussion

Here, using an *in vitro* human model of hiPSC-derived neurons, we identify a link between pharmacological stimulation of action potential firing and persistent changes to chromatin and gene expression. We identify activity-induced and activity-decommissioned chromatin loops connecting promoters to distal putative enhancers marked by noncoding activity-dependent H3K27ac signal. Persistently lost loops are associated with sustained, activity-downregulation of gene expression (**Fig. 6A, top**), whereas persistently gained loops are enriched for activity-induced gene expression that resumes baseline levels by five days post-stimulation (**Fig. 6A, bottom**). We identify a subset of persistently gained promoter-enhancer loops that maintain enduring traces of the H3K27ac histone mark at enhancers (**Fig. 6B**). The promoters looped to enduring activity-induced enhancers exhibit activity-induced gene expression that remains persistently high for five days after removal of the initial stimulation (**Fig. 6B, top**). Together, our results indicate that higher-order chromatin structure is plastic upon pharmacological stimulation of action potentials, and that persistent traces of structural chromatin memory exist for at least five days after removal of the stimulus.

**Fig. 6:**
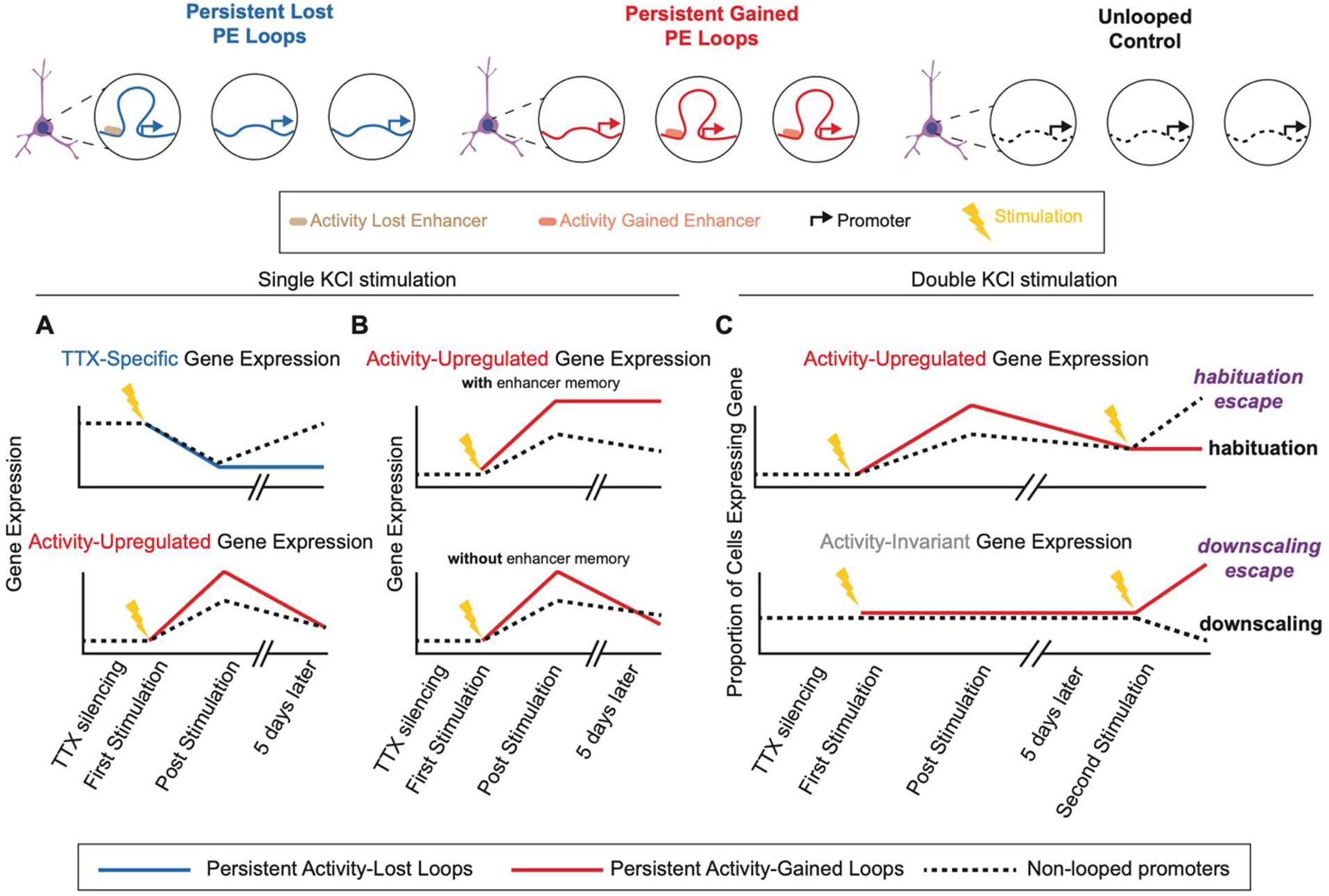
Link among persistent chromatin loops and plasticity in activity-dependent and - independent gene expression during a timecourse of stimulation and re-stimulation of human neurons. **(A)** Persistent activity-lost loops are enriched for activity-downregulated gene expression that remains persistently downregulated 5 days after simulation (top). Persistent activity-gained loops are enriched for activity-upregulated gene expression at the expected time point of 3-6 hours after stimulation, but the expression resumes to levels close to baseline by 5 days post-stimulation (bottom). **(B)** We identified a small subset of persistent activity-gained loops connecting enhancers to promoters of activity-upregulated genes that remain persistently upregulated at 5 days post-stimulation (top). The putative non-coding enhancers are bound by CTCF and exhibit enduring traces of the histone modification H3K27ac. **(C)** We treated human neurons to a second stimulation and measured the proportion of cells with detectable mRNA levels. Persistent activity-gained loops (red) exhibit two types of compensatory gene expression phenomena after repeated pharmacological stimulation. Activity-upregulated genes are re-induced after second stimulation when unlooped. By contrast at persistently gained loops, the activity-upregulated genes exhibit habituation and are not induced during second stimulation. Moreover, activity-independent, invariantly expressed synaptic genes are downregulated only upon second stimulation when unlooped. If anchoring persistently gained loops, such genes unexpectedly escape from downscaling and become activity-upregulated at second stimulation.

Recent studies demonstrate that DNA methyltransferases are required for memory formation and consolidation in murine models (*78, 79*), suggesting that activity-induced changes in DNA methylation might also play an important role in learning and memory. Furthermore, studies using *in vitro* and *in vivo* model systems and bulk DNA methylation assays have demonstrated dynamic changes in DNA methylation upon neural stimulation (*78-84, 95, 96*). However, activity-induced patterns of DNA methylation at regulatory elements anchoring chromatin looping have not yet been explicitly defined in human post-mitotic neurons. Classic foundational studies have demonstrated that regulatory elements bound by CTCF are refractory to DNA methylation at promoters of developmentally regulated and imprinted genes (*85-87*). Here, we find that a near negligible minor fraction of both promoters and enhancers anchoring loops exhibit dynamic changes in DNA methylation upon neural stimulation. Moreover, we find that differentially methylated CpG loci (DMLs) almost exclusively occur at promoters and enhancers devoid of CTCF occupancy. Our observations are consistent with our working model that CTCF protects looped promoters and enhancers from dynamic activity-dependent DNA methylation changes during the stimulation of human neurons.

A general model of memory posits that the repetitive exposure to a common experience or stimulus, such as multiple training events during learning (*48, 97, 98*), continued use of a drug in addiction (*99-101*), or repeated exposure to aversive stimuli leading to trauma (*102, 103*), underlies aspects of the encoding and storage of memory. To explore molecular correlates of the repetitive induction of action potential firing in human neurons, we re-exposed human neurons to a second round of KCl stimulation five days after the first treatment. We uncovered major, unexpected observations. First, activity-inducible genes were robustly restimulated upon second KCl exposure when unlooped. By contrast, activity-inducible genes connected to activity-inducible enhancers in persistently gained PE loops failed to upregulate upon re-stimulation, suggesting a form of chromatin habituation (**Fig. 6C, top**). Second, we identified a large subset of activity-independent, invariantly expressed synaptic genes. Although invariantly expressed on first stimulation, a subset of these genes responded to re-stimulation by further up regulation or severe down regulation when unlooped. By contrast, when anchoring persistently gained PE loops, activity-independent synaptic genes could be activity-stimulated and protected from the phenomena of chromatin downscaling in human neurons (**Fig. 6C, bottom**). Our data reveal a link between the persistence of chromatin structure and the plasticity of gene expression patterns, including chromatin downscaling and habituation, over time and in response to multiple exposures of a stimulus in human neurons.

It is established that organisms can reduce their response to a repetitive stimulus through habituation (*104*) and neuronal circuits can downscale their synaptic connections and strength upon repeated activity (*105*). Here, we observe striking parallels at the biological scale of the nucleus with phenomena of gene expression habituation and downscaling in human post-mitotic neurons. Our data are consistent with a model in which persistently gained loops protect genes from extreme expression up or downregulation upon second stimulation. Overall, our work reports the unexpected phenomena of persistent structural traces in the form of activity dependent higher-order chromatin loops and suggest that loop persistence shapes how the mammalian neural genome remodels, remembers and responds over time and across multiple stimulations. Future work integrating phenomena of chromatin and gene expression persistence with synapse and circuit physiology *in vivo* will shed important new light into the molecular and cellular mechanisms governing long-term memory storage.

## Acknowledgements

We thank Tim Pollex, Rahul Sureka, Harshini Chandrashekar, and all members of the Cremins lab for helpful discussions. We thank Rahul Sureka and Gui Hu for assistance with sequencing. K. Pham is the recipient of a F30 Training Grant. K. Titus is the recipient of a Fontaine fellowship. A. Nikish is the recipient of an NIH T32 Training grant. H. Ryu is the recipient of the Michael S. Brown Graduate Research Fellowship. R. Patel is the recipient of an F31 Training Grant. J. E. Phillips-Cremins was a New York Stem Cell Foundation Robertson Investigator and an Alred P. Sloan Foundation Fellow during this project. We thank the UPenn Cell and Developmental Biology Core (RRID SCR_022373) and the Children’s Hospital of Philadelphia Flow Cytometry Core Laboratory (RRID: SCR_009726) for providing instrumentation. We acknowledge the NIH S10 grant funding mechanism (J.E.P-C) for supporting the MEA equipment used to generate neurophysiology measurements in this manuscript.

## Funding

New York Stem Cell Foundation Robertson Investigator (NYSCF-R-124; J.E.P-C)

NIH NIMH (2R01MH120269; J.E.P-C)

NIH NINDS (1R01NS114226; J.E.P-C)

NSF CAREER Award (CBE1943945; J.E.P-C)

4D Nucleome Common Fund (1U01DK127405; J.E.P-C)

4D Nucleome Common Fund (1U01DA052715; J.E.P-C)

NIH Common Fund / NIMH (1DP1MH129957; J.E.P-C)

PA Department of Health – CURE (2020F07; J.E.P-C)

CZI Neurodegenerative Disease Pairs Award (DAF2022-250430; J.E.P-C)

CZI Neurodegenerative Disease Pairs Award (DAF2022-250577; J.E.P-C)

NIH Shared Instrumentation Grant (1S10OD032363; J.E.P-C)

Cure Huntington’s Disease Initiative (CHDI; ID A-18744, to J.E.P-C)

NSF Emerging Frontiers Research Innovation (EFMA19-33400; JEPC)

NIH F30 (F30HD114405; K.P.)

NIH F31 (F31HG014082; R.P.)

## Author contributions

Conceptualization: AJW, AN, JEPC

Formal Analysis: AJW, KP, KRT, CL, KRC, PX, RP

Methodology: AJW, KP, KRT, CL, KRC, HSR, RB

Visualization: AJW, KP, CL

Investigation: AJW, AN, KP, HSR, SM

Funding acquisition: JEPC

Project administration: JEPC

Writing: AJW, KP, TP, JEPC

## Data and materials availability

All data is provided at GEO under the accessions: GSEXXXA (snm3c), GSEXXXB (CUT&Tag), GSEXXXC (RNA-seq), GSEXXXD (ChIP-seq) and GSEXXXE (snRNA-seq), and all data and code will be available upon publication.

## Supplementary Materials

Materials and Methods

Supplementary Text

Figs. S1 to S21

Tables S1 to S13

## References and Notes

1. T. J. Jarome, F. D. Lubin, Epigenetic mechanisms of memory formation and reconsolidation. Neurobiol Learn Mem 115, 116–127 (2014).

2. M. Kyrke-Smith, J. M. Williams, Bridging Synaptic and Epigenetic Maintenance Mechanisms of the Engram. Front Mol Neurosci 11, 369 (2018).

3. W. C. Abraham, O. D. Jones, D. L. Glanzman, Is plasticity of synapses the mechanism of long-term memory storage? NPJ Sci Learn 4, 9 (2019).

4. T. Takeuchi, A. J. Duszkiewicz, R. G. Morris, The synaptic plasticity and memory hypothesis: encoding, storage and persistence. Philos Trans R Soc Lond B Biol Sci 369, 20130288 (2014).

5. J. A. Beagan et al., Three-dimensional genome restructuring across timescales of activity-induced neuronal gene expression. Nat Neurosci 23, 707–717 (2020).

6. E.-L. Yap, M. E. Greenberg, Activity-Regulated Transcription: Bridging the Gap between Neural Activity and Behavior. Neuron 100, 330–348 (2018).

7. T. H. Ch’ng, K. C. Martin, Synapse-to-nucleus signaling. Curr Opin Neurobiol 21, 345–352 (2011).

8. R. L. Redondo, R. G. Morris, Making memories last: the synaptic tagging and capture hypothesis. Nat Rev Neurosci 12, 17–30 (2011).

9. H. Kang, E. M. Schuman, A requirement for local protein synthesis in neurotrophin-induced hippocampal synaptic plasticity. Science 273, 1402–1406 (1996).

10. E. M. Schuman, Neuroscience: Local protein sources drive memory. Curr Biol 32, R786–R788 (2022).

11. C. Sun et al., The prevalence and specificity of local protein synthesis during neuronal synaptic plasticity. Sci Adv 7, eabj0790 (2021).

12. K. C. Martin et al., Synapse-specific, long-term facilitation of aplysia sensory to motor synapses: a function for local protein synthesis in memory storage. Cell 91, 927–938 (1997).

13. A. Casadio et al., A transient, neuron-wide form of CREB-mediated long-term facilitation can be stabilized at specific synapses by local protein synthesis. Cell 99, 221–237 (1999).

14. J. A. Beagan et al., Local Genome Topology Can Exhibit an Incompletely Rewired 3D-Folding State during Somatic Cell Reprogramming. Cell Stem Cell 18, 611–624 (2016).

15. S. C. Hsu et al., The BET Protein BRD2 Cooperates with CTCF to Enforce Transcriptional and Architectural Boundaries. Mol Cell 66, 102–116 e107 (2017).

16. J. H. Kim et al., 5C-ID: Increased resolution Chromosome-Conformation-Capture-Carbon-Copy with in situ 3C and double alternating primer design. Methods 142, 39–46 (2018).

17. H. K. Norton et al., Detecting hierarchical genome folding with network modularity. Nat Methods 15, 119–122 (2018).

18. T. G. Gilgenast, J. E. Phillips-Cremins, Systematic Evaluation of Statistical Methods for Identifying Looping Interactions in 5C Data. Cell Syst 8, 197–211 e113 (2019).

19. H. Zhang et al., Chromatin structure dynamics during the mitosis-to-G1 phase transition. Nature 576, 158–162 (2019).

20. D. Zhang et al., Alteration of genome folding via contact domain boundary insertion. Nat Genet 52, 1076–1087 (2020).

21. J. E. Phillips-Cremins et al., Architectural protein subclasses shape 3D organization of genomes during lineage commitment. Cell 153, 1281–1295 (2013).

22. J. A. Beagan et al., YY1 and CTCF orchestrate a 3D chromatin looping switch during early neural lineage commitment. Genome Res 27, 1139–1152 (2017).

23. J. H. Sun et al., Disease-Associated Short Tandem Repeats Co-localize with Chromatin Domain Boundaries. Cell 175, 224–238 e215 (2018).

24. J. H. Kim et al., LADL: light-activated dynamic looping for endogenous gene expression control. Nat Methods 16, 633–639 (2019).

25. L. R. Fernandez, T. G. Gilgenast, J. E. Phillips-Cremins, 3DeFDR: statistical methods for identifying cell type-specific looping interactions in 5C and Hi-C data. Genome Biology 21, 219 (2020).

26. T. Malachowski et al., Spatially coordinated heterochromatinization of long synaptic genes in fragile X syndrome. Cell 186, 5840–5858.e5836 (2023).

27. L. Calderon et al., Cohesin-dependence of neuronal gene expression relates to chromatin loop length. Elife 11, e76539 (2022).

28. D. J. Emerson et al., Cohesin-mediated loop anchors confine the locations of human replication origins. Nature, 1–8 (2022).

29. S. S. Rao et al., A 3D map of the human genome at kilobase resolution reveals principles of chromatin looping. Cell 159, 1665–1680 (2014).

30. A. S. Weintraub et al., YY1 Is a Structural Regulator of Enhancer-Promoter Loops. Cell 171, 1573–1588 e1528 (2017).

31. S. S. P. Rao et al., Cohesin Loss Eliminates All Loop Domains. Cell 171, 305–320 e324 (2017).

32. W. Schwarzer et al., Two independent modes of chromatin organization revealed by cohesin removal. Nature 551, 51–56 (2017).

33. D. H. Phanstiel et al., Static and Dynamic DNA Loops form AP-1-Bound Activation Hubs during Macrophage Development. Mol Cell 67, 1037–1048 e1036 (2017).

34. D. Hnisz et al., Activation of proto-oncogenes by disruption of chromosome neighborhoods. Science 351, 1454–1458 (2016).

35. V. Narendra et al., Transcription. CTCF establishes discrete functional chromatin domains at the Hox clusters during differentiation. Science 347, 1017–1021 (2015).

36. M. Franke et al., Formation of new chromatin domains determines pathogenicity of genomic duplications. Nature 538, 265–269 (2016).

37. D. G. Lupianez et al., Disruptions of topological chromatin domains cause pathogenic rewiring of gene-enhancer interactions. Cell 161, 1012–1025 (2015).

38. A. Sanyal, B. R. Lajoie, G. Jain, J. Dekker, The long-range interaction landscape of gene promoters. Nature 489, 109–113 (2012).

39. E. M. Smith, B. R. Lajoie, G. Jain, J. Dekker, Invariant TAD Boundaries Constrain Cell-Type-Specific Looping Interactions between Promoters and Distal Elements around the CFTR Locus. Am J Hum Genet 98, 185–201 (2016).

40. B. M. Javierre et al., Lineage-Specific Genome Architecture Links Enhancers and Non-coding Disease Variants to Target Gene Promoters. Cell 167, 1369–1384 e1319 (2016).

41. O. Joshi et al., Dynamic Reorganization of Extremely Long-Range Promoter-Promoter Interactions between Two States of Pluripotency. Cell Stem Cell 17, 748–757 (2015).

42. T. Yamada et al., Sensory experience remodels genome architecture in neural circuit to drive motor learning. Nature 569, 708–713 (2019).

43. A. Marco et al., Mapping the epigenomic and transcriptomic interplay during memory formation and recall in the hippocampal engram ensemble. Nat Neurosci 23, 1606–1617 (2020).

44. D. S. Sams et al., Neuronal CTCF Is Necessary for Basal and Experience-Dependent Gene Regulation, Memory Formation, and Genomic Structure of BDNF and Arc. Cell Rep 17, 2418–2430 (2016).

45. S. Kim et al., Remote Memory and Cortical Synaptic Plasticity Require Neuronal CCCTC-Binding Factor (CTCF). J Neurosci 38, 5042–5052 (2018).

46. G. Santoni et al., Chromatin plasticity predetermines neuronal eligibility for memory trace formation. Science 385, eadg9982 (2024).

47. S. Ramon y Cajal, The Croonian lecture.—La fine structure des centres nerveux. Proceedings of the Royal Society of London 55, 444–468 (1894).

48. E. R. Kandel, The Molecular Biology of Memory Storage: A Dialogue Between Genes and Synapses. Science, (2001).

49. R. A. Nicoll, A Brief History of Long-Term Potentiation. Neuron 93, 281–290 (2017).

50. E. F. Fornasiero et al., Precisely measured protein lifetimes in the mouse brain reveal differences across tissues and subcellular fractions. Nature Communications 9, 4230 (2018).

51. B. Schwanhäusser et al., Global quantification of mammalian gene expression control. Nature 473, 337–342 (2011).

52. F. Crick, Neurobiology: Memory and molecular turnover. Nature 312, 101–101 (1984).

53. J. E. Lisman, A mechanism for memory storage insensitive to molecular turnover: a bistable autophosphorylating kinase. Proceedings of the National Academy of Sciences 82, 3055–3057 (1985).

54. E. D. Roberson, J. D. Sweatt, A Biochemical Blueprint for Long-Term Memory. Learn. Mem. 6, 381–388 (1999).

55. J.-H. Tao-Cheng, Stimulation-induced structural changes at the nucleus, endoplasmic reticulum and mitochondria of hippocampal neurons. Molecular Brain 11, 44 (2018).

56. C. Sanchez-Priego et al., Mapping cis-regulatory elements in human neurons links psychiatric disease heritability and activity-regulated transcriptional programs. Cell Reports 39, 110877 (2022).

57. Y. Zhang et al., Rapid single-step induction of functional neurons from human pluripotent stem cells. Neuron 78, 785–798 (2013).

58. C. Wang et al., Scalable Production of iPSC-Derived Human Neurons to Identify Tau-Lowering Compounds by High-Content Screening. Stem Cell Reports 9, 1221–1233 (2017).

59. R. E. Dolmetsch, U. Pajvani, K. Fife, J. M. Spotts, M. E. Greenberg, Signaling to the nucleus by an L-type calcium channel-calmodulin complex through the MAP kinase pathway. Science 294, 333–339 (2001).

60. K. M. Tyssowski et al., Different Neuronal Activity Patterns Induce Different Gene Expression Programs. Neuron 98, 530–546.e511 (2018).

61. K. D. A. Rienecker et al., Mild membrane depolarization in neurons induces immediate early gene transcription and acutely subdues responses to a successive stimulus. J Biol Chem 298, 102278 (2022).

62. K. D. A. Rienecker, R. G. Poston, R. N. Saha, Merits and Limitations of Studying Neuronal Depolarization-Dependent Processes Using Elevated External Potassium. ASN Neuro 12, 1759091420974807 (2020).

63. X. Shan et al., Fully defined NGN2 neuron protocol reveals diverse signatures of neuronal maturation. Cell Reports Methods 4, (2024).

64. D.-S. Lee et al., Simultaneous profiling of 3D genome structure and DNA methylation in single human cells. Nat Methods 16, 999–1006 (2019).

65. H. Liu et al., Single-cell DNA methylome and 3D multi-omic atlas of the adult mouse brain. Nature 624, 366–377 (2023).

66. M. G. Heffel et al., Temporally distinct 3D multi-omic dynamics in the developing human brain. Nature 635, 481–489 (2024).

67. M. J. Solomon, P. L. Larsen, A. Varshavsky, Mapping protein-DNA interactions in vivo with formaldehyde: evidence that histone H4 is retained on a highly transcribed gene. Cell 53, 937–947 (1988).

68. 68. S. Henikoff, J. Henikoff, K. Ahmad, CUT&Tag-direct for whole cells with CUTAC. (2023).

69. S. Henikoff, J. G. Henikoff, H. S. Kaya-Okur, K. Ahmad, Efficient chromatin accessibility mapping in situ by nucleosome-tethered tagmentation. Elife 9, e63274 (2020).

70. K. R. Titus, Z. Simandi, H. Chandrashekar, D. Paquet, J. E. Phillips-Cremins, Cell-type-specific loops linked to RNA polymerase II elongation in human neural differentiation. Cell Genom 4, 100606 (2024).

71. J. E. Phillips-Cremins, V. G. Corces, Chromatin insulators: linking genome organization to cellular function. Mol Cell 50, 461–474 (2013).

72. H. Chandrashekar et al., A multi-looping chromatin signature predicts dysregulated gene expression in neurons with familial Alzheimer’s disease mutations. bioRxiv, (2024).

73. S. Zhang et al., The protective effect and potential mechanism of NRXN1 on learning and memory in ADHD rat models. Exp Neurol 344, 113806 (2021).

74. J. Feng et al., High frequency of neurexin 1beta signal peptide structural variants in patients with autism. Neurosci Lett 409, 10–13 (2006).

75. B. M. Schwenk et al., The FTLD risk factor TMEM106B and MAP6 control dendritic trafficking of lysosomes. EMBO J 33, 450–467 (2014).

76. K. H. Zent, M. L. Dell’Acqua, Synapse-to-nucleus ERK-CREB transcriptional signaling requires dendrite-to-soma Ca(2+)propagation mediated by L-type voltage-gated Ca(2+) channels. J Neurosci, (2024).

77. H. Bito, K. Deisseroth, R. W. Tsien, CREB phosphorylation and dephosphorylation: a Ca(2+)- and stimulus duration-dependent switch for hippocampal gene expression. Cell 87, 1203–1214 (1996).

78. C. A. Miller, J. D. Sweatt, Covalent Modification of DNA Regulates Memory Formation. Neuron 53, 857–869 (2007).

79. J. J. Day et al., DNA methylation regulates associative reward learning. Nat Neurosci 16, 1445–1452 (2013).

80. J. U. Guo et al., Neuronal activity modifies the DNA methylation landscape in the adult brain. Nat Neurosci 14, 1345–1351 (2011).

81. E. Hannon et al., A role for CaV1 and calcineurin signaling in depolarization-induced changes in neuronal DNA methylation. Neuroepigenetics 3, 1–6 (2015).

82. C. A. Miller et al., Cortical DNA methylation maintains remote memory. Nat Neurosci 13, 664–666 (2010).

83. R. Halder et al., DNA methylation changes in plasticity genes accompany the formation and maintenance of memory. Nat Neurosci 19, 102–110 (2016).

84. J. J. Day, J. D. Sweatt, DNA methylation and memory formation. Nat Neurosci 13, 1319–1323 (2010).

85. A. C. Bell, G. Felsenfeld, Methylation of a CTCF-dependent boundary controls imprinted expression of the Igf2 gene. Nature 405, 482–485 (2000).

86. J. E. Phillips, V. G. Corces, CTCF: Master Weaver of the Genome. Cell 137, 1194–1211 (2009).

87. A. Feldmann et al., Transcription factor occupancy can mediate active turnover of DNA methylation at regulatory regions. PLoS Genet 9, e1003994 (2013).

88. S. Cuartero et al., Control of inducible gene expression links cohesin to hematopoietic progenitor self-renewal and differentiation. Nat Immunol 19, 932–941 (2018).

89. B. D. Strahl, C. D. Allis, The language of covalent histone modifications. Nature 403, 41–45 (2000).

90. J. P. Lopez-Atalaya, A. Barco, Can changes in histone acetylation contribute to memory formation? Trends Genet 30, 529–539 (2014).

91. J. M. Alarcon et al., Chromatin acetylation, memory, and LTP are impaired in CBP+/- mice: a model for the cognitive deficit in Rubinstein-Taybi syndrome and its amelioration. Neuron 42, 947–959 (2004).

92. E. Korzus, M. G. Rosenfeld, M. Mayford, CBP histone acetyltransferase activity is a critical component of memory consolidation. Neuron 42, 961–972 (2004).

93. J. M. Levenson et al., Regulation of histone acetylation during memory formation in the hippocampus. J Biol Chem 279, 40545–40559 (2004).

94. L. Peixoto, T. Abel, The role of histone acetylation in memory formation and cognitive impairments. Neuropsychopharmacol. 38, 62–76 (2013).

95. D. K. Ma et al., Neuronal Activity–Induced Gadd45b Promotes Epigenetic DNA Demethylation and Adult Neurogenesis. Science 323, 1074–1077 (2009).

96. T. Kameda et al., Neuronal activation modulates enhancer activity of genes for excitatory synaptogenesis through de novo DNA methylation. J Reprod Dev 67, 369–379 (2021).

97. V. M. Ho, J. A. Lee, K. C. Martin, The cell biology of synaptic plasticity. Science 334, 623–628 (2011).

98. M. Fu, X. Yu, J. Lu, Y. Zuo, Repetitive motor learning induces coordinated formation of clustered dendritic spines in vivo. Nature 483, 92–95 (2012).

99. S. E. Hyman, Addiction: a disease of learning and memory. Am J Psychiatry 162, 1414–1422 (2005).

100. E. J. Nestler, Cellular basis of memory for addiction. Dialogues Clin Neurosci 15, 431–443 (2013).

101. S. E. Hyman, R. C. Malenka, E. J. Nestler, Neural mechanisms of addiction: the role of reward-related learning and memory. Annu Rev Neurosci 29, 565–598 (2006).

102. J. J. Kim, M. S. Fanselow, Modality-specific retrograde amnesia of fear. Science 256, 675–677 (1992).

103. S. Lissek, B. van Meurs, Learning models of PTSD: Theoretical accounts and psychobiological evidence. Int J Psychophysiol 98, 594–605 (2015).

104. V. Castellucci, H. Pinsker, I. Kupfermann, E. R. Kandel, Neuronal mechanisms of habituation and dishabituation of the gill-withdrawal reflex in Aplysia. Science 167, 1745–1748 (1970).

105. G. G. Turrigiano, The self-tuning neuron: synaptic scaling of excitatory synapses. Cell 135, 422–435 (2008).

106. W. Xie et al., Neuronal sequences in population bursts encode information in human cortex. Nature, 1–8 (2024).

107. H. Bading, D. D. Ginty, M. E. Greenberg, Regulation of gene expression in hippocampal neurons by distinct calcium signaling pathways. Science 260, 181–186 (1993).

108. P. Roussos, B. Guennewig, D. C. Kaczorowski, G. Barry, K. J. Brennand, Activity-Dependent Changes in Gene Expression in Schizophrenia Human-Induced Pluripotent Stem Cell Neurons. JAMA Psychiatry 73, 1180–1188 (2016).

109. M. Bitar, S. Kuiper, E. O’Brien, G. Barry, Using Human iPSC-Derived Neurons to Uncover Activity-Dependent Non-Coding RNAs. Genes (Basel) 8, E401 (2017).

110. V. Hook et al., Human iPSC Neurons Display Activity-Dependent Neurotransmitter Secretion: Aberrant Catecholamine Levels in Schizophrenia Neurons. Stem Cell Reports 3, 531–538 (2014).

111. I. L. Ibarra et al., Comparative chromatin accessibility upon BDNF stimulation delineates neuronal regulatory elements. Mol Syst Biol 18, e10473 (2022).

112. A. Meissner et al., Genome-scale DNA methylation maps of pluripotent and differentiated cells. Nature 454, 766–770 (2008).

113. D. S. Park et al., High-throughput Oligopaint screen identifies druggable 3D genome regulators. Nature 620, 209–217 (2023).

114. H. Huang et al., A subset of topologically associating domains fold into mesoscale core-periphery networks. Sci Rep 9, 9526 (2019).

115. K. Girdhar et al., Chromatin domain alterations linked to 3D genome organization in a large cohort of schizophrenia and bipolar disorder brains. Nat Neurosci 25, 474–483 (2022).

116. D. Y. Kwon et al., Neuronal Yin Yang1 in the prefrontal cortex regulates transcriptional and behavioral responses to chronic stress in mice. Nat Commun 13, 55 (2022).

117. E. A. Maury et al., Schizophrenia-associated somatic copy-number variants from 12,834 cases reveal recurrent NRXN1 and ABCB11 disruptions. Cell Genom 3, 100356 (2023).

118. M. H. Guo, W. P. Lee, B. Vardarajan, G. D. Schellenberg, J. E. Phillips-Cremins, Polygenic burden of short tandem repeat expansions promotes risk for Alzheimer’s disease. Nat Commun 16, 1126 (2025).

119. R. Patel, K. Pham, H. Chandrashekar, J. E. Phillips-Cremins, FISHnet: detecting chromatin domains in single-cell sequential Oligopaints imaging data. Nat Methods 22, 1255–1264 (2025).

120. K. Pham et al., Mosaic H3K9me3 at BREACHes predicts synaptic gene expression associated with fragile X syndrome cognitive severity. bioRxiv, (2025).

121. J. M. Goldstein, L. C. Litwin, NBQX is a selective non-NMDA receptor antagonist in rat hippocampal slice. Mol Chem Neuropathol 18, 145–152 (1993).

122. D. B. Wheeler, A. Randall, R. W. Tsien, Roles of N-Type and Q-Type Ca2+ Channels in Supporting Hippocampal Synaptic Transmission. Science 264, 107–111 (1994).

123. C. Stringer, T. Wang, M. Michaelos, M. Pachitariu, Cellpose: a generalist algorithm for cellular segmentation. Nat Methods 18, 100–106 (2021).

124. A. Nott, J. C. M. Schlachetzki, B. R. Fixsen, C. K. Glass, Nuclei isolation of multiple brain cell types for omics interrogation. Nat Protoc 16, 1629–1646 (2021).

125. D. L. Lafontaine, L. Yang, J. Dekker, J. H. Gibcus, Hi-C 3.0: Improved Protocol for Genome-Wide Chromosome Conformation Capture. Current Protocols 1, e198 (2021).

126. P. Xu et al., Learning induces persistent chromatin loops underlying robust gene expression during memory recall. bioRxiv, (2025).

127. D. Kim, J. M. Paggi, C. Park, C. Bennett, S. L. Salzberg, Graph-based genome alignment and genotyping with HISAT2 and HISAT-genotype. Nat Biotechnol 37, 907–915 (2019).

128. H. Li et al., The Sequence Alignment/Map format and SAMtools. Bioinformatics 25, 2078–2079 (2009).

129. S. Kovaka et al., Transcriptome assembly from long-read RNA-seq alignments with StringTie2. Genome Biol 20, 278 (2019).

130. M. I. Love, W. Huber, S. Anders, Moderated estimation of fold change and dispersion for RNA-seq data with DESeq2. Genome Biology 15, 550 (2014).

131. M. Martin, Cutadapt removes adapter sequences from high-throughput sequencing reads. EMBnet.journal 17, 10–12 (2011).

132. B. Langmead, S. L. Salzberg, Fast gapped-read alignment with Bowtie 2. Nat Methods 9, 357–359 (2012).

133. A. R. Quinlan, I. M. Hall, BEDTools: a flexible suite of utilities for comparing genomic features. Bioinformatics 26, 841–842 (2010).

134. Y. Zhang et al., Model-based Analysis of ChIP-Seq (MACS). Genome Biology 9, R137 (2008).

135. F. Ramirez et al., deepTools2: a next generation web server for deep-sequencing data analysis. Nucleic Acids Res 44, W160–165 (2016).

136. B. Langmead, C. Trapnell, M. Pop, S. L. Salzberg, Ultrafast and memory-efficient alignment of short DNA sequences to the human genome. Genome Biol 10, R25 (2009).

137. S. Marco-Sola, M. Sammeth, R. Guigo, P. Ribeca, The GEM mapper: fast, accurate and versatile alignment by filtration. Nat Methods 9, 1185–1188 (2012).

138. J. Zhou et al., Robust single-cell Hi-C clustering by convolution- and random-walk-based imputation. Proc Natl Acad Sci U S A 116, 14011–14018 (2019).

139. B. S. Pederson. (2014).

140. H. Liu et al., DNA methylation atlas of the mouse brain at single-cell resolution. Nature 598, 120–128 (2021).

141. T. S. Mikkelsen et al., Genome-wide maps of chromatin state in pluripotent and lineage-committed cells. Nature 448, 553–560 (2007).

142. S. Saxonov, P. Berg, D. L. Brutlag, A genome-wide analysis of CpG dinucleotides in the human genome distinguishes two distinct classes of promoters. Proc Natl Acad Sci U S A 103, 1412–1417 (2006).

143. H. Feng, K. N. Conneely, H. Wu, A Bayesian hierarchical model to detect differentially methylated loci from single nucleotide resolution sequencing data. Nucleic Acids Res 42, e69 (2014).

144. H. Wu et al., Detection of differentially methylated regions from whole-genome bisulfite sequencing data without replicates. Nucleic Acids Res 43, e141 (2015).

145. Y. Park, H. Wu, Differential methylation analysis for BS-seq data under general experimental design. Bioinformatics 32, 1446–1453 (2016).

146. J. M. Elizarraras et al., WebGestalt 2024: faster gene set analysis and new support for metabolomics and multi-omics. Nucleic Acids Res 52, W415–W421 (2024).

147. G. X. Zheng et al., Massively parallel digital transcriptional profiling of single cells. Nat Commun 8, 14049 (2017).

148. Y. Hao et al., Dictionary learning for integrative, multimodal and scalable single-cell analysis. Nat Biotechnol 42, 293–304 (2024).

