## Supplementary Material for "Persistent chromatin loops shape gene expression plasticity upon stimulation and restimulation of human neurons"

Science
AAAS

Supplementary Materials for

### Persistent chromatin loops shape gene expression plasticity upon stimulation and restimulation of human neurons

Abraham J. Waldman[1,2,3,†], Kenneth Pham[1,2,3,†], Katelyn R. Titus[1,2,3], Alexandria Nikish[1,2,3], Constin Liu[1,2,3], Han-Seul Ryu[1,2,3], Srikar S. Muppidi[2,3,4], Keerthivasan Raanin Chandradoss[1,2,3], Peibo Xu[1,2,3], Rohan Patel[1,2,3], Ravi Boya[1,2,3], Jennifer E. Phillips-Cremins[1,2,3,*]

**This PDF file includes:**

Materials and Methods
Supplementary Text
Figs S1 to S21
Tables S1-S13

#### Supplementary Text

Recent studies have suggested that population bursting activity in the human brain plays a role in encoding information upon neural activation (*106*). To assess how an external stimulus drives changes in population bursting in DIV35 human iPSC-derived neurons, we added a low concentration of KCl that has been previously shown to drive changes in neuronal firing (*61, 62*). Upon the addition of 10 mM KCl, the number of bursts increased 2.5-fold, from 1,006 to 2,540, in a representative well across a 5-minute recording (**Fig. S2D)**, and the number of network bursts increased more than 2-fold, from 86 to 210 bursts (**Table S3).** While the bursting and network burst frequency more than doubled, the overall number of spikes only marginally increased, as the number of spikes per network burst decreased from 1,049 to 437 Hz (**Table S3, Fig. S2E)**. The average burst duration also halved, decreasing from 0.79 seconds to 0.44 seconds **(Table S3)**. Additionally, KCl addition drove an increase in sporadic bursting, increasing the network inter burst interval coefficient of variation from 0.05 to 0.29 (**Table S3).** KCl driven increases in variation subsequently decreased the overall synchrony of firing, as the area under the normalized cross-correlogram decreased from 0.34 to 0.25 (**Table S3, Fig. S2F**). Our data demonstrate that DIV35 iPSC-derived neurons spontaneously fire action potentials in a manner that is dependent on the KCl concentration.

Stimulation of *in vitro* cortical neurons drives multiple waves of gene expression, beginning with the expression of immediate early genes (IEGs) on the timescales of minutes followed by the calcium-dependent expression of secondary response genes (SRGs) on the timescales of hours (*5*). The strongest activity-dependent gene expression response upon KCl addition to *in vitro* neurons has been observed at concentrations of 30-60 mM KCl (*6, 59, 107*). Multiple studies have observed robust upregulation of mRNA levels (*108, 109*) and neurotransmitter release (*110*) upon neuronal activation of hiPSC-derived neurons via treatment with 50 mM KCl. Neural activation with KCl concentrations of 10 mM and below, however, is unable to induce expression of IEGs possibly due to low calcium influx into the cell. Moreover, 10 mM KCl does not drive increased calcium response element levels in neurons (*59, 61, 62*). By contrast, high KCl concentrations drive increases in intracellular calcium required for synapse-to-nucleus signaling via the mitogen-activated protein kinase (MAPK) pathway, leading to persistent cAMP-response element binding protein (CREB) phosphorylation (pCREB) (*59*). Increases in intracellular calcium via L-type calcium channels are essential to relay signals from the synapse to nucleus to drive gene expression (*76*), and the inhibition of L-type calcium channels by dihydropyridines such as nimodipine blocks the persistent pCREB that otherwise occurs during KCl addition (*59*).

To confirm that DIV35 hiPSC-derived neurons respond to high KCl activation via the expected signaling pathways, we pre-silenced neuronal firing with TTX for 24 hours (*5*) to normalize cell state, stimulated neurons for 1 hour with 50 mM KCl in ACSF, and measured changes in the cFOS protein and the phosphorylation of CREB via IF and Western Blot, respectively. To confirm that 50 mM KCl induces robust expression of the cFOS protein, we performed IF staining for cFOS upon the addition of 50 mM KCl in the presence or absence of the pharmacological agent nimodipine (**Supplementary Methods**). We found that high KCl increased cFOS expression in comparison to TTX, and that this expression was blocked by the presence of nimodipine during stimulation **(Fig S3A,B).** We further confirmed that 50 mM KCl induces CREB phosphorylation, and that this phosphorylation can be blocked by the presence of nimodipine during stimulation (**Fig. S3C,D)**. Taken together, these results show that 50 mM KCl drove the expression of cFOS, and that pharmacological inhibition of L-type calcium channels attenuates the pCREB mediated signaling response.

We next confirmed that KCl addition drove the expected expression of IEGs and SRGs using nuclear RNA-seq (*56, 111*). We observed the maximum expression of most IEGs 1 hour after stimulation. This expression decreased but remained above baseline at the 3- and 6-hour marks for most genes. All observed IEGs and SRGs, except for *EGR1,* had repressed gene expression in the presence of nimodipine (**Fig. S3D,E)**.

Overall, the above findings confirm that DIV35 $i^3$Ns exhibit activity-dependent changes in population bursting, cFOS expression, CREB phosphorylation, and transcription through L-type calcium channels **(Fig. S1-S3)**. Having validated our model system as exhibiting physiologically relevant biomarkers of human neuron physiological traces, biochemistry, and gene expression during stimulation of action potential firing, we set out to advance our studies ascertaining the plasticity of higher-order chromatin architecture, DNA methylation, and gene expression in human neural stimulation.

#### Supplementary Figures

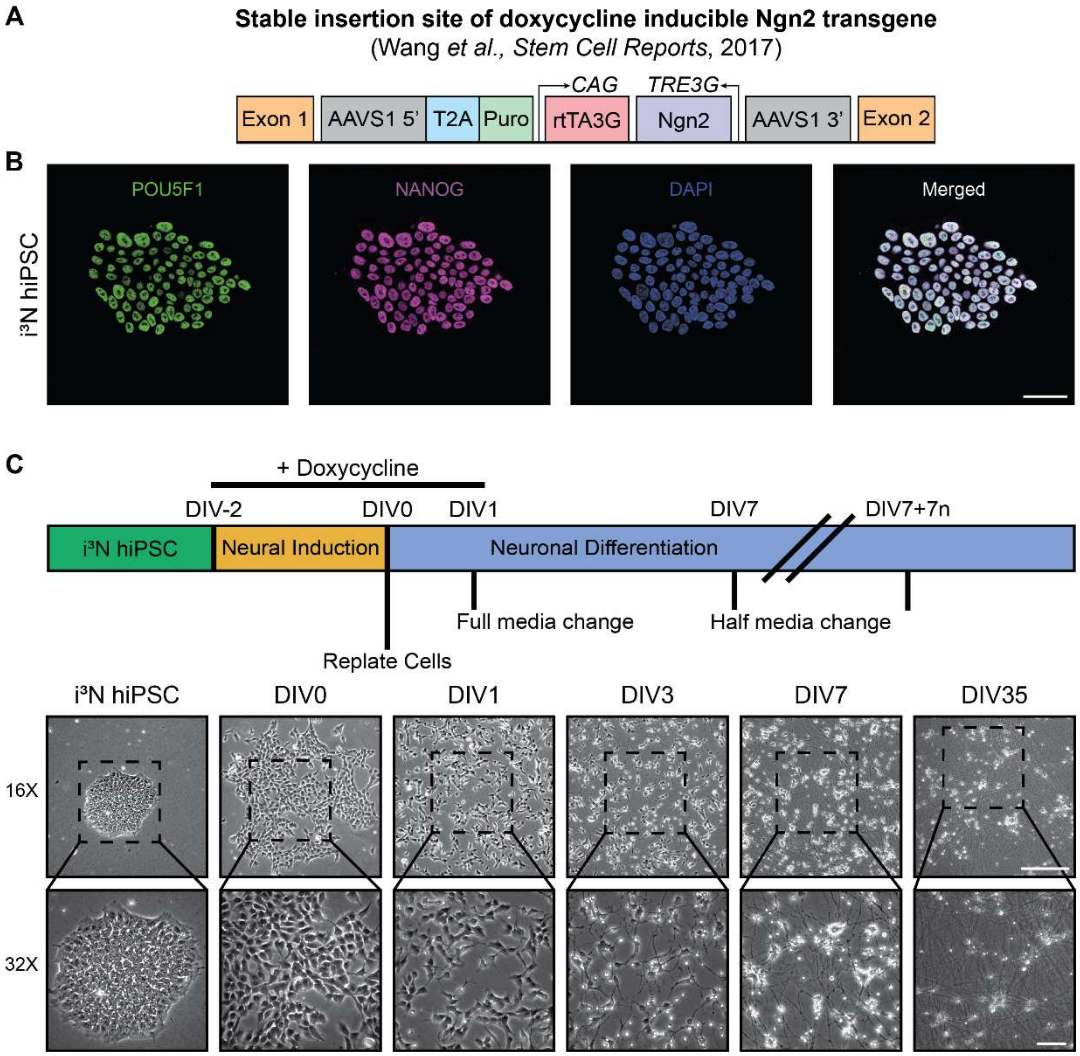


**Fig. S1. Validation of hiPSC state and successful neural induction of i$^3$N model system.** **(A)** Overexpression of Ngn2 via doxycycline induction drives hiPSC differentiation into cortical neurons (*58*). **(B)** Immunofluorescence staining for pluripotency factors POU5F1 and NANOG in i$^3$Ns prior to dox induction; nuclei stained with DAPI. Scale bar = 50 µM. **(C)** Schematic showing media change and neuronal induction schedule with accompanying phase images during differentiation of i$^3$Ns from hiPSCs to neurons. At 30-40% confluency, neuronal differentiation of iPSCs is induced with doxycycline (DIV-2). After two days, neurons are replated onto PLO / laminin coated plates (DIV0). Neurons receive a full media change with doxycycline on DIV1, and a weekly half media change starting on DIV7. Top row scale bar = 250 µm, bottom row scale bar = 75 µm.

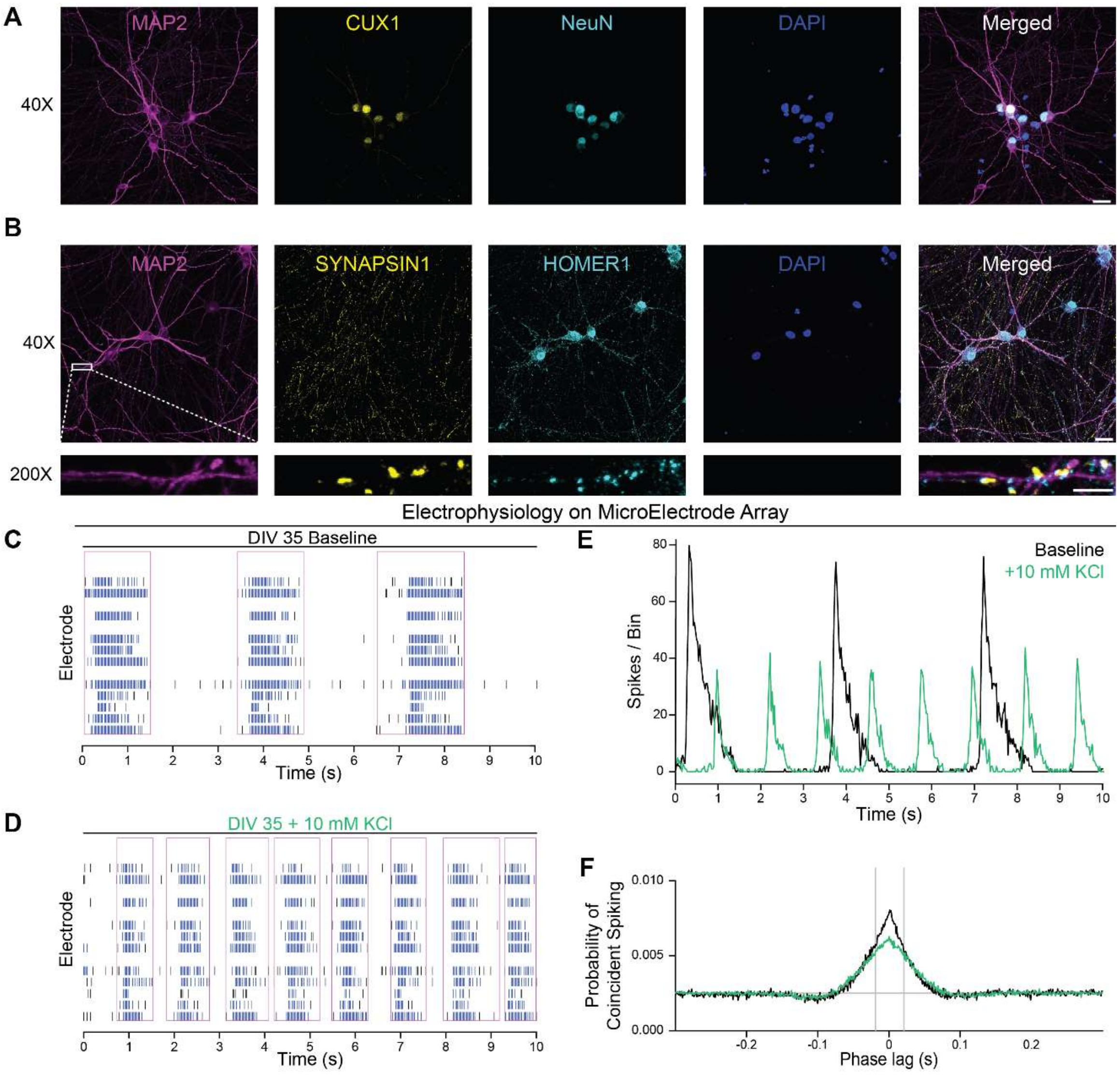


**Fig. S2: Validation of cortical character and development of synaptic connections in DIV35 hiPSC-derived neurons. (A)** Immunofluorescence staining of DIV35 hiPSC-derived i$^3$N neurons express the cortical marker CUX1 and the pan neuronal markers NEUN and MAP2. Scale bar = 25 µm. **(B)** IF of SYNAPSIN1 (yellow) and HOMER1 (cyan) colocalize to MAP2 (pink) in DIV35 hiPSC-derived cortical i$^3$N neurons. 40X scale bar = 25 µm, 200X scale bar = 5 µm. **(C)** MEA raster plot of spontaneous neuronal activity in a representative well of DIV35 hiPSC-derived neurons during a 10 second firing window. Individual spikes are demarcated in black, bursts of spikes in blue, and network bursts in pink. **(D)** MEA raster plot of spontaneous neuronal activity in the same well as **(C)** after the addition of 10 mM KCl. **(E)** **S**pikes in **(C)**, **(D)** binned at 25 ms. **(F)** Normalized cross-correlogram of DIV35 hiPSC-derived neurons before (black) and after (green) the addition of 10 mM KCl. Vertical gray bars represent the 10 ms synchrony window used to calculate synchrony metrics shown in **Table S3.**

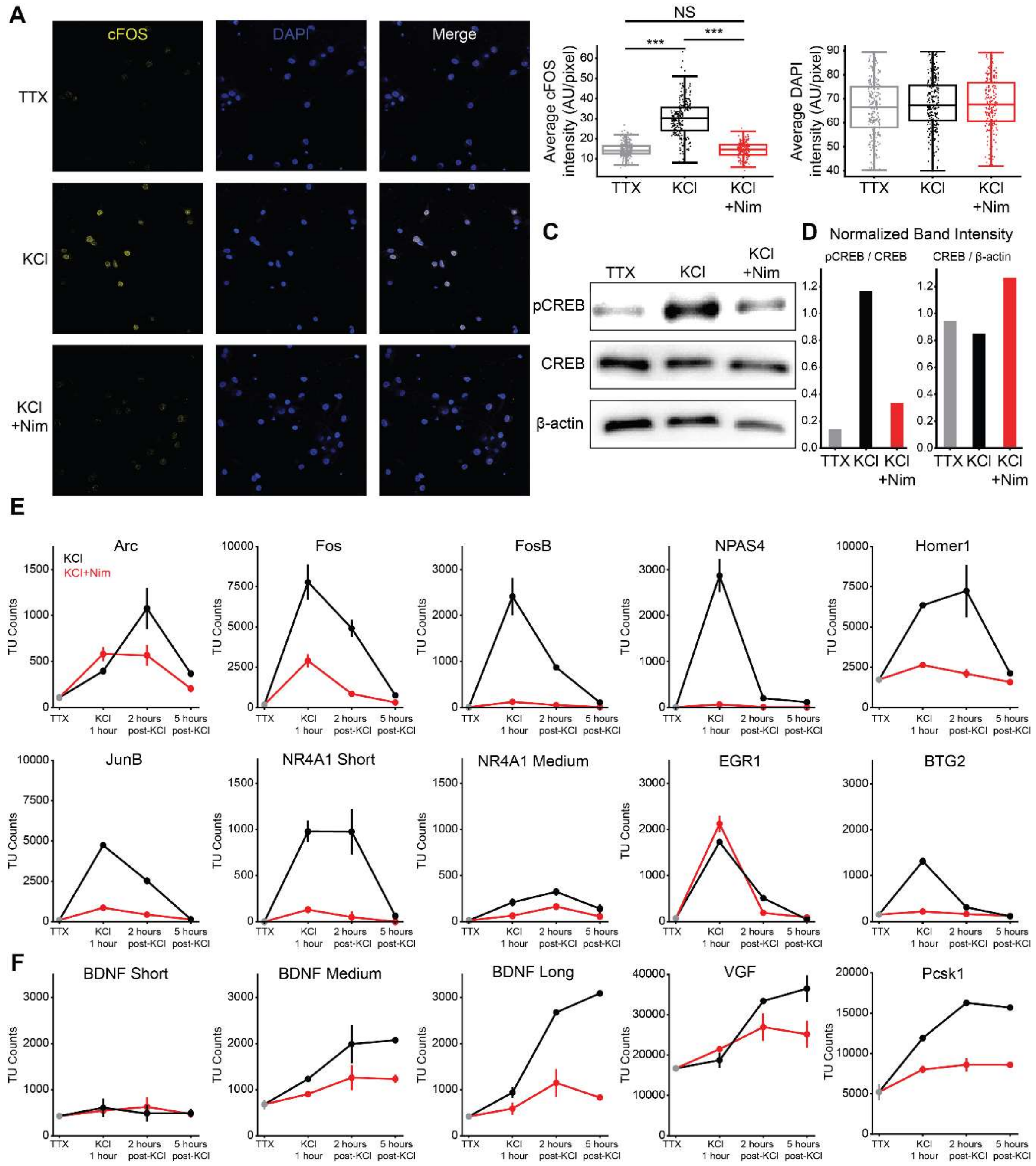


**Fig. S3. 50 mM KCl stimulation drives pCREB dependent expression of IEGs and SRGs**. **(A)** IF staining of cFOS, and DAPI in DIV35 $i^3N$ neurons treated with TTX, TTX followed by 50 mM KCl for 1 hour, and TTX followed by 50 mM KCl + Nim for 1 hour. **(B)** Quantification of cFOS (left) and DAPI (right) intensity per pixel per nucleus for TTX (n=256 nuclei), KCl (n=253 nuclei), and KCl + Nim (n=197 nuclei) conditions. Significance assessed using Welch's t-test; *** indicates significance of $p < .001$. If no significance shown, or NS shown, P value > 0.1. Significance that any DAPI signal is significantly different across conditions tested using ANOVA. **(C)** Western blot of pCREB, CREB, and β-actin signal in DIV 35 neurons treated with TTX, TTX followed by 50 mM

KCl for 1 hour, and TTX followed by 50 mM KCl + Nim for 1 hour. **(D)** Ratio of band intensities for (left) pCREB/CREB and (right) CREB/ β-actin for each condition from Western blot in **(C). (E,F)** Average expression of selected IEGs **(E)** and SRGs **(F)** over time upon the addition of 50 mM KCl in the presence or absence of Nimodipine. Error bars represent standard deviation between two biological replicates.

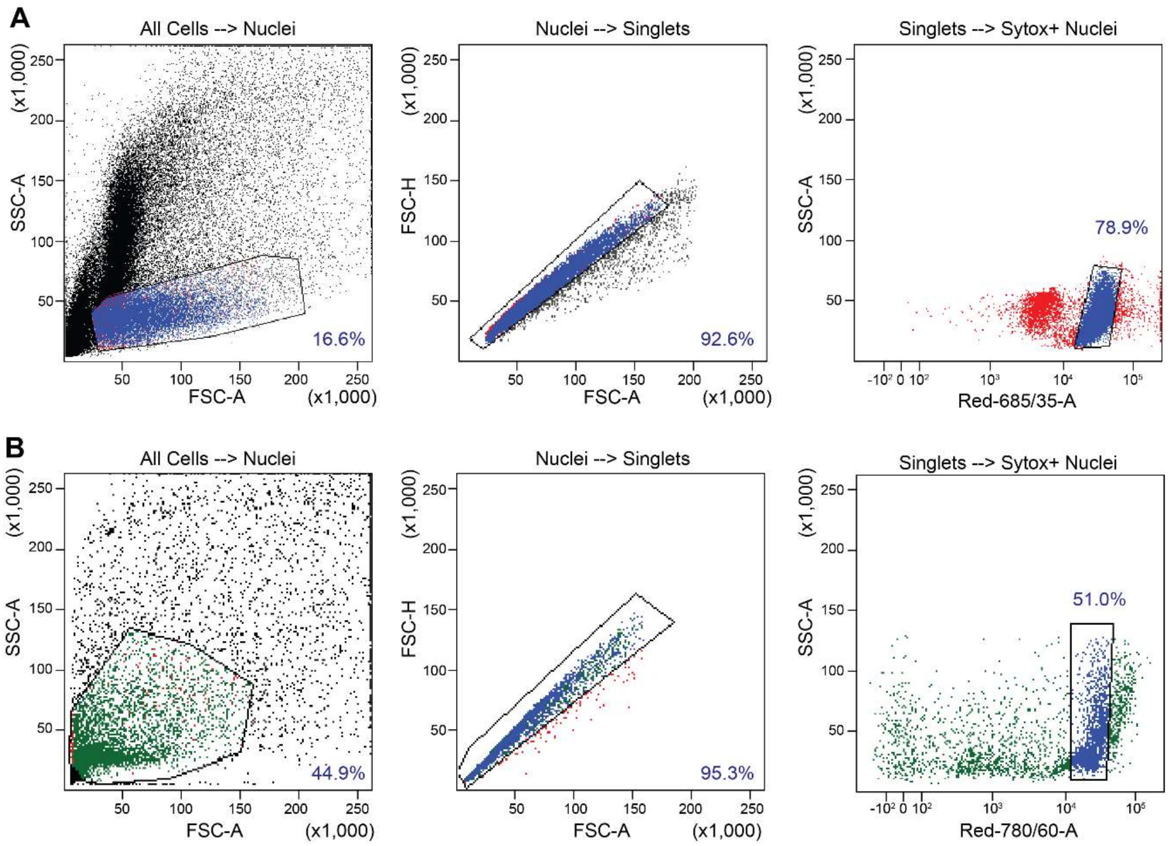


**Fig. S4. Gating strategies used for FANS during snm3c-seq3. (A)** Gating strategy upon nuclei isolation to isolate healthy nuclei for downstream processing. Nuclei are gated on Forward Scatter Area (FSC-A) vs Side Scatter Area (SSC-A) to identify healthy nuclei, on FSC-A vs Forward Scatter Height (FSC-H) to identify singlets, and finally on Sytox Red vs SSC-A to identify Sytox positive, singlet, nuclei. Sorted nuclei were subsequently carried forward for the 3C steps of snm3c (Supplementary Methods). **(B)** Gating strategy for sorting nuclei after proximity ligation step of 3C into single wells of a 384 well plate for downstream bisulfite conversion and single nuclei library preparation. Nuclei are gated on FSC-A vs SSC-A to identify intact nuclei, on FSC-A vs FSC-H to identify singlets and remove any doublets, and finally on Sytox red vs SSC-A to identify Sytox positive, intact singlet nuclei.

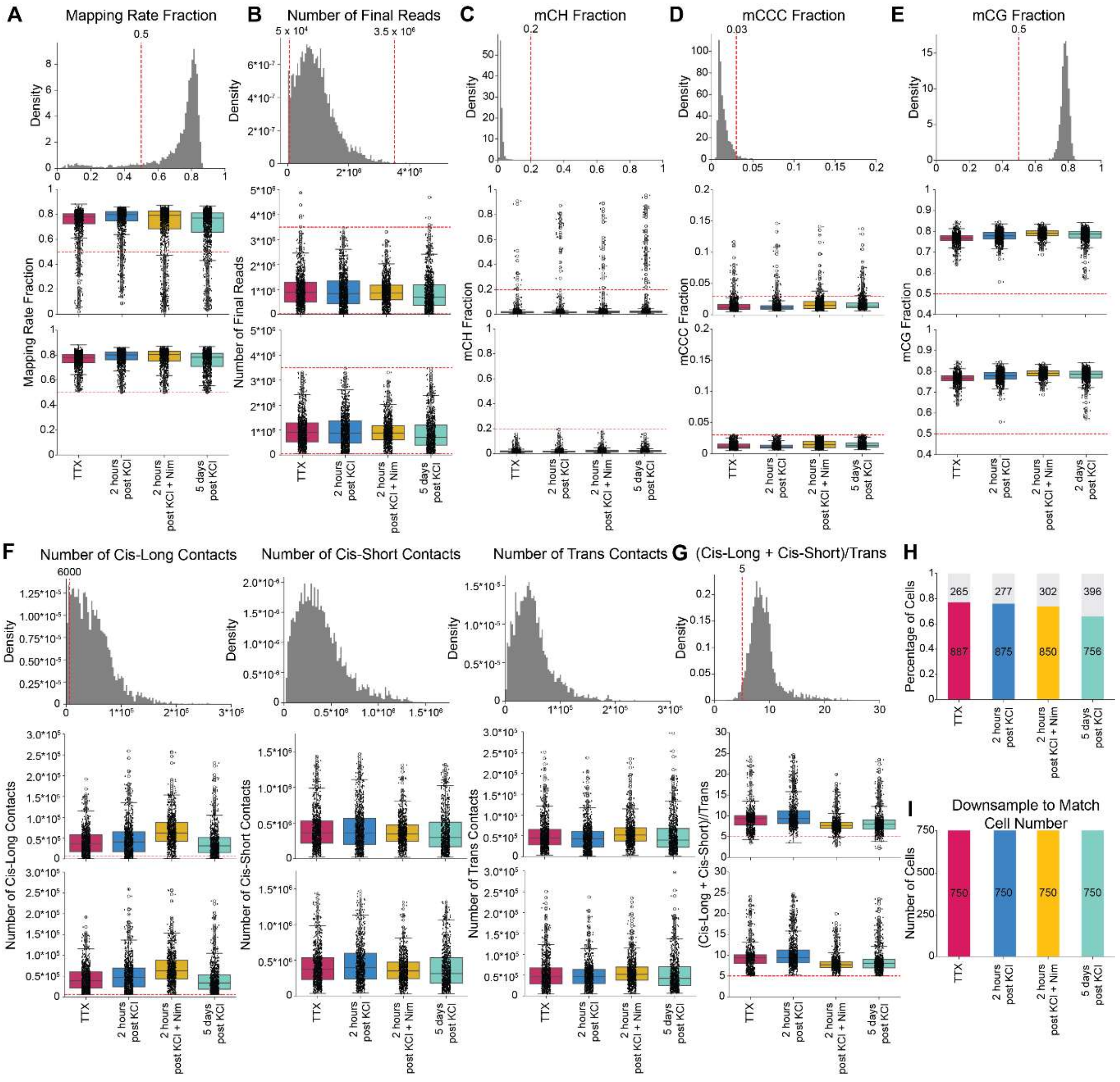


**Fig. S5. Filtering steps taken on snm3c-seq raw data at single cell level. (A-E):** Density plots (top row) and metrics per condition before (middle row) and after (bottom row) filtering out cells with a mapping rate of less than 0.5 **(A),** final reads less than 5,000 or greater than 3.5 million **(B)**, mCH fraction > 0.2 (**C**), mCCC fraction greater than 0.03 **(D)**, or an mCG fraction less than 0.5 **(E)**. **(F)** Density plots for number of cis-long contacts (left), number of cis-short contacts (middle) and number of trans contacts (right) before filtering, as well as number of contacts before (middle row) and after (bottom row) removing cells with less than 6,000 cis-long contacts. **(G)** Density plot and cis/trans ratio before (middle) and after (bottom) removing cells with a cis/trans ratio of less than five. **(H)** Percentage of cells remaining in each snm3c condition after all filtering steps. **(I)** Number of cells per condition after downsampling to match cell number.

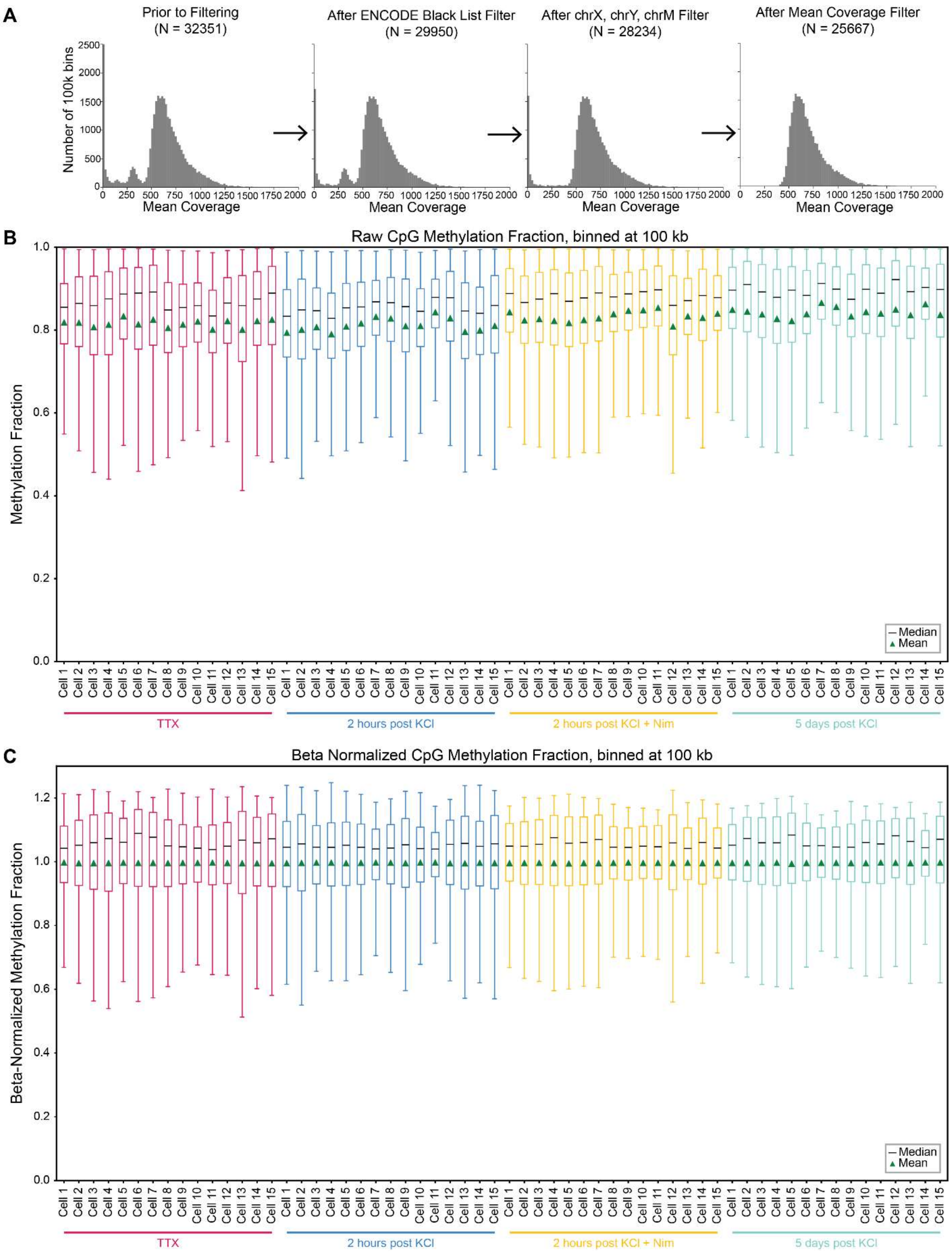


**Fig. S6. Beta normalization of methylation fraction. (A)** Mean coverage across 100k bins after filtering poor quality cells (far left), removing ENCODE blacklists (2nd from left), removing X, Y, and M chromosomes (second from right), and removing bins with a coverage of less than 400 or greater than 1600. (**B)** Distribution of raw CpG methylation fraction for 15 random cells per condition, binned at 100 kb. (**C)** Distribution of normalized CpG methylation fraction for the same 15 random cells per condition after beta normalization, binned at 100 kb.

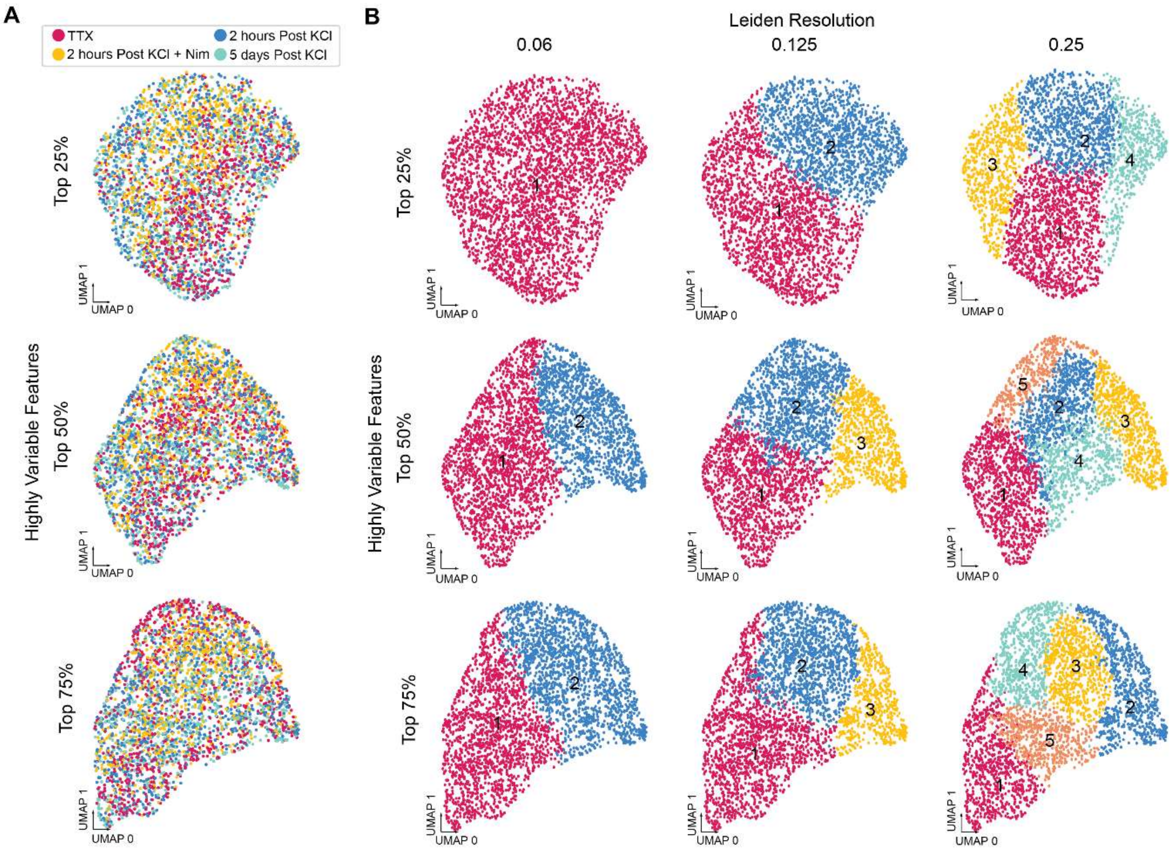


**Fig. S7. Clustering of snm3c filtered data beta-normalized at 100 kb bin size. (A)** UMAP colored by condition incorporating 25% (top), 50% (middle), or 75% (bottom) of Highly Variable Features (HVF). (**B)** Unbiased clustering at a Leiden Resolution of 0.06 (left), 0.125 (middle) and 0.25 (right) incorporating 25% (top), 50% (middle) or 75% (bottom) of HVF.

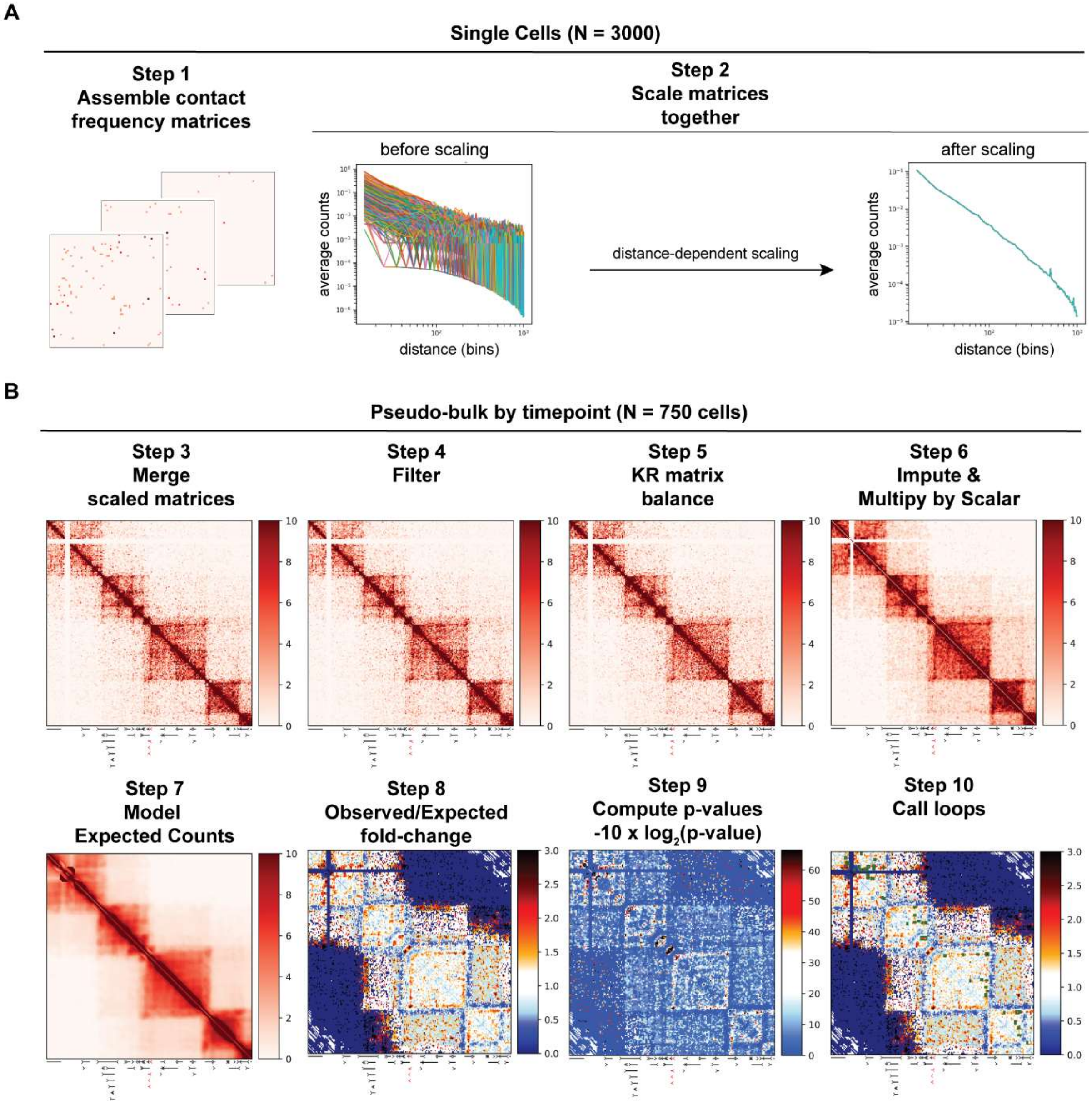


**Fig. S8. Loop calling pipeline using snm3C-seq data.** (**A**) Schematics showing the assembly (Step 1) and scaling of contact matrices across single cells using a custom distance-dependent scaling method (Step 2). (**B**) 30 kb resolution pseudo-bulk heatmaps showing contact frequencies within a 6 Mb region centered on the *BDNF* gene (chr11:24.6-30.7 Mb) at the 2 hour post-KCl timepoint. Each heatmap shows every step of the loop calling pipeline after single cell operations are completed. After merging scaled single cell matrices (Step 3), the next step is to filter out sparse rows and noisy pixels (Step 4), followed by Knight-Ruiz matrix balancing (Step 5), and performing random walk with restart (RWR) imputation followed by scalar multiplication by a power of 10 (Step 6). Then, expected "baseline" contact frequencies are calculated based on the donut expected model (Step 7) and the model fit is assessed by visualizing observed/expected fold-changes, where the "observed" corresponds to the matrix from step 6 (Step 8). P-values are then calculated (Step 9). The heatmap shown under Step 9 represents transformed p-values, -10 x log2(p-value), for visualization purposes only. Finally, loops are identified via clustering significant pixels by adjacency and are overlaid on top of the observed/expected (Step 10).

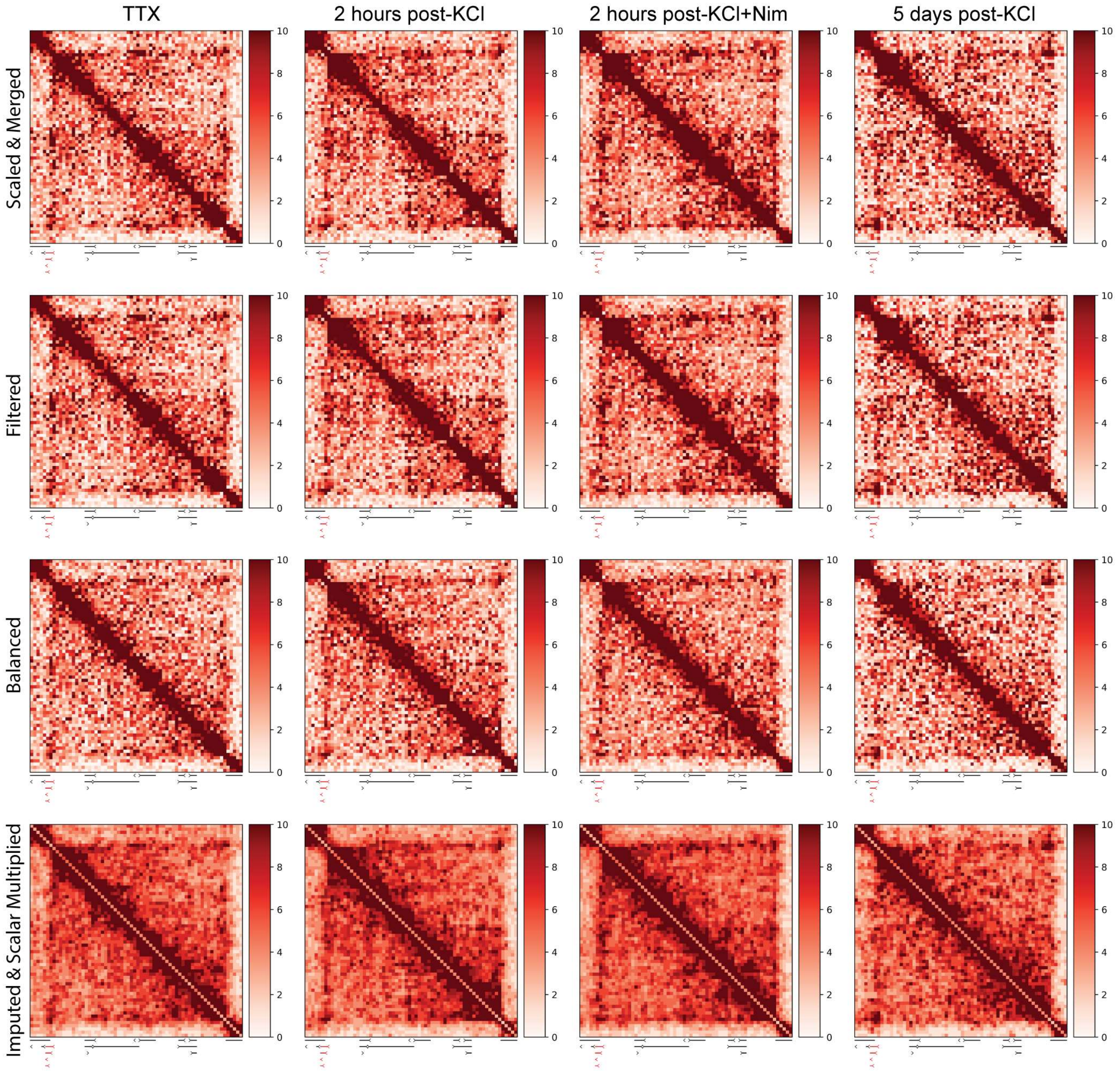


**Fig. S9. Locus-specific example of Fig. S8 normalization steps:** 30kb resolution pseudo-bulk heatmaps showing contact frequencies across all timepoints at a 2 Mb domain (chr11:27.5-29.5 Mb) containing the *BDNF* gene for each step of the loop calling pipeline leading up to the observed matrix: pseudo-bulking right after scaling (Step 3), filtering (Step 4), Knight-Ruiz matrix balancing (Step 5), and finally the observed matrix, which corresponds to Step 6 (imputation and scalar multiplication corresponding to a power of 10).

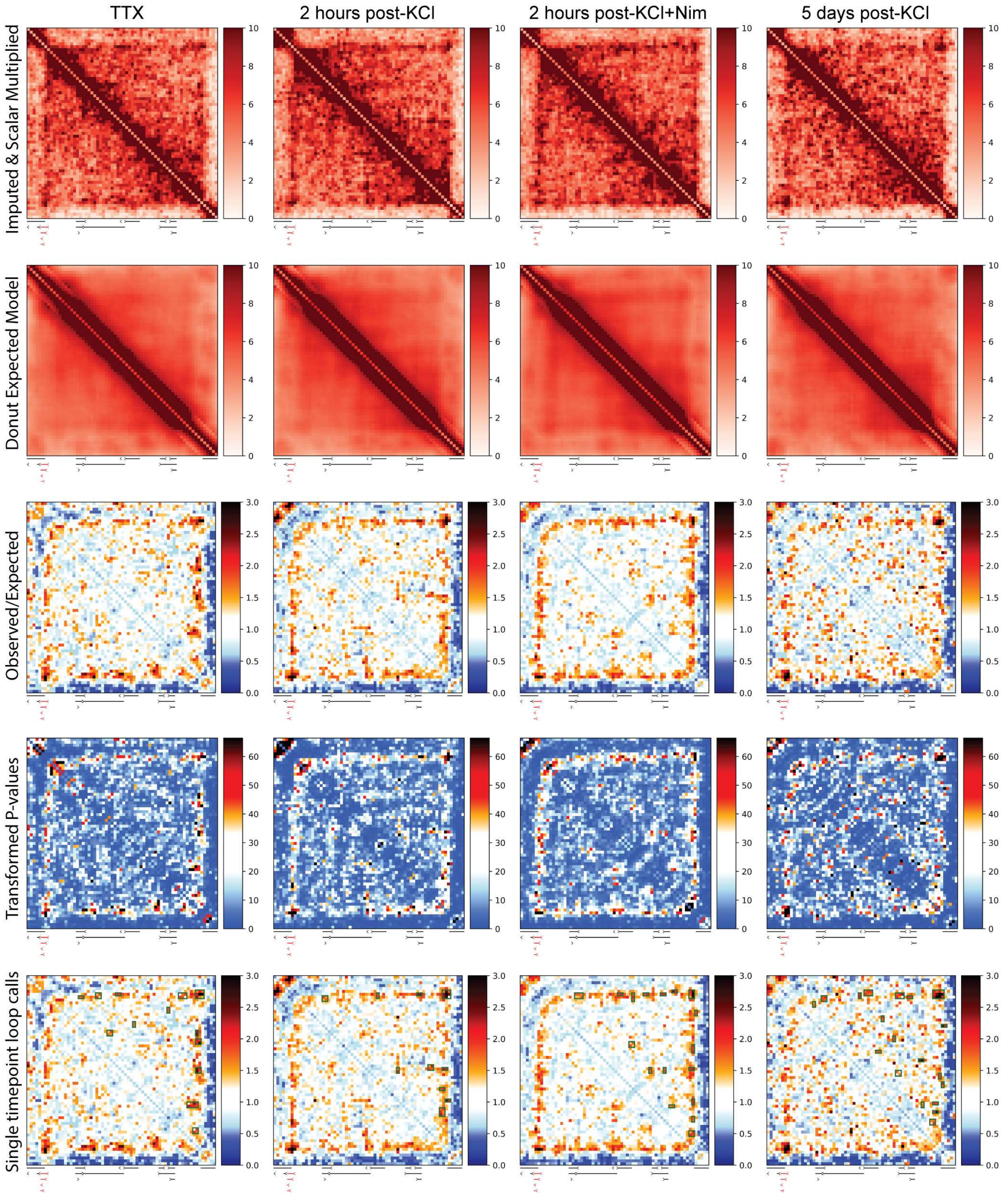


**Fig. S10. Locus-specific example of Fig. S8 loop calling steps:** 30kb resolution pseudo-bulk heatmaps showing contact frequencies across all timepoints at a 2 Mb domain (chr11:27.5-29.5 Mb) containing the *BDNF* gene for each step of the loop calling pipeline starting from the observed matrix: the observed matrix (Step 6), the donut expected model (Step 7), the observed/expected (Step 8), transformed p-values (Step 9), and loop calls overlaid on top of the observed/expected (Step 10).

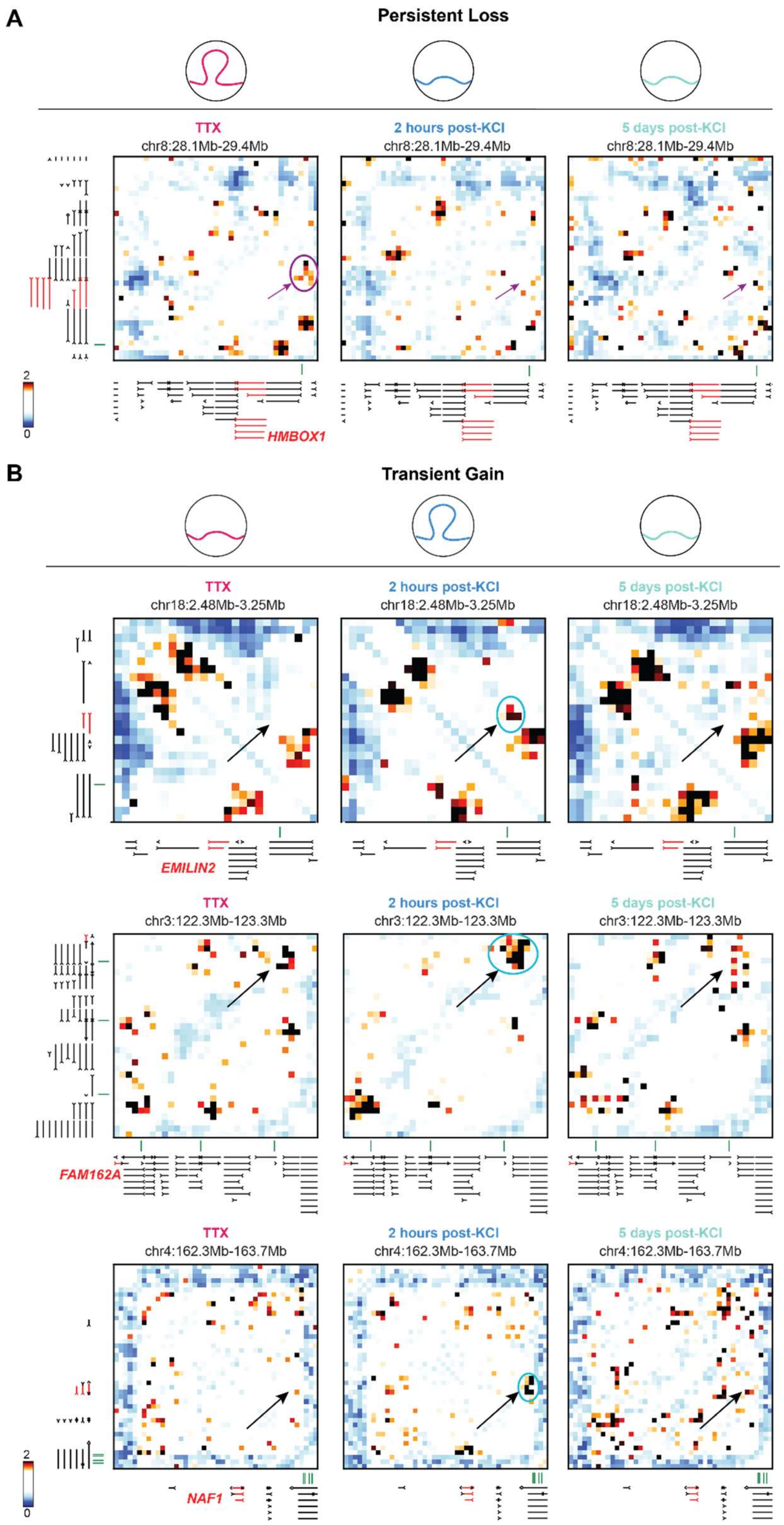


**Fig. S11. Example Heatmaps of Persistently Lost and Transiently Gained Loops: (A)** Observed over expected heatmap of persistently lost loop at *HMBOX1*. **(B)** Observed over expected heatmaps of transiently gained loops at *EMLIN2* (top), *FAM162A* (middle), and *NAF1* (bottom).

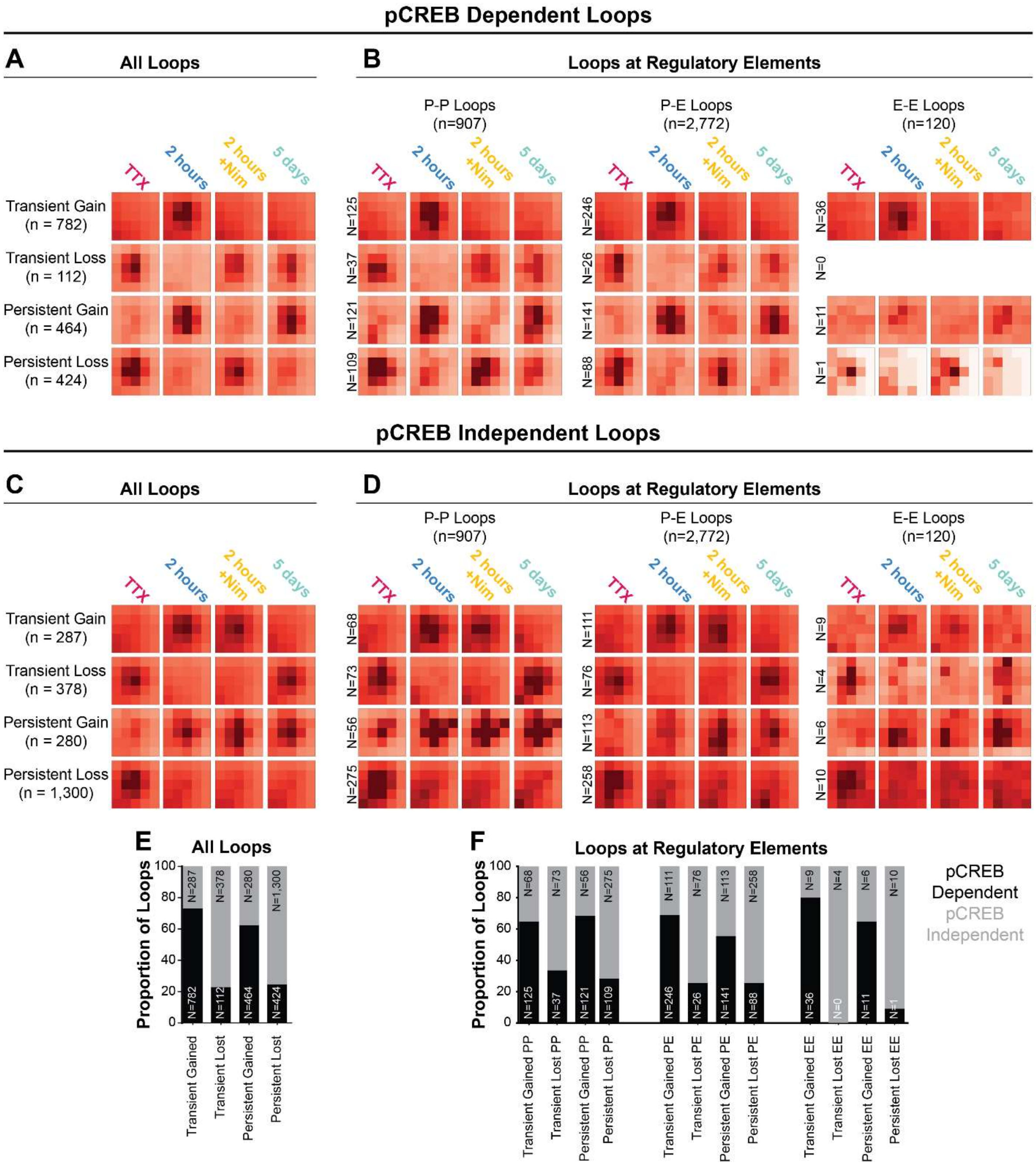


**Fig. S12. Nimodipine blocks a subset of activity-induced looping changes:** APAs for **A,C)** all loops in 5 looping classes and **B,D)** loops at regulatory elements for 5 looping classes for pCREB dependent (**A,B**) and pCREB independent **(C,D**) loops. **E,F)** stacked bar graphs for proportion of loops that are pCREB dependent or independent for **E)** all loop classes and **F)** loops at regulatory elements.

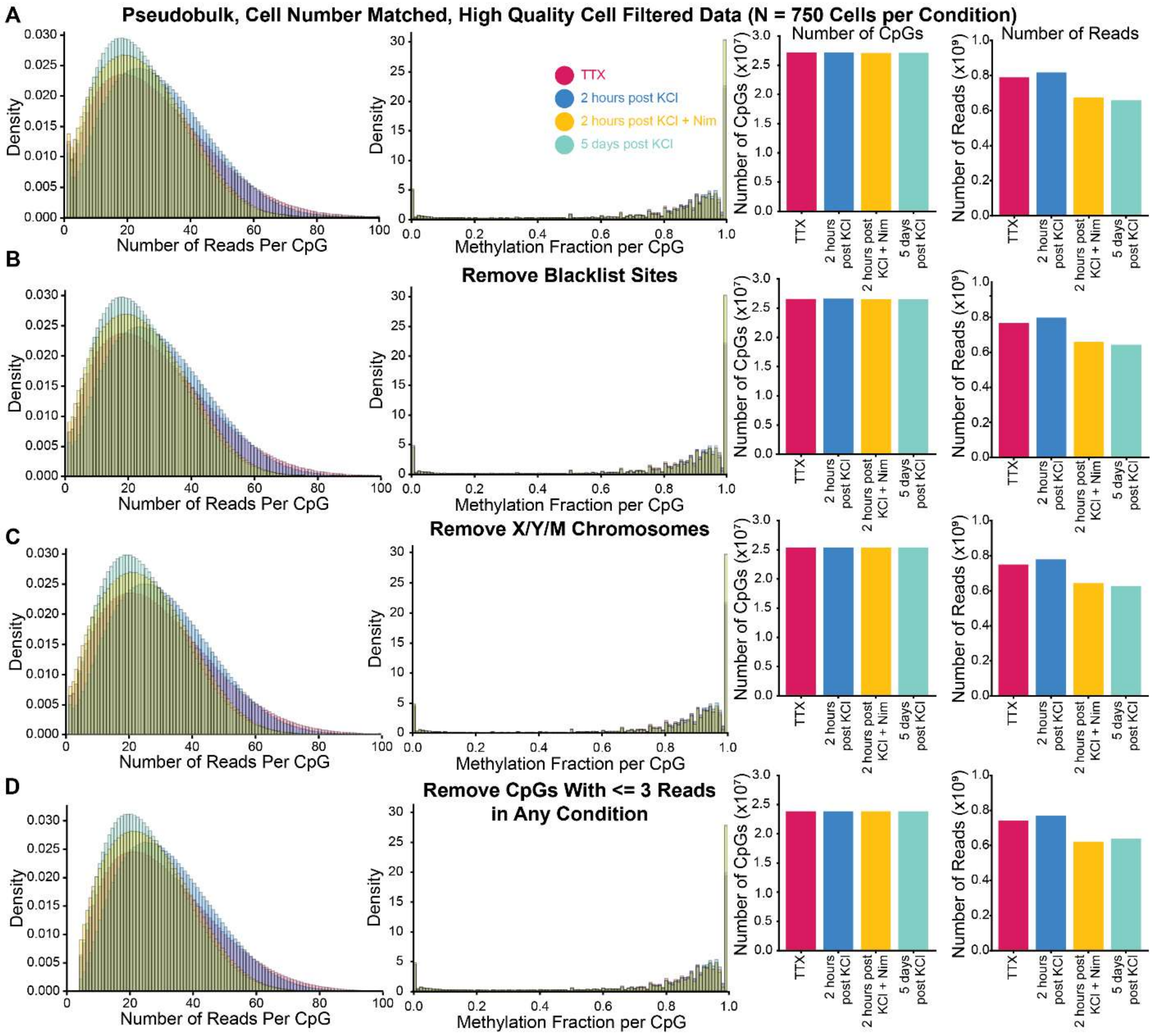


**Fig. S13. Filtering steps prior to calling differentially methylated loci (DMLs).** From left to right: Number of reads per CpG per condition, Methylation fraction per CpG per condition, Number of CpGs per condition, and Number of total reads per condition for **(A)** Pseudobulk, cell number matched, high quality cell filtered data (output of **Fig.** S5I); **(B)** after removing blacklist sites; **(C)** after removing chrX, chrY, chrM and any nonstandard chromosomes; and **(D)** Removing CpGs with 3 or fewer reads in any condition from all conditions.

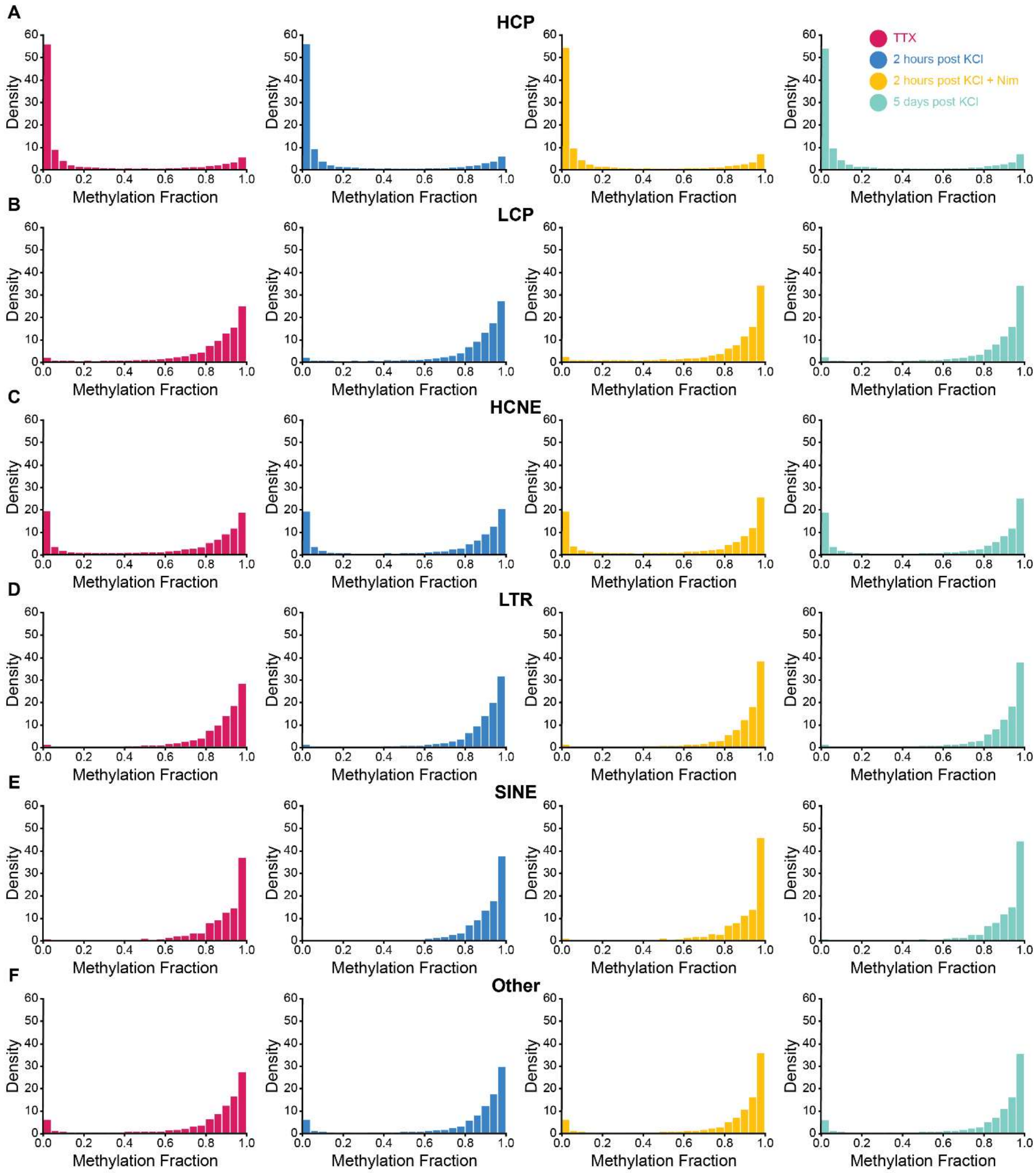


**Fig. S14. Measured methylation fractions at known mammalian genomic annotations are consistent with published results (*112*).** DNA methylation fractions at CpG resolution across conditions at **(A)** High-CpG density promoters (HCPs) (n = 631,232); **(B)** Low-CpG Density Promoters (LCPs) (n = 25,576); **(C)** highly conserved non-coding elements (HCNEs) (n = 545,343); **(D)** long terminal repeats (LTRs) (n = 1,886,294); **(E)** short interspersed nuclear elements (SINEs) (n = 6,920,181) and **(F)** All other genomic features (n = 15,146,283).

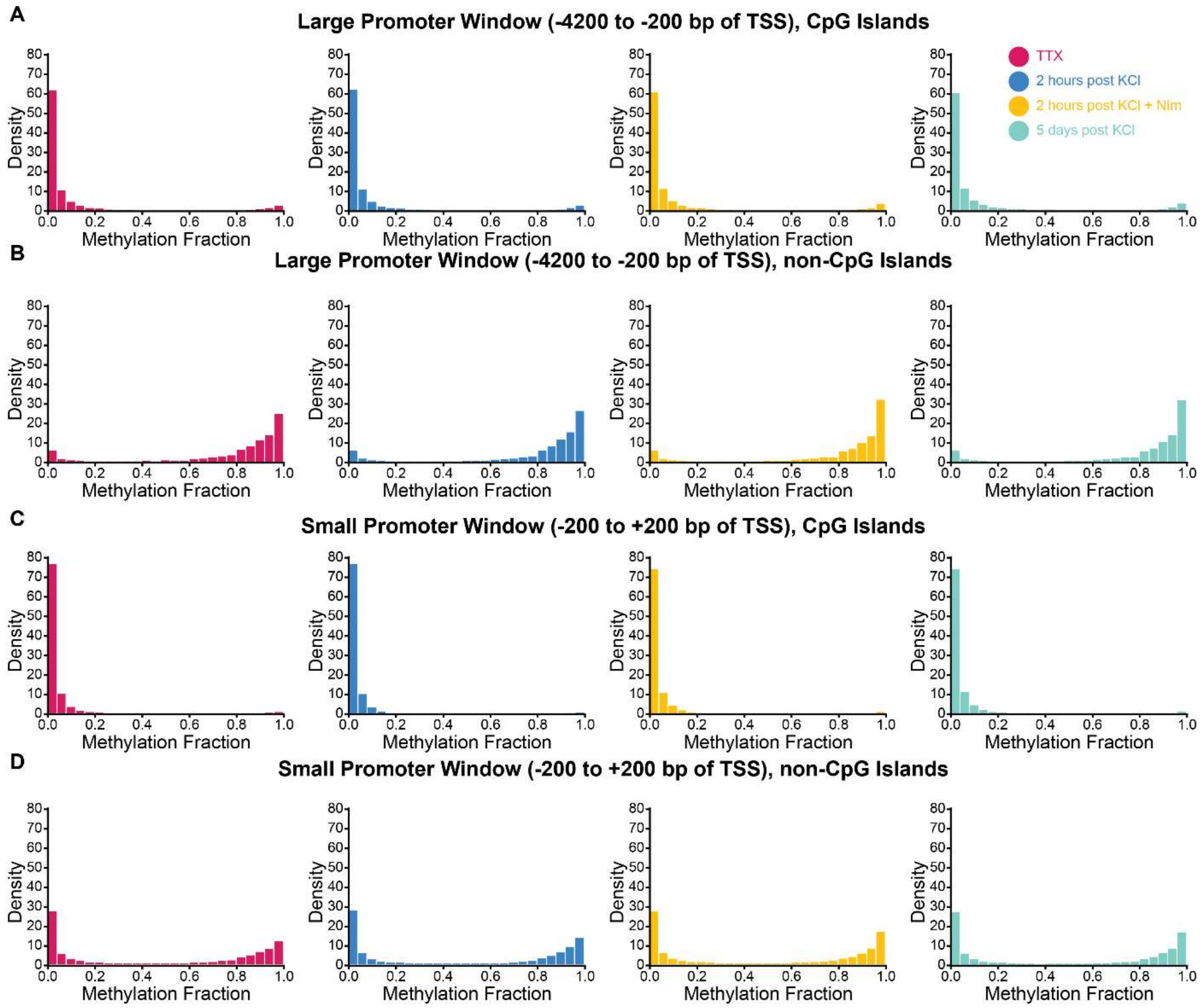


**Fig. S15. Measured methylation fractions at promoters +/- CpG islands.** DNA methylation fractions at CpG level across conditions for CpGs within **(A)** Large promoter windows (-4200 bp to -200 bp of TSS) and CpG islands (n = 452,725); **(B)** Large promoters windows and not in CpG islands (1,396,117); **(C)** Small promoter windows (-200 to +200 bp of TSS) and CpG islands (n = 500,436); and **(D)** Small promoter windows and not in CpG islands (n = 137,481).

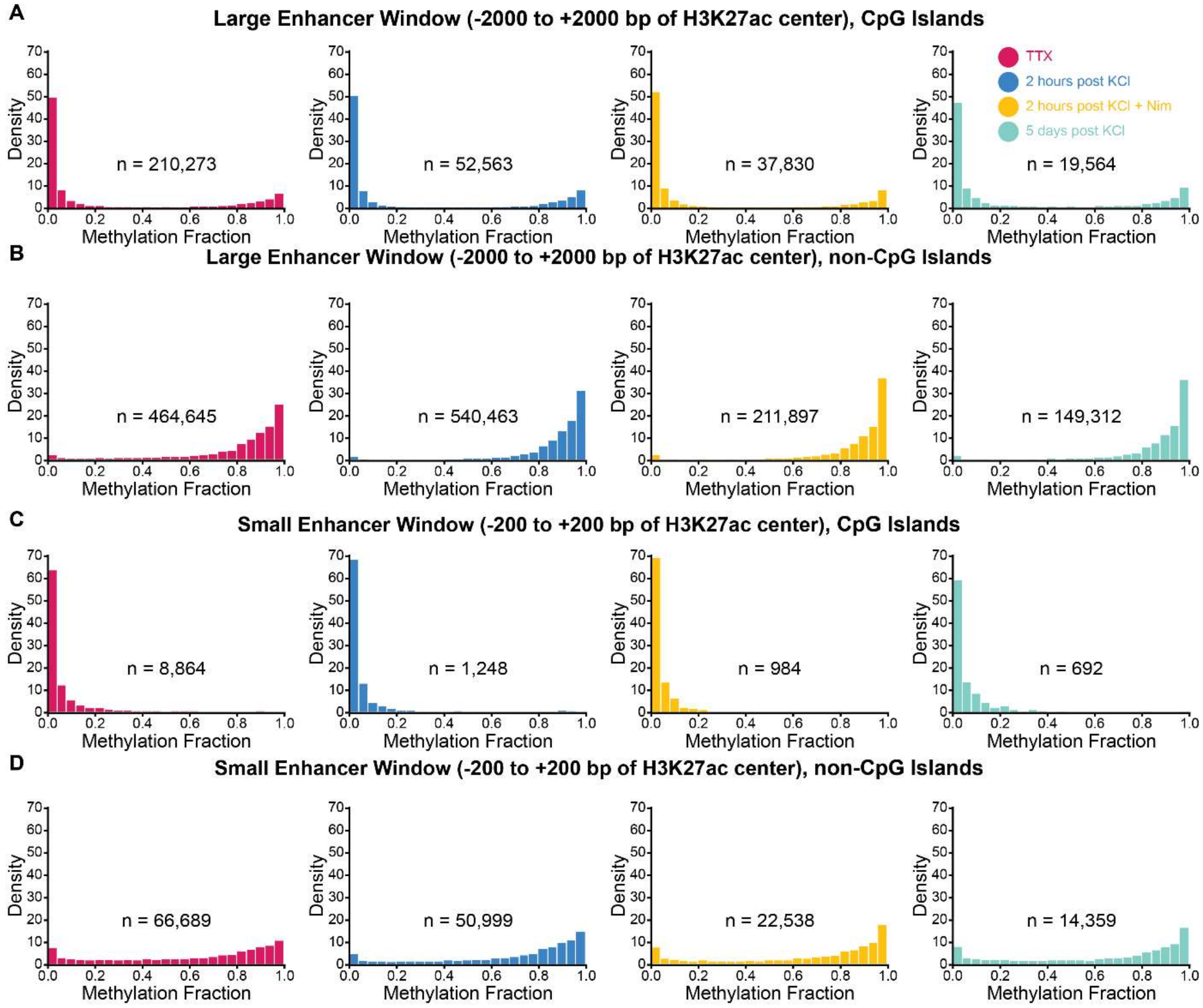


**Fig. S16. Measured methylation fractions at putative activity-decomissioned and activity-induced non-coding enhancers +/- CpG islands.** DNA methylation fractions at CpG level across conditions for CpGs within **(A)** Large enhancer windows (-2000 to 2000 bp from H3K27ac center) and CpG islands; **(B)** large enhancers windows and not within CpG islands; **(C)** Small enhancer windows (-200 to 200 bp form H3K37ac center) and CpG islands; and **(D)** Small enhancer windows and not within CpG islands.

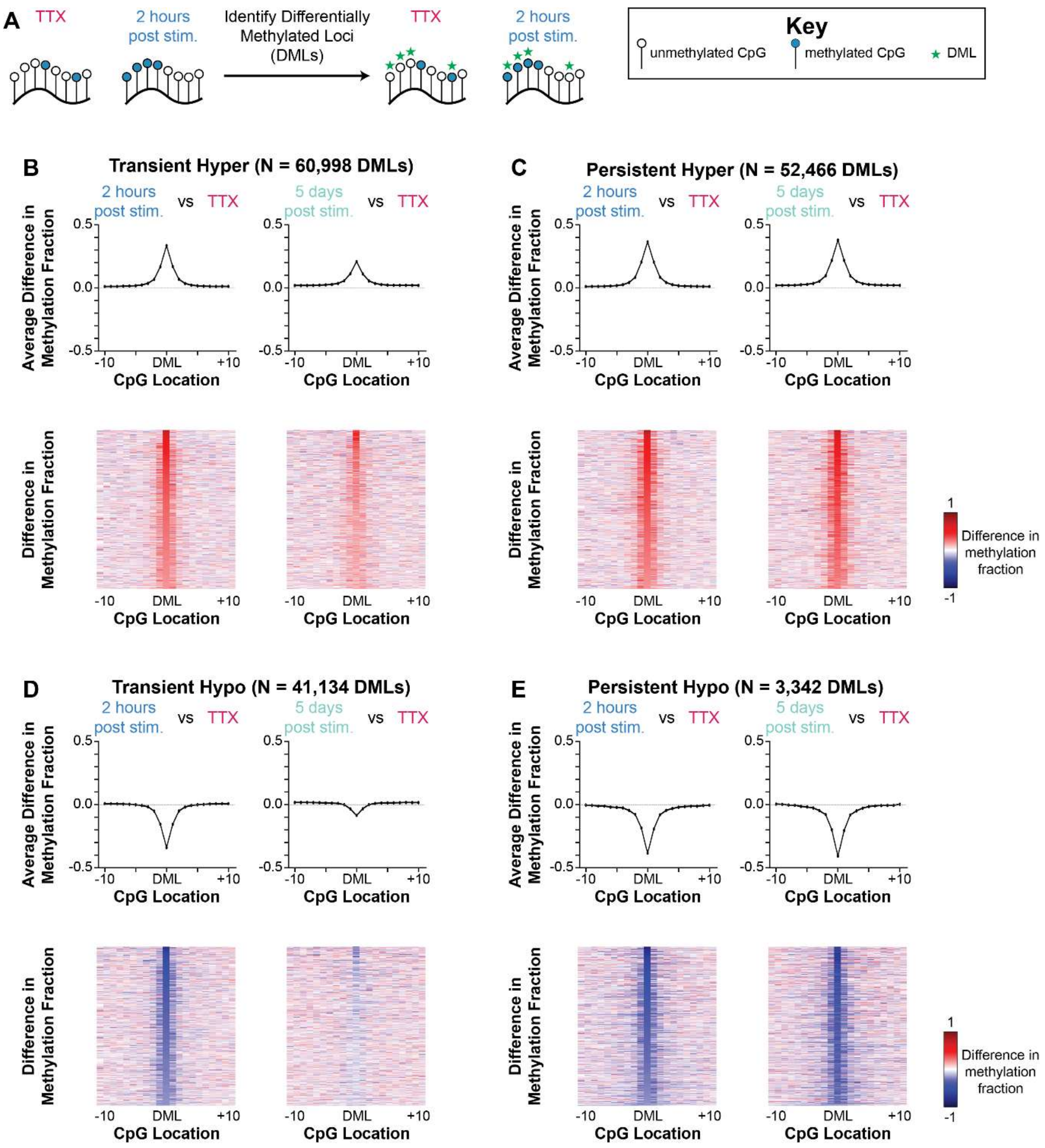


**Fig. S17. KCl induction of differential methylation at CpGs in human neurons. A)** Schematic of how differentially methylated loci (DMLs) are defined. **B-E)** Pileups (top row) and heatmaps (bottom row) of difference in methylation fraction for 2 hours post KCl vs TTX (left) and 5 days post KCl vs TTX (right) for **B)** Transiently hypermethylated CpGs; **C)** Persistently hypermethylated CpGs; **D)** Transiently hypomethylated CpGs; and **E)** Persistently hypomethylated CpGs.

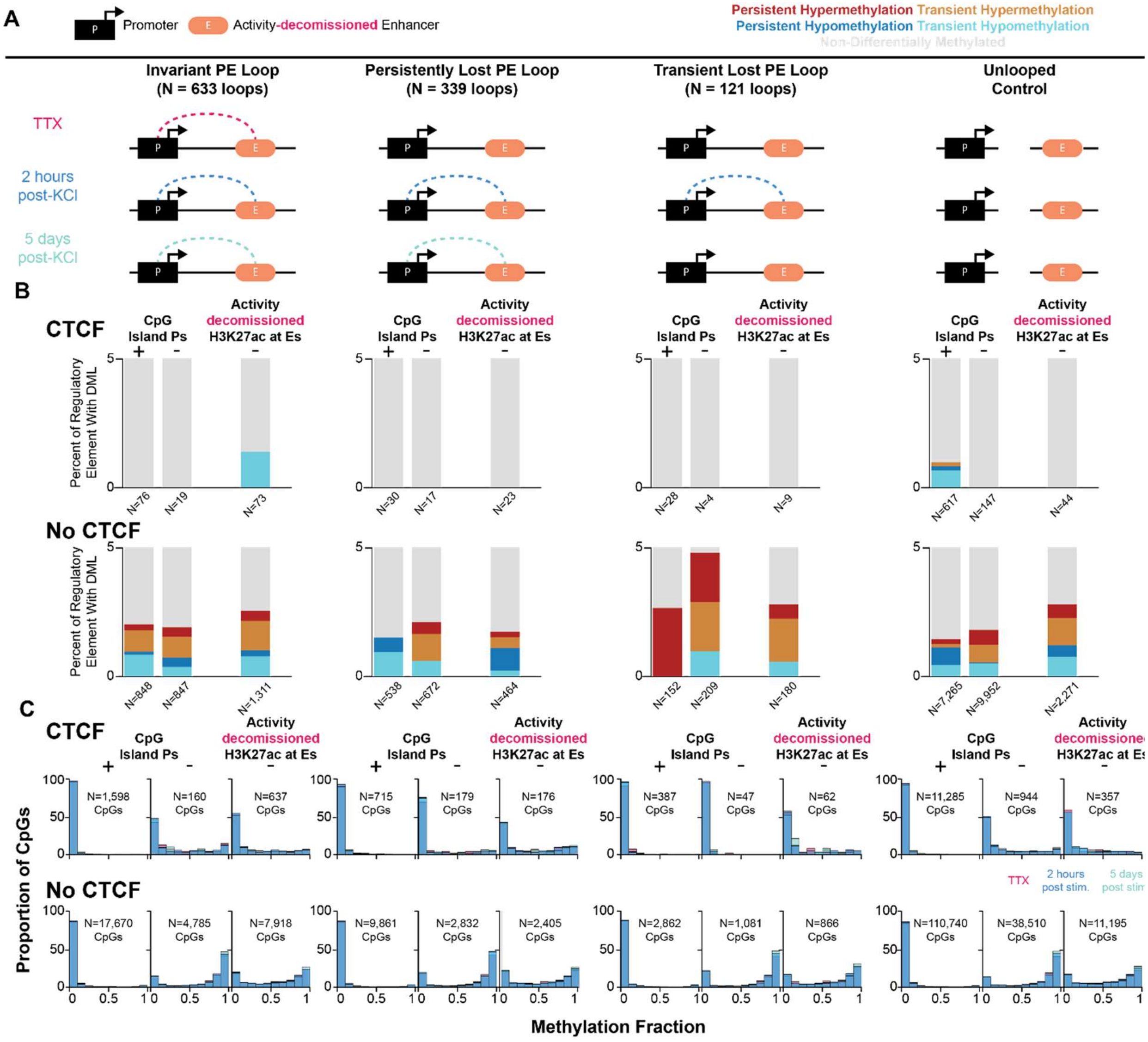


**Fig. S18. Regulatory elements at persistently and transiently lost loops bound by CTCF are devoid of changes in DNA methylation upon stimulation of human iPSC-derived neurons.** (**A**) From left to right: schematics of the invariant PE loops, persistently lost PE loops, transiently lost PE loops, and unlooped controls for putative noncoding enhancers that lose H3K27ac upon KCl stimulation (activity-decommissioned enhancers). **(B)** Percentage of regulatory elements with each type of DML for four looping classes defined in **(A)**. From left to right within a given class: promoters with CpG islands, promoters without CpG islands, and enhancers that lose H3K27ac upon KCl stimulation. Upper row represents regulatory elements bound by CTCF, and lower row represents regulatory elements not bound by CTCF. **(C)** Histograms of methylation fraction for each group defined in **(B)**.

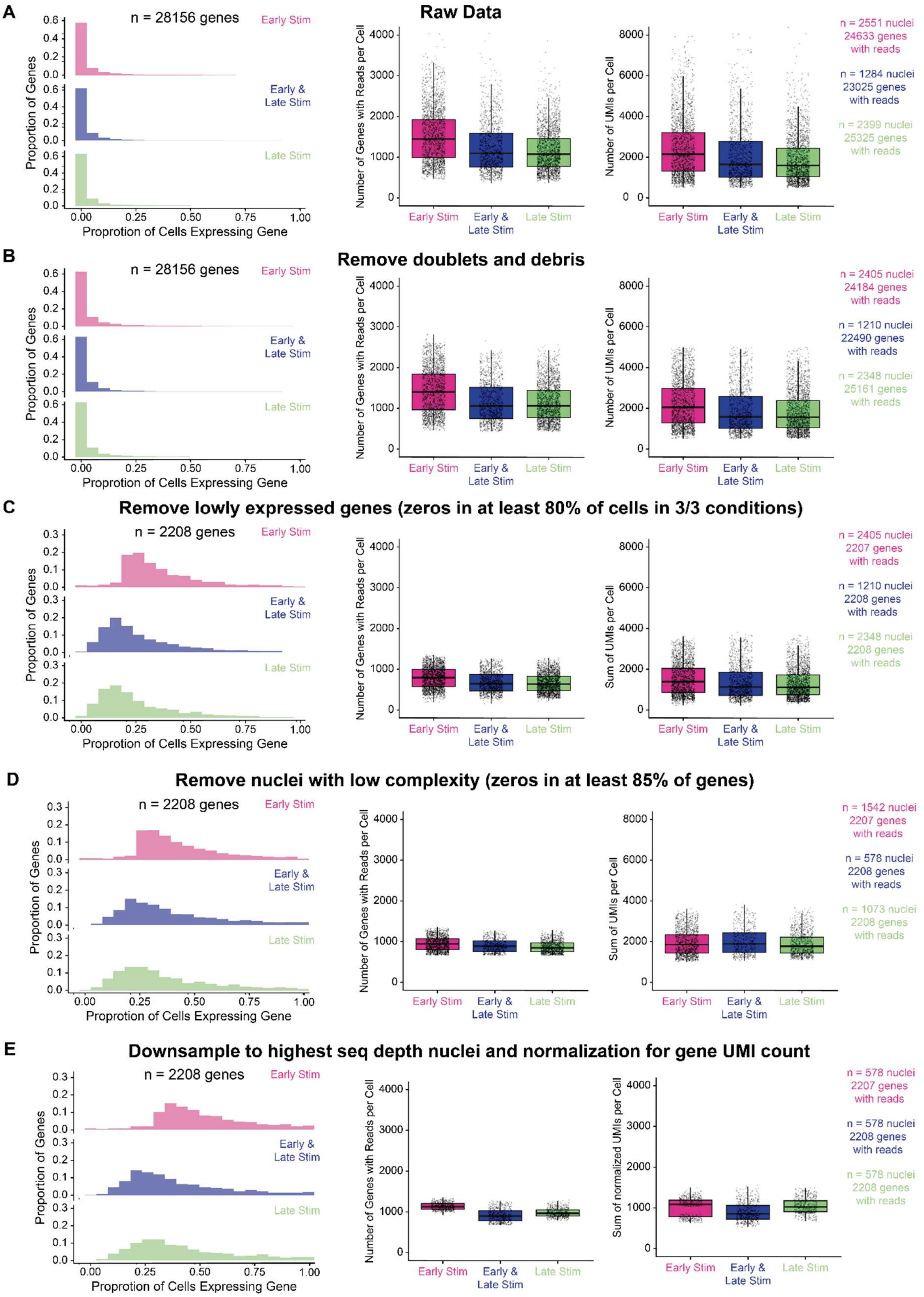
A
Raw Data
n = 28156 genes
Early Stim
Early & Late Stim
Late Stim
Proportion of Genes
Proprotion of Cells Expressing Gene
Number of Genes with Reads per Cell
Number of UMIs per Cell
n = 2551 nuclei 24633 genes with reads
n = 1284 nuclei 23025 genes with reads
n = 2399 nuclei 25325 genes with reads
B
Remove doublets and debris
n = 28156 genes
n = 2405 nuclei 24184 genes with reads
n = 1210 nuclei 22490 genes with reads
n = 2348 nuclei 25161 genes with reads
C
Remove lowly expressed genes (zeros in at least 80% of cells in 3/3 conditions)
n = 2208 genes
Sum of UMIs per Cell
n = 2405 nuclei 2207 genes with reads
n = 1210 nuclei 2208 genes with reads
n = 2348 nuclei 2208 genes with reads
D
Remove nuclei with low complexity (zeros in at least 85% of genes)
n = 2208 genes
n = 1542 nuclei 2207 genes with reads
n = 578 nuclei 2208 genes with reads
n = 1073 nuclei 2208 genes with reads
E
Downsample to highest seq depth nuclei and normalization for gene UMI count
n = 2208 genes
Sum of normalized UMIs per Cell
n = 578 nuclei 2207 genes with reads
n = 578 nuclei 2208 genes with reads
n = 578 nuclei 2208 genes with reads

**Fig. S19. Filtering and normalization of single-nucleus RNA-seq data and multiple quality control steps.** Proportion of cells expressing a gene in each condition (left), number of genes with reads per cell (middle), and number of unique molecular identifiers (UMIs) per cell (right) for **A)** raw data; **B )** data after removing doublets and debris; **C)** data after dropping nuclei low complexity; **D)** filtering for the top 300 nuclei in blue condition; **E)** downsampling to 300 nuclei across conditions; **F)** removing lowly expressed genes; and **G)** UMI scaling

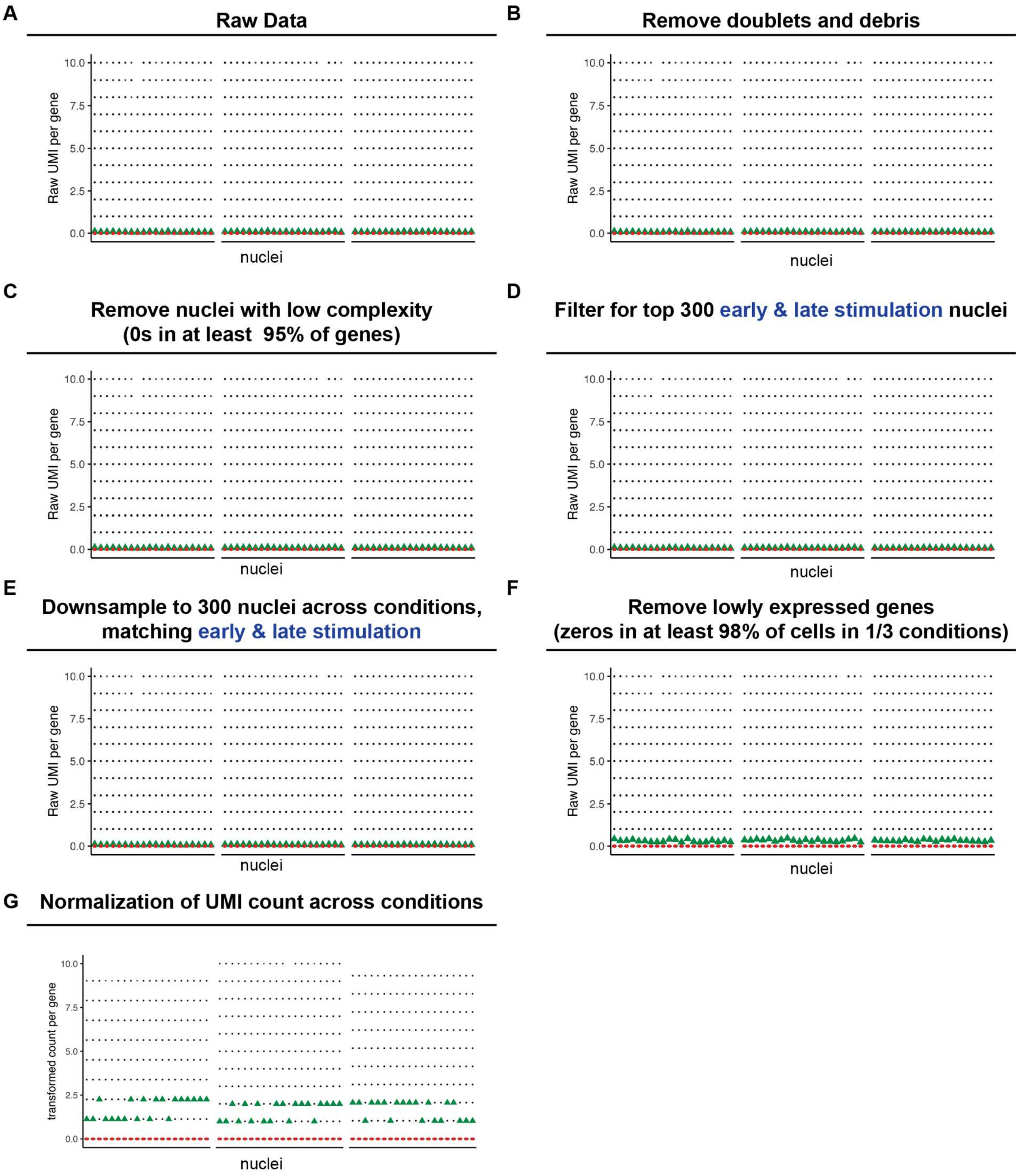


**Fig. S20. Filtering and normalization of single-nucleus RNA-seq data and multiple quality control steps for 20 cells per condition.** Log transformed counts per gene for the same 20 random cells per condition for **A)** raw data; **B)** after removing doublets and debris; **C)** after filtering nuclei; **D)** filtering for top 300 nuclei in blue condition; **E)** downsampling to 300 nuclei across conditions; **F)** removing lowly expressed genes **G)** Normalizing UMI count across conditions

A

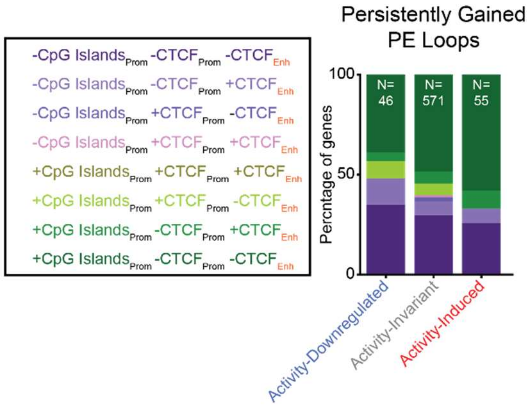


**Fig. S21. CTCF occupancy and CpG island status at promoters and enhancers of genes in persistently gained PE loops: (A)** Proportion of loops with CTCF occupancy at enhancers and promoters with and without CpG islands for activity-downregulated (left), activity-invariant (middle) and activity-induced genes at persistently gained PE loops.

#### Supplementary Tables

**Table S1.**
Data produced in Waldman, Pham et al.

**Table S2.**
Single-nucleus methyl 3C-seq barcode sequences

**Table S3.**
Microelectrode array settings, data, and analysis

**Table S4.**
Gene Expression with bulk nuclear RNA-seq

**Table S5.**
CUT&Tag H3K27ac peak calls

**Table S6.**
Putative noncoding activity-dependent enhancers

**Table S7**.
CTCF occupied sites genome-wide in human iPSC-derived neurons

**Table S8.**
Loop calls from snm3c-seq data across a human neural stimulation timecourse

**Table S9.**
Loop classes in human iPSC-derived neurons

**Table S10.**
Gained loop classes stratified by dependence on activity-induced CREB phosphorylation

**Table S11.**
Lost loop classes stratified by dependence on activity-induced CREB phosphorylation

**Table S12.**
Classes of activity-dependent gene expression across the timecourse of stimulation of human iPSC-derived neurons

**Table S13.**
Biologically relevant expression patterns for genes anchoring persistent promoter-enhancer loops and unlooped control

#### Materials and Methods

Some of the methods described below have been detailed in our own previous manuscripts (*5, 14-28, 70, 72, 113-120*). To ensure reproducibility and clarity, we state the same or near similar methodological steps below.

***Human induced pluripotent stem cell (hiPSC) Culture***

All $i^3N$ (WTC11.G3-WT, Gladstone) hiPSC cultures were maintained on Matrigel (Corning, 354277) / DMEM/F12 (Fisher, 11320033) coated plates at 37 ºC and 5% $CO_2$. The Matrigel:DMEM/F12 ratio was determined according to the manufacturer's dilution ratio. The $i^3N$ cell line contains a doxycycline (Dox) inducible Neurogenin2 (*Ngn2*) transgene inserted into the adeno-associated virus integration site 1 (AAVS1) of the WTC11 hiPSC line (*57, 58*) (**Fig. S1A).**

hiPSCs were cultured in 100 mm x 20 mm cell culture plates (Corning, 430167) in Cell Culture Media (mTeSR+ (STEMCELL Technologies, 100-0274) supplemented with 5X mTeSR+ Supplement (STEMCELL Technologies, 100-0275), and 5 mL Penicillin/ Streptomycin (PS) (Gibco, 15140-122) (1% final concentration of PS)). When cells reached 70-80% confluency, they were expanded up to 150 mm x 25 mm cell culture plates (Corning, 430599). Cell Culture Media was aspirated, cells were washed once with 1X $Mg^{2+}$ and $Ca^{2+}$-free PBS (Corning, 21-040-CV) and then incubated in half culture volume Versene (Thermo Fisher, 15040066) for 8 minutes at 37 ºC, or until cells had fully dissociated. Versene was neutralized with half culture volume Cell Culture Media, and cells were removed from cell culture plate, spun down at 250 xg for 3 minutes, and resuspended in fresh mTeSR+ before replating.

***Differentiation of hiPSCs into Functional Cortical Neurons***

*Neuronal Cell Culture Plate Coating*

To create a positively charged surface, cell culture plates were coated with 0.01% poly-l-ornithine (Sigma-Aldrich, P4957) (PLO) for 8-24 hours on a shaker at 100 RPM at room temperature. To promote neural adhesion, PLO was then aspirated, and 20 µg/mL laminin (Thermo, 23017015) in 1X Mg and Ca-free PBS was added. Plates were again incubated on a shaker at 100 RPM overnight. Laminin was then aspirated, and half culture volume of Induced Neuron Media (Neurobasal (Gibco, 21103), 1X B27 (ThermoFisher, 17504044), 1X glutamax (Life Technologies, 35050-061), 1% PS) was added. To continue neural differentiation, 1 µg / mL doxycycline (Millipore Sigma, D3347) was added to cell culture plates on the day of replating differentiating cells. Plates were warmed to 37 ºC.

*Neuronal Induction, Replating, and Differentiation*

When $i^3N$ cultures reach 40-50% confluency, 1 µg/mL doxycycline was added to media to induce differentiation. Throughout this manuscript, DIV-2 refers to $i^3N$ in the hiPSC state prior to neural induction. Cells were maintained for 48 hours in Cell Culture Media supplemented with 1 µg / mL dox, during which time plate coating was performed as described in *Cell Culture Plate Coating*. This solution was then aspirated, and half culture volume of Induced Neuron Media (Neurobasal (Gibco, 21103), 1X B27 (ThermoFisher, 17504044), 1X glutamax (Life Technologies, 35050-061), 1% PS, 10 ng/mL BDNF (Peprotech, 450-02), 10 ng/mL NT-3 (Peprotech, 430-03)) supplemented with 1 ug /mL doxycycline, 10 µM DAPT (Selleckhem, S2215), 5 µM 5-Fluorouracil (Sigma Aldrich, F6627-1G) and Uridine (Sigma Aldrich, U3750-1G), was added to pre-warmed plates at 37 ºC. At DIV0, after 48 hours of doxycycline induction at 37 ºC, induced neurons were replated onto PLO/laminin-coated plates. To do so, induced neurons were dissociated with half culture volume Accutase (Thermo Fisher, A1110501) for eight minutes at 37 ºC, neutralized with half culture volume Cell Culture Media, and centrifuged to pellet cells at room temperature for 3 minutes at 250 xg. Cells were then resuspended in 1 mL Induced Neuron Media and gently pipetted up and down to break up the cell pellet, at which time 4 more mL neuron media was added. This 5 mL of media was passed through a 70 µM cell strainer to remove clumps. Cells were counted on a hemocytometer, diluted to 78,000 cells per square centimeter of cell culture plate area, and replated (for example, for a 100 mm plate, cells were diluted to 442,260 cells per mL of cell suspension, and 10 mL of cell suspension was then added to the previously PLO/laminin coated plates). 24 hours after the initial replate (DIV0), a full media change plus doxycycline was done (DIV1). Following this, every 7 days from the initial replate of induced neurons, a half media change of Induced Neuron Media was performed. Dox was not included in all subsequent half media changes with Induced Neuron Media after the DIV1 full media change. Differentiating neurons were incubated at 37 ºC with 5% $CO_2$.

***Potassium Chloride Stimulation of Neurons***

5 weeks after cells were replated (DIV34), cells were pretreated with a half media change of 2 µM Tetrodotoxin (TTX) (Cayman Chemical, 14964-1) in neuron media (final concentration 1 µM) for 24 hours to silence neuronal firing (*5*). After 24 hours, the *TTX* condition was collected, and High KCl Artificial Cerebral Spinal Fluid (ACSF) (122.6 mM sodium chloride (NaCl), 26.2 mM sodium bicarbonate ($NaHCO_3$), 50 mM potassium chloride (KCl), 1.2 mM monosodium phosphate ($NaH_2PO_4$), 2 mM calcium chloride ($CaCl_2$), 10 mM D-glucose in Cell Culture Grade Water) (*121*) was added to all other conditions. Cells received either high KCl no MG ACSF with no inhibitors, or with the addition of 5 µM Nimodipine (abbreviated as 'Nim') (Thermo Fisher, 329280010) for the nimodipine conditions (*59, 122*). Cells were incubated for one hour in ACSF at 37 ºC, after which time they were returned to Induced Neuron Media. The *KCl one-hour* timepoint was collected immediately, and *2 hours post-KCl, 5 hours post-KCl,* and *5 days post-KCl* timepoints were collected 2 hours, 5 hours, or 5 days after the media change back to Induced Neuron Media.

***Immunofluorescence and Confocal Microscopy***

*Plating Cells for Immunofluorescence*

To plate cells for immunofluorescence, 18 mm coverslips (Warner Instruments, 64-0714) were placed in 80 % ethanol (Decon Laboratories, 2716) in Cell Culture Grade Water (Corning, 25-055-CM) for 15 minutes, the ethanol was aspirated, and coverslips were rinsed with pure ethanol and allowed to air dry. One coverslip was then placed per well of a 12 well Cell Culture Plate (Corning, 3513). Coverslips were either coated in matrigel/DMEM/F12 or PLO/laminin, as described above in ***Human induced pluripotent stem cell (hiPSC) Culture*** or *Neuronal Cell Culture Plate Coating*, respectively, depending on if cells to be plated were hiPSCs or neurons. Cells were plated on coated coverslips as described above. After coating, hiPSCs were grown to 40% confluency, and neurons were grown to DIV35, prior to fixation.

*Fixation for Immunofluorescence*

Cells were fixed in 4% paraformaldehyde (Thermo Fisher, 28908) in PBS for 10 minutes. Media was then aspirated, and cells were washed 3x with PBS, with five-minute room temperature incubations in PBS between each wash step. Plates were sealed with parafilm and stored in 1 mL PBS at 4 ºC in the dark until use.

*Immunofluorescence Staining*

Fixed cells were permeabilized in 0.2% Triton X-100 (Sigma Aldrich, 93443-100mL) in PBS for 10 minutes. Media was aspirated, and cells were washed 3X with PBS. Cells were then blocked in Blocking Solution (filtered 5% BSA (NEB, B9000S) in PBS) for 1 hour at room temperature. To prepare Primary Antibody Solutions, primary antibodies were diluted to their required concentration in Blocking Solution and spun down for 5 minutes at 11,600 xg to remove precipitates. Blocking solution was aspirated from cells, and the primary antibody solution was added. Plates were sealed with parafilm and incubated overnight at 4 ºC in the dark. Secondary Antibody Solutions were prepared by diluting secondary antibodies to their required concentration in filtered 5% BSA in PBS and spinning down at for 5 minutes 11,600 xg to remove precipitates. Primary Antibody Solution was then aspirated from cells, cells were washed 3X with PBS for five minutes per wash step, and Secondary Antibody Solution was added. Cells were incubated at room temperature for 2 hours in Secondary Antibody Solution, media was aspirated, and cells were washed 3X with PBS. Coverslips were then mounted facedown onto a glass slide with 10 µL VECTASHIELD Antifade Mounting Medium with DAPI (Vector Laboratories, H-1200). Coverslips were then sealed onto the glass slide with nail polish and stored at 4 ºC. The following antibodies were used in this study: rabbit anti-OCT4, 1:200 dilution (Cell Signaling Technologies, 2750S); mouse anti-NANOG, 1:500 dilution (Sigma-Aldrich, MABD24); mouse anti-CUX1, 1:200 dilution (Santa Cruz Biotechnology, sc-514008); rabbit anti-NeuN, 1:100 dilution (Cell Signaling Technologies, 24307S); chicken anti-MAP2, 1:1000 dilution (Abcam, ab5392); mouse anti-Synapsin-1, 1:500 dilution (Synaptic Systems, 106011), rabbit anti-Homer1, 1:500 dilution (Synaptic Systems, 160 008); Guinea Pig anti-cFOS, 1:500 dilution (Synaptic Systems, 226004), Goat anti-Rabbit 488, 1:1000 dilution (Thermo Fisher, A11034); Donkey anti-Mouse 647, 1:1000 dilution (Thermo Fisher, A31571); Donkey anti-mouse 594, 1:1000 dilution (Thermo Fisher, A21203), Goat anti-chicken 647, 1:1000 dilution (Thermo Fisher, A21449), Goat anti-Guinea Pig 647 (abcam, ab150187).

*Immunofluorescence Imaging*

Leica LAS X software (v3.5.2.18963) and a TCS SP8 Multiphoton Confocal (Leica TCS SP8 Multiphoton Confocal) microscope were used for imaging. Images were processed in FIJI (v2.14.0/1.54f) and Adobe Illustrator, where image brightness and contrast levels were equivalently adjusted across images to provide higher quality images. Imaging parameters were selected using secondary only controls. All images taken for multiple experimental conditions were acquired at identical acquisition settings.

*Computational Image Analysis*

The expression of cFOS and DAPI for 256 nuclei treated with TTX, 253 nuclei treated with KCl, and 197 nuclei treated with KCl + Nim was quantified from z-stack images using Cellpose (version 0.7.2) (*123*). The z-stack images contained channels for DAPI staining as well as for cFOS. Cellpose's built in nuclei model with a diameter value of 75 pixels on DAPI stained images was used to segment the nuclei. The stitch_threshold was set to 0.5. Once segmentation was finished, a custom script was used to quantify the volume of every nucleus segmented as well as the total sum of pixel intensity within the cFOS and DAPI channels. The total sum of cFOS and DAPI pixel intensity was normalized by dividing the sum by the nuclei volume, yielding the average cFOS or DAPI signal for a given nucleus. Finally, using the average DAPI signal per nucleus and the nuclear volume, nuclei that were too small or too large, or had too high or too low DAPI signal were filtered, as these values indicated segmentations of debris or nuclei on top of each other. Nuclei with an average DAPI signal per nuclei of between 40 to 90 Arbitrary Units (AU) and a volume between 12000 to 40000 pixels were used. Boxplots plotting the average cFOS and DAPI signal per nuclei from the KCl, TTX, and KCl + Nim conditions were then created, as shown in **Fig. S3.** To test the null hypothesis that the mean amount of cFOS expression within the KCl condition is less than the TTX or KCl + Nim conditions, a Welch's t-test was performed. In both conditions the null hypothesis was rejected (KCl vs TTX pvalue = 9.82216128599208e-75, one-sided) (KCl vs KCl + Nim pvalue = 2.418652397129192e-75, one-sided). In addition, the null hypothesis that the mean amount of cFOS expression is equal in the TTX and KCl + NIM condition could not be rejected using a Welch's t-test (pvalue = 0.6039786961141639, two-sided). Finally, to test if there is a difference among average DAPI intensity per nuclei within the KCl, TTX, and KCl + Nim conditions, an Analysis of Variance (ANOVA) was conducted. The null hypothesis was not rejected, as the ANOVA yielded a pvalue = 0.11396972569868866l; it is likely there are not differences among DAPI staining per nuclei within the KCL, TTX, and NIM conditions.

***Electrophysiology***

Electrophysiology readings were recorded on a Maestro Pro multiwell electrode array (Axion Biosystems) (MEA) on 48 well MEA plates. Each well of the MEA plate well was coated with PLO / laminin as described in *Cell Culture Plate Coating*, with volumes scaled for well size. To increase humidity, 8 mL of Cell Culture Grade Water (Corning, 25-055-CM) was added to on-plate reservoirs. On DIV0, cells were plated at a density of 78,000 cells per $cm^2$. A full media change was performed on DIV1, and half media changes were performed each week from DIV7 to DIV35. Electrophysiology measurements were recorded in the same wells at DIV35 before and after 10 mM KCl stimulation. 10 mM KCl stimulation was performed with a half media change of ACSF with 20 mM KCl using the recipe described in ***Potassium Chloride Stimulation of Neurons***. A minimum of 5 spikes per minute was required as a threshold for an electrode to be considered active. The Axion Biosystems Neural Metrics tool was used to perform batch processing of electrical activity data from the recordings. MEA raster plots were generated before and after stimulation for a representative well, depicting individual spike activity, burst spike activity, and network bursts over time. Burst parameters for analysis were set to require a minimum of 5 spikes with a maximum inter-spike interval (ISI) of 100 ms. Network burst frequency and mean firing rate were also calculated for each timepoint and compared over time using the Neural metrics tool. For network bursts, thresholds were a minimum of 50 spikes with a maximum ISI of 100 ms, as well as a minimum of 35% active electrodes and no burst exclusion. Mean firing rate was calculated in hertz. Summary statistics for a representative well, as well as individual electrodes of said well, before and after 10 mM KCl treatment at DIV35 can be found in **Table S3**. To plot the spikes/bin before and after 10 mM KCl treatment seen in **Fig. S2E,** the spiking data found on sheet three in **Table S3** was converted from seconds to milliseconds, binned in 25 millisecond intervals, and plotted as a histogram.

***Western Blot***

To prepare protein extracts, neurons were collected and washed with cold PBS before performing cell lysis in RIPA buffer (Sigma-Aldrich, R0278) (25 mM Tris-HCl [pH 7.6], 150 mM NaCl, 1% Nonidet P-40, 1% sodium deoxycholate, and 0.1% SDS (Thermo Fisher, BP1311-200)) containing a protease inhibitor cocktail (Sigma-Aldrich, P8340). Neuronal lysates were incubated on ice for 30 minutes. Subsequently, the samples were centrifuged at 21,100 xg for 20 minutes at 4 ºC on a

Sorvall Legend Micro 21R centrifuge, and the supernatant was transferred to a new tube. To determine protein concentration, the Bradford protein assay was used. Before analysis, 2× Laemmli SDS sample buffer (Bio-Rad, 1610737) (125 mM Tris-HCl [pH 6.8], 4% SDS, 20% glycerol, 0.01% bromophenol blue, and 10% 2-mercaptoethanol) was added to the samples. The samples were then denatured at 95 °C for 5 minutes, and equal amounts of total protein were separated using SDS-PAGE.

Proteins were transferred to a polyvinylidene fluoride (PVDF) membrane using Trans-Blot® Turbo™ Mini PVDF Transfer Packs. The membrane was blocked in 5% skim milk diluted in Tris-Buffered Saline with .05% Tween 20 in TBS (TBS-T) (Bio-Rad, 1706435) for 1 hour at room temperature. The PVDF membrane was then incubated with primary antibodies CREB (Cell Signaling Technology, 9197S), pCREB (Cell Signaling Technology, 9198S), and beta-actin (Sigma-Aldrich, A5316) diluted in 5% skimmed milk diluted in TBS-T overnight at 4°C. After primary antibody incubation, the membrane was washed with TBS-T three times for 5 minutes each. Subsequently, the membrane was incubated with 1:4000 Anti-IgG HRP rabbit (Thermo Fisher, HRP31460) and Anti-IgG mouse (Thermo Fisher, HRP31430) secondary antibodies diluted in 5% skim milk in TBS-T for 1 hour at room temperature. TBS-T was used to wash the three times, incubating for 5 minutes per wash.

Following the washes, excess buffer was removed from the membrane, which was then incubated with a chemiluminescence reagent (Surmodics, CHMI-0060-2C) for 1 minute. The excess reagent was removed by dabbing the membrane with a paper towel, after which detection was performed using a ChemiDoc Touch system (Bio-Rad). All membranes were imaged using the Chemi Hi Resolution setting with automatically adjusted exposure times based on band intensity.

Following imaging, all image brightness and contrast levels for a given protein were equivalently adjusted across conditions in FIJI. To quantify protein intensities, the intensity profile for each horizontal band in the blot was plotted in FIJI, and one intensity profile was generated for each of the three proteins tested (pCREB, CREB, β-actin). The area under the intensity profile curve corresponding to each condition was calculated using the Integrated Density tool in FIJI and compared across conditions.

***Nuclear RNA Sequencing***

*Nuclear Isolation*

Nuclei were isolated as described in literature (*124*). Cells were placed on ice, and media was aspirated. Cells were gently washed twice with ice cold PBS to remove dead cells. Half culture volume of PBS was added, and cells were scraped using a silicon cell lifter (Corning, 3008). The cell suspension was transferred to a 15 mL conical using a 1 mL wide bore pipette tip and spun down at 1000 xg for 5 minutes at 4 ºC in a large swinging bucket centrifuge. The supernatant was removed, and the pellet was resuspended in 1 mL NF1 buffer (10 mM Tris-HCl pH8 (Tris Base (Fisher Scientific, BP154), 1 mM EDTA (Sigma, 11873580001), 5 mM MgCl2 (Avantor, MK5958-04), 0.1M Sucrose (Sigma Aldrich, S0389), 0.5% Triton X-100 (Sigma, 93443), 4 units/mL SUPERase RNAse Inhibitor (Thermo, AM2696), 1 tab / 50 mL Protease inhibitor (Roche, 11873580001), DEPC-Treated water (Thermo Fisher, 4387937). Samples were incubated for 3 minutes on ice and then transferred into a 50 mL conical. The tube was washed with an additional 1 mL of NF1 buffer. 18 mL of NF1 buffer was added, and 5 mL Sucrose Cushion Buffer (1.2 M Sucrose, 10 mM Tris-HCl pH8, 3 mM MgCl2, 1 mM DTT (Sigma Aldrich, D0632), 4 units/mL RNAse Inhibitor, 1 tab/50 mL Protease Inhibitor)) was slowly added to the bottom of the tube. The 50 mL conical was then spun at 3900 xg for 30 minutes at 4 ºC, with the brakes set to low (acceleration and deceleration set to 1). The interface and upper aqueous phase were removed using a 25 mL pipette, and the sucrose cushion was gently removed using a p1000 pipette. The pellet was then partially suspended in 8 mL NF1 Buffer by pipette mixing to wash the nuclei and then spun down at 1600 xg for 5 minutes at 4 ºC to pellet the nuclei. The nuclei pellet was resuspended in 1 mL NF1 buffer, passed through a FlowMi strainer (Millipore Sigma, H136800040) and transferred to a 15 mL conical. The tube was washed with an additional 1 mL of NF1 buffer and passed through a FlowMi strainer. Isolated nuclei were counted on a hemocytometer and spun down at 1600 xg for 5 minutes. The supernatant was aspirated, and pellets were stored at -80 ºC after flash freezing in liquid nitrogen.

*RNA Purification and Library Preparation*

Isolated nuclei were resuspended in Trizol (Invitrogen, 10296028) at a concentration of 1 million cells per 350 µL of Trizol. 350 µL of Trizol was then carried forward for RNA isolation. RNA isolation was performed with the Direct-zol total RNA isolation kit (Zymo, R2061) following the manufacturer's protocol. Sample purity and integrity was assessed using an Agilent RNA 6000 Pico reagent kit on a Bioanalyzer 2100 (Agilent Technologies, Santa Clara, CA, USA). To deplete ribosomal RNA from total RNA, the NEBNext rRNA Depletion Kit v2 (NEB, E7405L) was then used, and NEBNext ultra II Directional RNA Library Prep with Sample Purification Beads (NEB, E7765S) was used for library preparation. NEBNext ultra II DNA Library Prep (NEB, E7645S) was used to ligate unique indices to cDNA for multiplexed sequencing. Adaptor ligated products were amplified via polymerase chain reaction (PCR) for 13 cycles. Samples were then size selected with a target of 300 base pairs (bp) and cleaned up with AMPure XP Beads (Beckman Coulter, A63881) (45 µL beads for 50 µL of solution, 0.9X ratio). Pooled samples were sequenced on an Illumina NextSeq1000/2000 Instrument using a 200-cycle (2 x 100) P2 v3 kit.

***Single-Nucleus Methyl-3C sequencing***

*Fixation*

Single Nucleus Methyl 3C-seq3 was performed as described in literature with minor modifications (*64-66*). Cells were fixed with 1% formaldehyde as described in literature (*5, 14, 23, 29, 64, 65, 72, 125*), with some modifications. Cell culture medium was aspirated, and cells were rinsed with PBS. Media was aspirated, and 10 mL of PBS was added to the plate, followed by 1 mL 11% Formaldehyde Solution (50 mM Hepes-KOH, pH7.5 (Hampton Research, HR2-729), 100 mM NaCl (Invitrogen, AM9760G), 1 mM EDTA (Invitrogen, 15575-038), 0.5 mM EGTA (Bioworld, 405200080-1), 37% formaldehyde (Millipore Sigma, F8775)), bringing the final formaldehyde concentration to 1%. Cells were fixed in 1% formaldehyde for 10 minutes, swirling the plates briefly every 2 minutes to ensure equal mixing. To quench the fixation, 580 µL of 2.5 M glycine (Sigma Aldrich, 50046-250G) was added to a final concentration of 125 mM. Samples were incubated for 5 minutes at room temperature, then transferred to 4 ºC for 15 minutes. The media in the plate was then aspirated, and 10 mL PBS was added to the plate. Cells were harvested using a silicon cell lifter, the solution was transferred to a 15 mL conical tube, and cells were pelleted at 1,350 xg for 5 minutes at 4 ºC in a swinging bucket centrifuge. The supernatant was discarded, and cells were rinsed with 1 mL ice cold PBS and spun down again at 1,350 xg for 5 minutes. Cell pellets were stored at -80C after aspirating the supernatant was aspirated and flash freezing in liquid nitrogen.

*Nuclei Isolation*

3C was performed as described in literature (*21, 22, 24, 26, 28, 29, 120, 125*), with some alterations to adapt it to *in vitro* neuronal culture. To isolate nuclei, cell pellets were thawed for 10 minutes on ice, and then resuspended in 1 mL Lysis Buffer (10 mM Tris HCl pH 8.0 (Thermo Fisher, 15568025), 10 mM NaCl, 0.2% Igepal CA630 (Sigma, I8896-50ML), and 1X Protease Inhibitor Cocktail (Sigma, 11873580001) (PIC) in Molecular Biology grade Water (Sigma, W4502)) by adding 50 µL Lysis Buffer, flicking the tube to detach the pellet, adding 950 µL (total volume 1 mL), and pipetting up and down 20 times or until solution appeared homogenous. The sample was then transferred to a 1.5 mL DNA Low Bind Tube (Eppendorf, 022431021) and placed on a rotator (10 *RPM*) at 4 ºC for 20 minutes. The sample was then spun down at 600 xg in a Swinging Bucket Centrifuge (Thermo Fisher, 75004521), the supernatant was aspirated, and the sample was resuspended in fresh ice-cold Lysis Buffer. To isolate nuclei, the sample was transferred to a dounce homogenizer (DWK Life Sciences, 885300-0002), and dounced 120 times with Pestle A. A DAPI stain and fluorescent microscopy was used to confirm successful nuclei isolation into a single-nuclei suspension. If nuclei were not fully isolated, an additional 60 strokes of Pestle A were used.

*Flow Cytometry*

Upon nuclei isolation, samples were then passed through a 35 µm cell strainer cap into a 5 mL polystyrene round bottom tube (Falcon, 352235). Nuclei were stained with 1:1000 Sytox Red Cell Stain (Thermo Fisher, S34859) and sorted on a FACSAria II Sorter in the Flow Cytometry Core Laboratory in the Children's Hospital of Philadelphia (CHOP) Research Institute using Fluorescence Activated Nuclei Sorting (FANS) to identify intact, Sytox+ nuclei. Samples were gated on Forward and Side Scatter Area to identify intact nuclei, Forward Scatter Area and Height to identify singlets, and on 685 nm emission and Side Scatter Area to identify Sytox+ Nuclei (**Fig. S4A).** 200,000 cells were sorted per condition.

*Chromosome Conformation Capture*

Upon isolation of intact nuclei via FANS, samples were spun down for 5 minutes at 600 xg, resuspended in 50 µL 0.5 % sodium dodecyl sulfate (SDS), and incubated at 62C for 10 minutes to permeabilize nuclei. SDS was then quenched via the addition of 145 µL water and 25 µL 10% Triton X-100 (Sigma, 93443), and samples were incubated at 37 ºC for 15 minutes. To digest chromatin, 25 µL of 10X NEBuffer r3.1 (NEB, B6003S), 100 units of NlaIII (NEB, R0125S), and 100 units of MboI (NEB, R0147M) were added, and samples were incubated for 16 hours at 37 ºC with rotation at 850 RPM. Samples were then incubated at 65 ºC for 20 minutes to inactivate the restriction enzymes, and restriction fragment overhangs were blunted by adding 1.5 µL each of 10 mM dATP, dCTP, dTTP, and dGTP (NEB 10297018), as well as 8 µL of 5 U/µL DNA Polymerase I Large (Klenow) fragment (NEB, M0210). Samples were incubated at 37C for 45 minutes with rotation at 850 RPM. To proximity ligate blunt ends, 900 µL of Ligation Master Mix (669 µL water, 120 µL 10X NEB T4 DNA ligase buffer (NEB, B0202), 100 µL 10% Triton X-100, 6 µL 20 mg/mL Bovine Serum Albumin (NEB, B9000S) and 5 µL 400 U/µL T4 DNA Ligase (NEB, M0202)) was added, and samples were incubated with slow rotation (10 RPM) for 4 hours at room temperature.

*FANS Sorting to Single Nuclei*

Cells were sorted into single wells as described in literature with minor modifications (*64-66*). While samples were ligating, three 384 well plates (Thermo Fisher, 4483351) per condition were prepared by adding 1.3 μL Digestion Mixture (0.650 uL M-Digestion Buffer (Zymo, D5021-9), 0.043 μL Proteinase K (Zymo, D3001-2-20), 0.607 uL water) per well. Samples were spun down at 600 xg for 5 minutes at 4 ºC, resuspended in 800 μL 1% BSA in PBS, stained with 1:1000 Sytox red, and filtered into a polystyrene round bottom tube through a 35 μm cell strainer. Samples were sorted into one Sytox+ nuclei per well using a FACSAria II Sorter in the Flow Cytometry Core Laboratory in the CHOP Research Institute using FANS (**Fig. S4B)**. After sorting, plates were sealed using MicroAmp Optical Adhesive Film (Fisher, 4360954), spun down for 2 minutes at 1000xg at 4 ºC, and incubated for 20 minutes at 50 ºC to degrade crosslinks and proteins. Samples were spun down at 2000 xg for 2 minutes at 4 ºC and stored at -20 ºC.

*Bisulfite Conversion and Cleanup*

To prepare the CT Conversion Reagent, 7.9 mL Solubilization Buffer (Zymo, D5021-7) and 3 mL Dilution Buffer (Zymo, D5006-2) were added to a CT Conversion Reagent bottle (Zymo, D5003-1) and mixed at room temperature for 10 minutes. 1.6 mL of Reaction Buffer (Zymo, D5021-8) was then added, and the resulting solution was mixed at room temperature for an additional 5 minutes. To convert unmethylated cytosines to uracils, 6.2 μL CT Conversion Reagent was added to each well, sample was mixed 8 times, the 384 well plate was sealed, and samples were spun down at 1000xg for 20 seconds at 4 ºC. Samples were then incubated at 98 ºC for 8 minutes, followed by 64 ºC for 3.5 hours, and then held at 4 ºC for up to 20 hours. To clean up samples after the bisulfite conversion, samples were then spun down for 2 minutes at 2000xg at 4 ºC, followed by the addition of 20 μL M-Binding Buffer (Zymo, D5040-3) to each well. Samples were mixed 8 times and transferred to a 384 Well DNA Binding Plate (Zymo, C2012). To load DNA into the spin column, samples were spun down for 5 minutes at 4000xg at room temperature. Samples were then washed with 100 μL of M-Wash Buffer (D5040-4) and spun down for 5 minutes at 4000xg at room temperature. To desulphonate the bisulfite-converted DNA, 50 μL M-Desulphonation Buffer (Zymo, D5040-5) was added to each well. Plates were incubated for 15 minutes at room temperature and then spun down for 5 minutes at 4000 xg at room temperature. Plates were then washed twice with 100 μL M-Wash Buffer, spinning down for 5 minutes at 4000xg each time. To elute DNA from the spin column, 4.4 μL elution buffer was added to each well, samples were incubated at room temperature for 5 minutes and then spun down into a new 384 well plate for 5 minutes at 4000xg at room temperature.

*Random Priming and Inactivation of Free Primers and dNTPs*

To individually barcode each well, 1.1 μL of 2.5 μM of a unique random priming barcode (*64-66, 126*) was added to each well of the 384 well plate. Samples were sealed, spun down at 1000xg for 2 minutes to mix, denatured at 98C for 3 minutes, immediately placed on ice to prevent rehybridization, and spun down again at 2000xg for 1 minute at 4 ºC. To anneal the random priming barcodes, 5 μL of the Random Priming Master Mix (1.025 μL 10X Blue Buffer (Qiagen, B0110L), 0.025 μL 50 U/μL Klenow Exo- (Qiagen, P7010-HC-L), 0.5 μL 10 mM dNTP (NEB, N0447L), 3.45 μL water) was added to each well. Plates were sealed and spun down at 2000xg for 1 minute at 4 ºC and then incubated with the following program: 4 ºC for 5 minutes; 25 ºC for 5 minutes, 37 ºC for 60 minutes, hold at 4 ºC. Samples were spun down at 2000xg for 1 minute at 4 ºC. To deactivate free primers and dNTPs, 1.5 μL of the Inactivation Master Mix (0.2 μL 10X Blue Buffer, 0.1 μL 20U/μL Exonuclease I (Qiagen, X8010L), 0.05 μL 1U/μL rSAP (NEB, M0371L), 1.15 μL water) was added to each well. Plates were sealed, spun down at 2000xg for 1 minute at 4 ºC, incubated for 30 minutes at 37 ºC, and spun down at 2000xg for 1 minute at 4 ºC.

If multiple plates were being processed in parallel, plates were placed at -20 ºC avoid excess exonuclease activity.

*3' Hybridization and PCR amplification*

To compress barcoded samples from 384 well plates down to 6 PCR strip tubes, prior to the addition of the 3' adaptor, 8 wells from the 384 well plate were combined into one well of a 96 well plate, with a total volume of 92 µL per well of the 96 well plate. To cleanup and size select DNA, 73.6 µL of Agencourt AMPure XP beads (Beckman Coulter, A63881) was added to each well and pipetted up and down 20 times to mix. Samples were incubated for 8 minutes and placed on a magnet for 5 minutes. The supernatant was removed, and 180 µL of 80% ethanol in water was added to each well. Samples were incubated for 30 seconds, the ethanol was removed, and the wash step was repeated. Samples were removed from beads, incubated for 2 minutes to allow the ethanol to evaporate, and resuspended in 10 µL elution buffer per well. Plates were sealed and gently vortexed to remove beads from the side of the well, and spun down for 5 seconds at 150xg. Samples were incubated for 5 minutes at room temperature, spun down for 5 seconds at 100xg, and placed on a magnet for 2 minutes. Eight wells of the 96 well plate were then condensed down into 1 PCR strip tube (6 PCR strip tubes total per 384 well plate), with a final volume of 80 µL. 64 µL AMPure XP beads were added to each tube, and samples were pipetted up and down 20 times to mix. Samples were incubated at room temperature for 8 minutes, followed by incubation on a magnet for 5 minutes, and the supernatant was discarded. Samples were washed twice with 180 µL 80% ethanol, the supernatant was removed, and samples were air-dried for two minutes to allow any excess ethanol to evaporate. Samples were resuspended in 10 µL elution buffer, incubated for 5 minutes, and placed on a magnet for 2 minutes. The supernatant containing the barcoded DNA was then transferred to a fresh PCR strip tube.

To anneal the 3' adaptor, samples were denatured at 98 ºC for 3 minutes, immediately placed on ice for 2 minutes and briefly spun down. 10.5 µL of the Adaptase Master Mix (4.25 µL Low EDTA TE (Swift Biosciences, 90296), 2 µL Buffer G1, 2 µL Buffer G2, 1.25 µL Buffer G3, 0.5 µL Enzyme G4, 0.5 µL Enzyme G5 (all products from xGen Adaptase Module (IDT, 10009826)) was added to each tube, tubes were flicked to mix and briefly spun down. Tubes were incubated on a thermalcycler for 30 minutes at 37 ºC, followed by 2 minutes at 95 ºC, and held at 4 ºC.

To PCR amplify the DNA, 30.3 µL of the Indexed PCR Master Mix (25 µL 2X KAPA HiFi Hotstart ReadyMix (Roche, KK2602), 0.3 µL 100 µM Custom P5L indexing primer (IDT) (*64, 65, 126*) 0.5 µL 100 µM Custom P7L indexing primer (IDT) (*64-66, 126*) (4.5 µL water) was added to the tube. Samples were briefly vortexed and spun down. Samples were then PCR amplified using the following program: i) 95 ºC for 2 minutes; ii) 98 ºC for 30 seconds; iii) 98 ºC for 30 seconds; iv) 64 ºC for 15 seconds; v) 72 ºC for 30 seconds; repeat steps iii-v for a total of 15 cycles; 72 ºC for 5 minutes; hold at 4 ºC. Tubes were spun down, 40.6 µL of AMPure XP beads were added, and samples were pipetted up and down. Samples were incubated for 8 minutes, placed on a magnet for 5 minutes, and the supernatant was discarded. Samples were washed twice with 180 µL of 80% ethanol, the supernatant was discarded, and samples removed from the magnet and allowed to airdry to evaporate any residual ethanol. Samples were resuspended in 20 µL elution buffer, incubated for 5 minutes at room temperature, and put on a magnet for 2 minutes. The supernatant from each of the 6 PCR strip tubes containing the PCR amplified DNA was then combined into one DNA low bind tube. 96 µL of AMPure XP beads were added to the sample and pipetted up and down 20 times. Samples were incubated at room temperature for 8 minutes, placed on a magnet for 5 minutes, and the supernatant was discarded. Samples were washed twice with 180 µL of 80% ethanol, the supernatant was discarded, and samples were allowed to airdry. Samples were resuspended in 20 µL elution buffer, incubated for 5 minutes, and placed on a magnet for 2 minutes. The 20 µL supernatant containing the final libraries was then transferred to a new tube and stored at -20 ºC. Sample concentration was assessed using a Qubit fluorometer, and the size distribution of the DNA was assessed using an Agilent DNA High Sensitivity reagent kit on a Bioanalyzer 2100 (Agilent Technologies, Santa Clara, CA, USA). Pooled samples were sequenced on NextSeq 1000/2000 and Novaseq 6000.

***CUT&Tag Direct***

*Collection and Fixation*

Neuronal nuclei were isolated using a sucrose density gradient as described above in the *Nuclear Isolation* section of ***Nuclear RNA Seq*** with minor modifications. After the sucrose cushion step, nuclei were washed in 10 mL 0.1% BSA in PBS, supplemented with 1X Protease inhibitor and RNase inhibitor, and centrifuged for 5 minutes at 1600 xg at 4 ºC. The supernatant was aspirated, and nuclei were resuspended in 950 µL 0.1% formaldehyde for 2 minutes to lightly fix the nuclei. The fixation was quenched with 66 µL 1.25 M glycine. The samples were spun down at 1600 xg for 5 minutes at 4 ºC, resuspended in Synthafreeze, and slow-frozen at -80 ºC.

*Primary Antibody Incubation*

Nuclei were processed for CUT&Tag Direct as described in literature (*68, 69*), with minor modifications. Briefly, frozen, isolated $i^3$N neuronal nuclei were thawed at room temperature and bound to BioMag®Plus Concanavalin A beads (Polysciences, 86057-3) and incubated bead-bound nuclei in primary antibody overnight at 4 ºC with rotation. All primary antibodies (H3K27ac (abcam, ab4729), CTCF (Epicypher, 13-2014), IgG, (MilliporeSigma, I8140-10MG)) were used at a 1:100 dilution in antibody binding buffer (20 mM HEPES KOH pH 7.5 (ThermoFisher, J607120-AK), 150 mM NaCl, 0.5 mM spermidine (MilliporeSigma, S2501-5G), 0.1% digitonin (Calbiochem, 300410-1GM), 2 mM EDTA).

*Secondary Antibody Incubation*

The following day, primary antibody solution was replaced with a secondary antibody in antibody binding buffer, 1:100 of secondary antibody, Guinea Pig anti-Rabbit IgG (Antibodies Online, ABIN10196). Samples were incubated at room temperature for 1 hour with gentle agitation. The secondary antibody solution was then removed, and samples were washed twice with Digi-Wash buffer (20 mM HEPES KOH pH 7.5, 150 mM NaCl, 0.5 mM spermidine, 0.1% digitonin). Samples were resuspended in 300-Digi-Wash buffer and CUTANA™ pAG-Tn5 for CUT&Tag (Epicypher, 15-1017) at 1:20 dilution per manufacturer's recommendation. After incubation at room temperature for 1 hour with gentle agitation, samples were washed with 300-Digi-Wash buffer three times. For tagmentation, samples were resuspended in 300-Digi-Wash with 10 mM $MgCl_2$ and incubated on a thermocycler at 37 ºC for 1 hour. Samples were then washed with Digi-Wash buffer and resuspended in 5 µL SDS release buffer (0.01 M TAPS pH 8.5 (Boston BioProducts, BB-2375), 0.1% SDS in water). Samples were then incubated on a thermocycler at 58 ºC for one hour, followed by the addition of 15 µL SDS quench buffer (0.67% Triton X-100 in water) to quench the reaction.

*Library Amplification*
To amplify the samples, 25 µL of NEBNext® High-Fidelity 2X PCR Master Mix (NEB, M0541S) and 2 µL of i5 and i7 Nextera Index Primers (Illumina, FC-131-1001) were added. Samples were amplified via PCR using the following settings: 58 ºC for 5 minutes, 72 ºC for 5 minutes, 98 ºC for 45 minutes, then 16 cycles of 98 ºC for 15 seconds and 60 ºC for 10 seconds, followed by 72 ºC for 1 minute. PCR libraries were cleaned up with 1.1X AMPure XP beads (Beckman, A63881). Pooled samples were sequenced on a NextSeq 1000/2000.

***ChIP-seq***
*Fixation and Antibody Coupling*
Cells were single fixed in 1% formaldehyde as described in the *Fixation* section of ***Single Nucleus Methyl 3C-Seq.*** CTCF ChIP-seq was performed as previously described (*5, 14, 16, 22, 23, 26, 28*) with minor optimizations for efficient neuronal membrane lysis and nuclei number normalization prior to nuclear lysis. The day before cell lysis, CTCF antibody was coupled to agarose beads overnight. For each sample, 10 µL of CTCF antibody (Sigma-Aldrich, 07-729), 20 µL Protein A agarose beads (Thermo Scientific, 20333), and 20 µL Protein G agarose beads (Thermo Scientific, 20398) were added to 1 mL of cold PBS without calcium or magnesium in a 1.5 mL LoBind microcentrifuge tube and rotated at 10 rpm at 4 °C overnight.

*Cell and Nuclear Lysis*
Each cell pellet was thawed on ice for 15 min, resuspended in 5 mL cold ChIP Cell Lysis Buffer (approximately 1 mL per million cells; 10 mM Tris pH 8.0, 10 mM NaCl , 0.2 % (v/v) IGEPAL CA-630, 1X Protease Inhibitor Cocktail, 1 mM phenylmethanesulfonyl fluoride (PMSF; Sigma, 93482) in molecular biology grade water), and incubated on ice for 15 min. The cell lysate was centrifuged at 600 xg for 5 min at 4 °C in a swinging bucket centrifuge. After removing supernatant, the nuclei pellet was resuspended in 1 mL fresh ChIP Cell Lysis Buffer and incubated on ice for 15 min. The cell lysate was further homogenized on ice using a dounce homogenizer and 30 strokes with pestle A. After confirming successful single-nuclei isolation via fluorescence microscopy using a DAPI stain, nuclei were counted using an automated cell counter. 2.3 million nuclei per sample were centrifuged at 600 xg for 5 min at 4 °C in a swinging bucket centrifuge, resuspended in 500 µL cold Nuclear Lysis Buffer (50 mM Tris pH 8.0, 10 mM EDTA, 1% (w/v) sodium dodecyl sulfate (SDS; Invitrogen, 15553027), 1X PIC, 1 mM PMSF in molecular biology grade water), and incubated on ice for 20 min.

*Sonication and Pre-clearing*
The nuclei lysate was transferred to a polystyrene sonication tube (Active Motif, 53071) and 300 µL of cold Dilution Buffer (20 mM Tris pH 8.0, 2 mM EDTA, 150 mM NaCl, 1% (v/v) TritonX-100, 0.01% (w/v) SDS, 1X PIC, 1 mM PMSF in molecular biology grade water) was added to each tube. Chromatin was sheared at 4 °C using the following parameters on a Qsonica Q800R3 (1 hour duration, 30 sec on/30 sec off pulses, 100% amplitude). Sheared chromatin was centrifuged at 16,200 xg for 5 min at 4 °C to pellet nuclei debris, and the supernatant containing chromatin (around 800 µL) was added to a pre-chilled 5 mL LoBind tube (Eppendorf, 0030108310) containing 3.7 mL Dilution Buffer, 500 µL Nuclear Lysis Buffer, 87.5 µL each of Protein A and Protein G agarose beads, and 50 µg of IgG (Sigma, I8140) for pre-clearing. Each pre-clearing sample was rotated at 10 rpm at 4 °C for 2 hours.

*Immunoprecipitation*

Protein A/G agarose beads coupled to the CTCF antibody (post 12 hr overnight incubation) were centrifuged at 400 xg at 4 °C for 5 min, and the supernatant was removed. Pre-cleared samples were centrifuged at 875 xg at 4 °C for 5 min in a swinging bucket centrifuge to pellet the agarose beads. The supernatant containing pre-cleared chromatin was transferred to a new pre-chilled 5 mL LoBind tube. 50 μL of the pre-cleared chromatin was saved as the "Input" sample and stored at -20 °C until DNA purification the next day. Then, 100 μL of the pre-cleared chromatin was used to wash and transfer the CTCF antibody-coupled agarose bead to the 5 mL LoBind tube. Each immunoprecipitation sample was rotated at 10 rpm at 4 °C for 16 hours.

*Bead Washing and Elution*

Immunoprecipitation (IP) reactions were centrifuged at 3,485 xg for 5 min at 4 °C using a swinging bucket centrifuge to pellet the agarose beads, and the supernatant was removed. 1 mL of cold Wash Buffer I (20 mM Tris pH 8.0, 2 mM EDTA, 50 mM NaCl, 1% (v/v) TritonX-100, 0.1% (w/v) SDS in water) was added to each sample to resuspend the beads, and the entire sample was transferred to a new 1.5 mL LoBind tube. After vortexing gently (speed 7) for 5 seconds, samples were centrifuged at 5,400 xg for 2 min at 4 °C. The supernatant was removed and beads were washed twice with 1 mL of cold High Salt Buffer (20 mM Tris pH 8.0, 2 mM EDTA, 500 mM NaCl, 1% (v/v) TritonX-100, 0.01% (w/v) SDS in water) and once with cold Wash Buffer II (10 mM Tris pH 8.0, 1 mM EDTA, 250 mM lithium chloride (Sigma, L9650), 1% (w/v) IGEPAL CA-630, 1% (w/v) sodium deoxycholate (Sigma, D6750) in water) in the same sample tube. After the Wash Buffer II step, supernatant was removed, 1 mL of cold Tris-EDTA, pH 8.0 (TE; Fisher BP2473100) was added to the beads, and the entire solution was transferred to a new 1.5 mL LoBind tube to minimize the transfer of lithium chloride. After one more wash in cold Tris-EDTA (for a total of two washes), sample tubes were moved to room temperature, and 103 μL of room temperature Elution Buffer (0.084 g sodium bicarbonate (Fisher, S233-500) dissolved in 10 mL 1% (w/v) SDS solution in water) was added to each sample. Samples were vortexed on high (speed 10) for 5 seconds and spun down at room temperature for 2 min at 5,400 xg. 100 μL of the supernatant containing eluted chromatin was transferred to a new 1.5 mL LoBind tube. Another round of elution was performed with 100 μL of fresh Elution Buffer, and 100 μL of the supernatant was combined with the previous elution for a total volume of 200 μL eluted chromatin.

*DNA Purification and Library Preparation*

150 μL of TE was added to Input samples to bring the total volume to 200 μL. 2 μg RNase A (Thermo Scientific, EN0531) and 12 μL 5 M NaCl were added to each IP and Input sample, followed by incubation at 65 °C for 1 hour. Then, 3 μL of Proteinase K (NEB, P8107S) was added to each IP and Input sample, followed by incubation at 65 °C for 16 hours. Each sample was brought up to 350 μL with TE, and Phenol:Chloroform:Isoamyl Alcohol (PCIA) DNA extraction was performed. Specifically, 350 μL of PCIA (Thermo Fisher, 15593031) was added to each sample, and each sample was vortexed at maximum speed (speed 10) for 30 sec. The 700 μL mix was transferred to a 2 mL phase lock gel tube (Quantabio, 2302830) and centrifuged at 17,000 xg for 10 min at room temperature. Approximately 350 μL of the aqueous phase was transferred to a new 1.5 mL LoBind tube. 50 μL of fresh TE was added to the phase lock tube, and another round of phase separation was performed as above to maximize yield. 40 μL of 3 M sodium acetate (pH 5.2, Thermo Scientific, R1181) and 30 μg glycogen (Invitrogen, AM9510) were added to 400 μL

of extracted chromatin sample and mixed well by inversion, followed by incubation at -80 °C overnight.

Samples were centrifuged at 21,000 xg for 30 min at 4 °C to pellet the chromatin, and the pellets were washed once with 1 mL cold 80% ethanol. After centrifugation at 16,000 xg for 15 min at 4 °C, supernatant was removed, and pellets were dried at 37 °C for 5 min to remove residual alcohol. Each pellet was resuspended in 22 µL of pre-warmed TE and incubated at 37 °C for 10 min to dissolve the DNA. 1 µg of each IP and Input sample were prepared for Illumina sequencing using NEBNext Ultra II DNA Library Prep Kit for Illumina (NEB, E7645) and NEBNext Multiplex Oligos for Illumina (NEB, E7335) following manufacturer instructions. IP and Input libraries were sequenced with single-end sequencing on a NextSeq™ 1000/2000 P2 XLEAP-SBS™ flow cell (100 Cycles; Illumina, 20100987) for an average of 50 million sequenced reads per library.

***Single-nucleus RNA seq***

Three experimental conditions were collected for single-nucleus RNA seq (snRNAseq): *KCl DIV35 Collect DIV40, TTX DIV35 KCl DIV40,* and *KCl DIV35 KCl DIV40.* Neurons were differentiated as described in ***Differentiation of hiPSCs into Functional Cortical Neurons.*** At DIV34, all three conditions received 24 hours of 1 µM TTX. At DIV35, the *KCl DIV35 Collect DIV40* and *KCl DIV35 KCl DIV40* conditions were stimulated for one hour in High KCl ACSF and returned to basal media as described in ***Potassium Chloride Stimulation of Neurons.*** The *TTX DIV35 KCl DIV40* condition did not receive KCl and was returned to basal media after 24 hours of TTX treatment. At DIV40, *KCl DIV35 Collect DIV40* was collected, and *TTX DIV35 KCl DIV40* and *KCl DIV35 KCl DIV40* were stimulated for 1 hour in high KCl, no Mg ACSF and then collected. To collect cells, plates were washed once in cold PBS, cells were scraped in cold PBS and then centrifuged for 3 minutes at 250 xg. Cell pellets were resuspended in Synthafreeze (Gibco, A1254201), transferred to a cryotube, and slow-froze to -80 ºC. Samples were recovered by thawing at room temperature and extracting nuclei via sucrose gradient centrifugation as described in ***Nuclear RNA Sequencing:*** *Nuclear Isolation.* Upon nuclei isolation, nuclei were resuspended in FANS Buffer (0.2% BSA in PBS, supplemented with protease inhibitor and RNAse inhibitor (Invitrogen, AM2696)), stained with 1:1000 DAPI (Sigma-Aldrich, D9542-1MG) and sorted on an AurorsaCS-2 (Cytek) sorter with a 70 µm nozzle. High-quality nuclei populations were identified by sorting first on forward-scatter area and side scatter area to remove debris, then forward-scatter area and forward-scatter heigh to identify single nuclei, then side-scatter area and UV7 area to identify DAPI positive nuclei using SpectroFlo CS (Cytek) software. High quality nuclei fell into a low variability positive sub-population within the UV7 area measurement. After sorting, nuclei were counted and diluted to 700 nuclei/µL in FANS buffer. To generate droplets containing a single nuclei per droplet, the Chromium Next GEM Chip G Single Cell Kit (10X Genomics, PN-1000120) on the Chromium Controller (10X Genomics, PN-120270) was used, targeting the recovery of 10,000 nuclei per condition. Single nucleus RNA seq was then performed using the Chromium Next GEM Single Cell 3' Reagent Kit v3.1 (10X Genomics, PN-1000268) following the manufacturers protocol. Samples were indexed using the Dual Index Kit TT Set A (10X Genomics, PN-1000127), and libraries were amplified using 15 PCR cycles. Samples were sequenced on the NextSeq™ 1000/2000 P2 flow cell (100 cycles; Illumina, 20046811), setting Read 1 to 28 cycles and Read 2 to 90 cycles.

***Nuclear RNA Seq Computational Analysis***

First, the NCBI Refseq GTF and FASTA files corresponding to the hg38 genome build were downloaded on February 6th, 2021, from the UCSC Genome Browser. (https://hgdownload.soe.ucsc.edu/goldenPath/hg38/bigZips/). Then the GTF and FASTA files were modified to include the feature and the genetic sequences corresponding to the mouse *Ngn2* transgene and the rtTA transgene (*57, 58*), respectively. To align reads to both exons and introns, the HISAT2 software package (version 2.2.0) was used (*127*). The reference index was generated using hisat2-build using the parameters --exon and --ss. These parameters correspond to exon and splice site lists respectively. The exon and splice site lists were generated using the hisat2_extract_exons.py and hisat2_extract_splice_sites.py scripts from the HISAT2 package. Once the index was created, HISAT2 was run with default settings. The resulting SAM files were converted to bam and then filtered to remove unmapped reads using samtools (version 1.3.1) (*128*). To account for read depth differences, all filtered bam files were downsampled to 63.3 million reads using the reformat.sh script from the BBMap package (version 39.01) with the following parameters: samplereadstarget=66300000 and sampleseed=0. The StringTie software package (version 2.2.1) (*129*) was used to quantify transcript counts. Downsampled bam files and the modified GTF file were used as inputs and the software was run with default parameters. To extract estimated transcript counts for downstream analyses, the prepDE.py script from the StringTie package was run with 116 as the read length parameter (-1). To reduce redundancy, estimated counts of transcripts whose transcriptional start site (TSS) positions are within 1 base pair (bp) of each other were summed together into transcriptional units (TUs). Then, TUs without any conditions where both replicates have at least 25 counts were removed. Throughout this document, when the language ‘gene expression’ or ‘gene counts’ is used when referring to bulk nuclear RNA seq, this refers to TU expression or counts, respectively. After filtering, the resulting counts matrix was quantile normalized using the second replicate of the 0-hour condition after KCl treatment as the reference. The reference was selected based on principal component analysis with the goal of reducing variance between both replicates across all conditions (**Table S4**).

To identify differentially expressed transcriptional units, the DESeq2 pipeline (version 1.36.0) (*130*) was ran with modifications. Rather than running the default pipeline, all size factors were set to 1 to avoid double normalization. Then dispersions were estimated using the estimateDispersions function, followed by statistical testing using the nbinomWaldTest function. The first set of comparisons were 2 hours post-KCl vs. TTX and 5 hours post-KCl vs. TTX. Significant genes were selected based on a “padj” value less than 0.05. This value is an adjusted p-value after Benjamini Hochberg multiple testing correction.

***CUT&Tag Analysis***

*Pre-processing:* Paired end fastq reads from H3K27ac and IgG CUT&Tag libraries were merged across all runs and trimmed using cutadapt (version 1.18) (*131*) using -a CTGTCTCTTATACACATCT -A CTGTCTCTTATACACATCT --nextseq-trim 20 -q 20,20 --minimum-length 15 as arguments. Trimmed reads were then aligned to the hg38 genome build using bowtie2 (version 2.4.1) (*132*) using –local, --very-sensitive-local, --no-mixed, --no-discordant, --phred33, -I 10, and -X 700 as arguments. Aligned bam files were sorted using samtools (version 1.9) using the -n and -l 9 arguments and then filtered for unmapped, chrM, and duplicate reads (version 1.15.1) (*128*). Additionally, reads that overlapped the hg38 unified blacklist regions bed file from ENCODE (https://www.encodeproject.org/files/ENCFF356LFX/) were removed using bedtools (version 2.30.0) (*133*). Filtered reads were then downsampled with samtools view -s where the seed was 42. Peaks were called using macs2 (version 2.2.9.1) Signal per million reads normalized bedgraph files were generated from macs2 using the -B and --SPMR arguments (*134*).

H3K27ac bedgraphs were input normalized with respect to IgG. Because the genomic coordinates outputted by macs2 are not regular intervals, entire chromosomes were re-binned at 10 base pair resolution using the "max" value from the pyBigWig stats function (version 0.3.18) (*135*). After rebinning, IgG signal was subtracted from the H3K27ac signal for each condition. After IgG normalization, the resulting H3K27ac bedgraphs were peak called directly using macs2 bdgpeakcall (version 2.2.9.1) (*134*). The signal cutoff parameter -c was set to 1 with otherwise default settings.

After peak calling, peak windows within 400 bp distance were merged using bedtools merge -d 400 (version 2.30.0) (*133*). Pairwise differential classification of merged peak windows for each treatment was conducted only for H3K27ac libraries using a similar procedure from (*70, 72*) for ChIP-seq data but with modifications. First, merged peaks from the two conditions being compared were concatenated together and then overlapping windows were merged using bedtools merge (version 2.30.0) (*133*). Any peaks overlapping exons, UTRs, or any promoter region spanning 200 bp upstream to 200 bp downstream of merged TSSs (see ***Nuclear RNA Seq Computational Analysis)*** were filtered out using bedtools intersect (version 2.30.0) (*133*). Then, for each non-coding peak (i.e. not overlapping a promoter, exon, or UTR), a sum value across all overlapping genomic intervals were computed from bigWig values obtained using pyBigWig (version 0.3.18) (*135*). Each of these sum values were then divided by 400, corresponding to the total window size in base pairs. Next, a fold-change $U_n^{AB}$ between signal per base pair values, $h$, for condition A and B was computed for the nth peak (**Equation 1)**.

$$U_n^{AB} = \frac{h_n^B}{h_n^A} \qquad \textbf{(1)}$$

Differential classification of each peak window was carried out based on the 25th percentile of signal per base pair values across all peak windows, G, and fold-change, U. A peak was considered B-specific when $h_n^B \geq G$ and $U_n^{AB} \geq 2$ or $h_n^A = 0$ and $U_n^{AB} \geq 2$. A peak was considered A-specific if $h_n^A \geq G$ and $U_n^{AB} \leq \frac{1}{2}$. A peak was considered invariant when $h_n^B \geq G, h_n^A \geq G$, and $\frac{1}{2} \leq U_n^{AB} \leq 2$. Any peaks that did not meet these conditions were excluded from further analysis. Noncoding peaks identified as TTX-specific were defined putative activity-decomissioned enhancers, and noncoding peaks defined as KCl-specifc were defined as putative activity-induced enhancers (**Tables S5-S6)**.

##### *ChIP-seq Analysis*

*Pre-processing:* Paired end fastq reads from CTCF IP and Input ChIP-seq libraries were concatenated across all runs and reads were aligned with Bowtie (version 0.12.7) (*136*) using the --tryhard and -m 2 flags. After alignment, unmapped and duplicate reads were removed using samtools (version 1.3.1) (*128*). Then both CTCF and Input libraries were downsampled to 36 million reads, which is the rounded number of paired reads of the smallest library using a custom script. Then to identify genomic locations of CTCF peaks, the macs2 callpeak (version 2.2.9.1)

(*134*) function was ran with the -p 1e-4, -B and –SPMR flags and downsampled BAM files were passed in. After obtaining narrowPeaks, rows with a fold-change "signalValue" less than 5 were removed. Filtered peaks across TTX, 2 hours post-KCl, and 5 days post-KCl were then merged together using bedtools merge (v2.26.0) (*133*) (**Table S7**).

Macs2 signal per million reads normalized bedgraphs across TTX, 2 hours post-KCl, and 5 days post-KCl were scaled together to address signal bias that remained unaccounted for after downsampling and running macs2. Scale factors were computed for each condition and chromosome. For a given chromosome, a list of max signals across all peak windows from the merged peaks list was computed for each condition. Then, the 75$^{th}$ percentile of the list max signals was obtained for each condition. The final scale factor for each condition and chromosome was the 75th percentile divided by the 75$^{th}$ percentile of the TTX condition. After obtaining scalar factors, each value in the bedgraph was normalized by dividing that value with the appropriate scalar with respect to condition and chromosome.

***<u>Single-Nucleus methyl 3C seq Computational Analysis</u>***

*Running the YAP Pipeline*

Paired end fastq files were merged across all runs for each 384-well plate. To demultiplex the merged fastq files into single cell level fastq files, the 'yap demultiplex' command from the YAP pipeline was run (https://hq-1.gitbook.io/mc/inhouse-demultiplex)(*64, 65*). The same virtual environment and similar steps detailed in the supplemental text of Xu and Chandradoss et al. 2025 (*126*) were used. In brief, 'm3c-V2.human.mapping_config.ini' was downloaded from the YAP GitHub page (https://hq-1.gitbook.io/mc/prepare/prepare-mapping-config) with default settings. A default reference FASTA, bismark reference, and chrom sizes file corresponding to the hg38 genome build was specified. Demultiplexing generated six subfolders for each plate containing single cell fastq files and a snakemake command for generating BAM, ALLC, and 3C valid pairs files. The snakemake commands were modified to run in parallel on a LSF high performance cluster with the bsub command. After snakemake commands finished running, all resulting 'MappingSummary.csv.gz' files were merged and additional columns such as 'PlateID' (e.g. TTX_Plate1) and 'SampleID' (e.g. TTX) were added.

*Computing Quality Control Metrics*

For each cell, additional mapping-related metrics were computed: 'TotalReads', the sum of 'R1InputReads' and 'R2InputReads', 'TotalMapped', the sum of 'R1UniqueMappedReads' and 'R2UniqueMappedReads', 'TotalDedupped' (also referred to as final reads), the sum of 'R1DeduppedReads' and 'R2DeduppedReads', 'MappingRate', the ratio of 'TotalMapped' and 'TotalReads', and 'DeduppedRate', the ratio of 'TotalDedupped' and 'TotalDedupped'. Then, 'CisTransRatio', the sum of 'CisShortContact' and 'CisLongContact' divided by 'TransContact', was computed.

*Filtering Poor-Quality Cells*

Poor-quality cells were identified and removed based on 5 established quality control metrics such as 'MappingRate', 'TotalDedupped', 'mCHFrac', 'mCCCFrac', and 'mCGFrac' (*64, 65*). 'mCHFrac' is the proportion of reads indicating non-CpG methylation to the total reads covering cytosines in the non-CpG context. 'mCCCFrac' is the proportion of reads supporting methylation of cytosines to the total reads covering cytosines in the CCC context. 'mCGFrac' is the proportion of reads supporting CpG methylation to the total reads covering cytosines in the CpG context. Based on the density histogram for the mapping rate across all cells, a mapping rate cut-off of ‘MappingRate’ > 0.5 was set (**Fig. S5A**). Then, the density histogram of the ‘TotalDedupped’ (i.e. final read) metric was plotted across all cells passing the mapping rate filter, and minimum and maximum thresholds of 50,000 ($5 \times 10^4$) and 3,500,000 ($3.5 \times 10^6$) were determined (**Fig. S5B**). An mCH fraction filter of ‘mCHFrac’ > 0.2 was determined based on the density histogram of the cells that passed the final read filter (**Fig. S5C**). A mCCC fraction filter of ‘mCCCFrac’ > 0.03 was used based on the density histogram of the cells that passed the mCH fraction filter (**Fig. S5D**). Finally, a mCG fraction filter of ‘mCGFrac’ > 0.5 was used based on the density histogram of the cells that passed the mCCC fraction filter (**Fig. S5E**). An additional long-range cis contact filter of ‘CisLongContact > 6000 was also implemented on the cells passing the prior 5 filters based on the ‘CisLongContact’ density histogram (**Fig. S5F**; left column) and the effect of this filter on ‘CisShortContact’ and ‘TransContact’ distributions was assessed (**Fig. S5F**; middle and right columns). Finally, cells with low cis/trans ratios less than a cuttoff of ‘CisTransRatio’ > 5 were filtered based on the density histogram of cells passing the prior 6 filters (**Fig. S5G**). After all filters, ~200-300 poor-quality cells out of 1152 total cells per sample were removed, and the remaining cells were carried forward for downsampling (**Fig. S5H**). For each condition, cells that passed the filters described in **Fig. S5** were sorted in descending order based on ‘CisLongContact’ value. The first 750 cells were selected from this sorted list for each condition. In total, 3,000 cells across all conditions were used for downstream analysis.

##### *3C Analysis*

*Preprocessing:* After obtaining a list of 3,000 cells selected based on highest ‘CisLongContact’ values, default valid pairs files outputted by YAP were used to assemble 30kb resolution single cell matrices per autosome, resulting in 66,000 square, symmetric matrices (**Fig. S8A**). For each autosome, all 3,000 cells were scaled together to normalize for sequencing depth. Before scaling, a pseudocount of 1e-5 was added to all contact frequencies across all bins in all cells. Pseudocounted contact frequencies were normalized using size factors based on distance from the diagonal. Size factors were computed for each cell by summing the counts between pairs of bins separated by the same distance and then dividing by the geometric mean (**Fig S8A**). After scaling, single cell matrices per autosome were merged via summation into pseudo-bulk matrices, where each pseudo-bulk matrix was derived from merging counts corresponding to the downsampled 750 single cells (**Fig. S8B, S9**). To prevent balancing artifacts, poorly mapped regions were wiped from the pseudo-bulk matrices based on an hg38 150-mer alignability track generated using the GEM-mappability alignment software (*137*) (**Fig. S8B, S9**). Pairs of bins were wiped if a 50kb window centered on that pixel had a mean mappability below 25%. An additional filter, which removes high outliers, was also implemented: pairs of bins that exhibited high fold changes (>8) relative to the median value of a neighborhood defined by a 5x5 footprint were removed (**Fig. S8B, S9**). After filtering, the Knight-Ruiz matrix balancing algorithm was implemented for each matrix (**Fig. S8B, S9**). To mitigate matrix sparsity and increase signal to noise ratios, a previously published reverse-random walk (RWR) imputation from the scHiCluster package (*138*) was implemented with modifications, where the default square-root vanilla coverage (SQRTVC) normalization step was removed to avoid double balancing. After imputation, the resulting matrices consisted of very small values on the order of 1e-3. Counts were brought to a similar magnitude of counts before imputation by multiplying each value by 1000 (**Fig. S8B, S9**).

*Expected Model:* The analysis was restricted to pairs of bins within 9.99 Mb of each other from imputed 30kb resolution contact matrices. First, to account for distance dependence of 3C signal, a one-dimensional expected model, $D$, was determined from geometric mean of counts for each of the first 333 diagonals spanning 9.99 Mb (**Equation 2**):

$$D_d = \left(O_{a,b}\right) \; \forall \, d \; such \; that \; 0 \leq d \leq 333 \qquad \textbf{(2)}$$

where $D_d$ is the expected value for interactions between bins $(a, b)$ separated by $d$ bins and $O$ is the scalar multiplied imputed contact matrix, which will be referred to as the "observed" for the rest of the 3C analysis methods (**Fig. S8B, S10**).

Then, to correct the model for local architectural features, each expected value $D_{i,j}$ was multiplied by three correction factors separately (**Equations 3, 5, and 7**). These correction factors were computed by summing all bin-pairs $(a, b)$ within geometric footprints with size parameters p and w centered on bin $i, j$ in the observed and the one-dimensional expected matrix. The final factor is the ratio between the two sums. The three geometric footprints are the donut footprint (**Equations 3-4**),

$$E_{i,j}^{DF} = D_{i,j} \times \frac{\sum_{(a,b)\in DF_{i,j}} O_{a,b}}{\sum_{(a,b)\in DF_{i,j}} D_{b-a}} \qquad \textbf{(3)}$$

$$DF_{i,j} = \{(a,b) \mid (|a-i| \leq w) \wedge (|b-i| \leq w) \wedge (a \neq i) \wedge (b \neq j) \wedge ((|a-i| > p) \vee (|b-j| > p))\} \qquad \textbf{(4)}$$

the lower left footprint (**Equations 5-6),**

$$E_{i,j}^{LLF} = D_{i,j} \times \frac{\sum_{(a,b)\in LLF_{i,j}} O_{a,b}}{\sum_{(a,b)\in LLF_{i,j}} D_{b-a}} \qquad \textbf{(5)}$$

$$LLF_{i,j} = \{(a,b) \in DF_{i,j} \mid (a < i) \wedge (b < j)\} \qquad \textbf{(6)}$$

and the upper triangle footprint (**Equations 7-8**),

$$E_{i,j}^{UTF} = D_{i,j} \times \frac{\sum_{(a,b)\in UTF_{i,j}} O_{a,b}}{\sum_{(a,b)\in UTF_{i,j}} D_{b-a}} \qquad \textbf{(7)}$$

$$UTF_{i,j} = \{(a,b) \in DF_{i,j} \mid b - a \geq j - i\} \qquad \textbf{(8)}$$

respectively. The size parameters of all geometric footprints were p = 2 and w = 6. A footprint must contain greater than 0.25 finite values for an expected value to be computed. The final expected value of bin-pair $i, j$ $E_{i,j}$ was computed based on interaction distance. When the interaction distance was less than 180kb, the $E_{i,j}^{UTF}$ was the final expected value. When the interaction distance was greater than 180kb but less than 9.99 Mb, the final expected value was computed by finding the maximum expected value between the donut and lower left footprints (**Equation 9**, **Fig. S8B, S10**).

$$E_{i,j} = \begin{cases} E_{i,j}^{UTF}, & for\ b-a \leq 6 \\ max\left(E_{i,j}^{DF}, E_{i,j}^{LLF}\right), & for\ 6 < b-a < 333 \end{cases} \quad \textbf{(9)}$$

*P-value Calculation:* To compare the imputed count $X_{i,j}$, with its final expected value $E_{i,j}$, a p-value $P_{i,j}$ was computed by testing the null hypothesis that $X_{i,j}$ was less than or equal to a Poisson-distributed random variable $X'_{i,j}$ with mean $E_{i,j}$ (**Equation 10, Fig. S8B, S10**):

$$P_{i,j} = P\left(X_{i,j} \leq X'_{i,j}\right); \quad X'_{i,j} \sim Poisson\left(E_{i,j}\right) \quad \textbf{(10)}$$

*Loop Clustering*: After computing $P_{i,j}$, clusters of nearby significant bin-pairs were identified. First, an initial set of significant bin-pairs was identified using: (1) a p-value threshold $P_{i,j} \leq 0.075$, an imputed contact value threshold $S_{i,j} \geq 8$ and an observed over expected fold-change threshold $FC \geq 1.5$. Single significant pixels without neighbors were removed to reduce false positives. The initial calls had larger footprints than what was reflected in the observed matrices. Therefore, the calls were further refined by iterating two times through smaller p-value thresholds ranging from 0.075 to 0.04. Finally, to avoid calling loops near the diagonal of the contact matrix, all clusters containing at least 1 pixel within 4 bins of diagonal were removed (**Table S8, Fig. S8B, S10**).

*Differential Cluster Classification*

To identify differential loops, comparisons were performed between the TTX treatment, 2 hours post-KCl treatment, and 5 days post-KCl conditions. Overlapping clusters across all conditions were merged if they contained at least 1 exactly overlapping bin-pair. These merged clusters formed set $C$. The mean of pseudo-counted-observed over pseudo-counted-expected values from bin-pairs (a,b) within cluster $C_k$ for each condition P was computed (**Equation 10**).

$$\mu_k^P = \frac{1}{|C_k|} \sum_{(a,b) \in C_k} \frac{O_{a,b}^P + 0.1}{E_{a,b}^P + 0.1} \quad \textbf{(11)}$$

Mean observed over expected values, $\mu_k^P$, across TTX (condition A), 2 hours post-KCl (condition B), 2 hours post-KCl+Nim (condition C) and 5 days post-KCl (condition D) were then used to determine a set of five sequential fold-change thresholds. The fold-change between TTX and 2 hours post-KCl was defined as

$$F_k^{AB} = \frac{\mu_k^B + 0.1}{\mu_k^A + 0.1} \quad \textbf{(12)}$$

The fold-change between TTX and 2 hours post-KCl+Nim was defined as

$$F_k^{AC} = \frac{\mu_k^C + 0.1}{\mu_k^A + 0.1} \quad \textbf{(13)}$$

The fold-change between TTX and 5 days post-KCl was defined as

$$F_k^{AD} = \frac{\mu_k^D + 0.1}{\mu_k^A + 0.1} \quad \textbf{(14)}$$

The fold-change between 5 days and 2 hours post-KCl was defined as

$$F_k^{DB} = \frac{\mu_k^B + 0.1}{\mu_k^D + 0.1} \quad \textbf{(15)}$$

The fold-change between 2 hours post-KCl and 2 hours post-KCl+Nim was defined as

$$F_k^{BC} = \frac{\mu_k^C + 0.1}{\mu_k^B + 0.1} \quad \textbf{(16)}$$

Differential classes were assigned based on logic gates related to fold-change and mean observed over expected value:

- transiently gained when $\mu_k^A < 1.6, \mu_k^B \geq 1.625, \mu_k^D < 1.6,\ F_k^{AB} \geq\ 1.3$, and $F_k^{DB} \geq 1.3$,
- transiently lost when $\mu_k^A \geq 1.625, \mu_k^B < 1.6, \mu_k^D \geq 1.625,\ F_k^{AB} \leq \frac{1}{1.3},\ \frac{1}{1.3} F_k^{AD} \leq 1.3$, and $F_k^{DB} \leq \frac{1}{1.3}$
- persistently gained when $\mu_k^A < 1.6, \mu_k^B \geq 1.625, \mu_k^D \geq 1.625,\ F_k^{AB} \geq\ 1.3, F_k^{AD} \geq\ 1.3$, and $\frac{1}{1.3} <\ F_k^{DB} < 1.3$
- persistently lost when $\mu_k^A \geq 1.625, \mu_k^B < 1.6, \mu_k^D < 1.6, F_k^{AB} \leq \frac{1}{1.3}, F_k^{AD} \leq \frac{1}{1.3}$
- invariant when $\mu_k^A \geq 1.625,\ \mu_k^B \geq 1.625,\ \mu_k^D \geq 1.625$
- unclassified otherwise.

The number of loops in the initial persistence classes were small, but it was also observed that some of the invariant clusters still exhibited increasing or decreasing loop strength between TTX, 2 hour post-KCl, and 5 days post-KCl. Clusters classified as invariant across the 3 conditions were further stratified as increasing in strength when $F_k^{AB} \geq 1.3,\ F_k^{AD} \geq 1.3$ and decreasing in strength when $F_k^{AB} \leq \frac{1}{1.3},\ F_k^{AD} \leq \frac{1}{1.3}$ Invariant loops that met these conditions were moved from the original invariant sub-class and pooled together with the original persistent gained and lost loop sub-classes, respectively, for downstream analyses. Additionally, clusters with two significant pixels were only kept if both anchors colocalized with any H3K27ac CUT&Tag peaks called in any condition to further mitigate noisy loop calls (**Table S9**).

*Loop Stratification with respect to Regulatory Elements:* Each loop in each persistence sub-class was classified as promoter-promoter, enhancer-promoter, enhancer-enhancer, structural, or ambiguous. Using interlap (v. 0.2.7) (*139*), genomic coordinates of each loop anchor was intersected with coordinates of exons, UTRs, additive promoter regions spanning 600 bp upstream to 200 bp downstream of the merged TSS (TSSs within 1 basepair), 200 bp regions centered on the midpoint of differential enhancers, and 200bp regions centered on original H3K27ac peaks merged across all conditions.

For the persistent and transient gained loops, enhancer regions were derived from 2 hours post KCl-specific (activity-induced) non-coding H3K27ac peaks with respect to TTX. For the persistent and transient lost loops, enhancer regions were derived from TTX-specific (activity-decommissioned) non-coding H3K27ac peaks with respect to 2 hours post-KCl. For the remaining invariant loops, enhancer regions were derived from a concatenated list of TTX-specific non-coding H3K27ac peaks, 2 hours post KCl-specific non-coding H3K27ac peaks, and invariant non-coding H3K27ac peaks.

A loop was classified as promoter-promoter (P-P) if both anchors intersected at least one promoter but neither intersected any enhancers. A loop was classified as enhancer-promoter if one anchor intersected at least one promoter and no enhancers and the other anchor intersected at least one enhancer and no promoters. However, because of the low resolution of the 3C modality of the snm3c-seq3 assay, called loops often contained a combination of enhancers and promoters within the same anchor. Therefore, the definition of an enhancer-promoter loop was expanded to include the following scenarios:

- Both anchors intersected at least one promoter and one enhancer (PE-PE).
- One anchor intersected at intersected at least one promoter and one enhancer and the other anchor intersected at least one promoter and no enhancers (PE-P).
- One anchor intersected at least one promoter and one enhancer and the other anchor intersected at least one enhancer and no promoters (PE-E).

A loop was classified as enhancer-enhancer (E-E) if both anchors intersected at least one enhancer and neither intersected any promoters. A loop was classified as “structural” if neither anchor intersected H3K27ac peaks called in any condition, promoters, exons, or UTRs and both anchors intersected at least one CTCF peak called in any condition. Loops that did not meet any of these specific criteria were assigned an “ambiguous” label.

Next, promoter-promoter (P-P), enhancer-promoter (P-E ,PE-PE, PE-P, and PE-E), and enhancer-enhancer (E-E) loops were further classified across all regulatory elements at the anchor that contains promoters (“promoter anchor”) and the anchor that contains enhancers (“enhancer anchor”). Given the ambiguity of which anchor is the “promoter anchor” and which is the “enhancer anchor” when the loop has a PE-PE, PE-P, or PE-E configuration, the following procedures were used for anchor assignments:

- PE-PE loop anchors were assigned twice. During the first assignment, the upstream anchor was the “promoter anchor” and the downstream anchor was the “enhancer anchor.” During the second assignment, the labels were switched where the upstream anchor was the “enhancer anchor” and the downstream anchor was the “promoter anchor.”
- For PE-P loops, the anchor without any enhancers (P) was the “promoter anchor” and the anchor with both enhancers and promoters (PE) was the “enhancer anchor.”

- For PE-E loops, the anchor without any promoters (E) was the "enhancer anchor" and the anchor with both enhancers and promoters (PE) was the "promoter anchor."

For elements associated with a loop, a promoter passed if it was on a "promoter anchor" and an enhancer passed if it was on an "enhancer anchor." Conversely, a promoter was removed if it was on an "enhancer anchor" and an enhancer was removed if it was on a "promoter anchor" Once lists of "passing" additive promoter and enhancer windows overlapping classified loop anchors were obtained, the additive promoters were divided into "proximal" promoters defined as plus or minus 200 bp of the merged TSSs (see ***Nuclear RNA Seq Computational Analysis)***, and "distal" promoters, defined as the upstream bound of the additive promoter to the upstream bound of the proximal promoter. These promoter sub-windows and the size matched enhancer windows were used for downstream analyses.

*Determining Dependence on CREB phosphorylation*
PE loops in all activity-induced loop classes were classified as either 'pCREB-dependent' or 'pCREB-independent' based on the mean observed / expected signal of 2 hours post-KCl and 2 hours post-KCl + Nim to control for the possibility of TTX removal affecting looping. Loops in the transient gained, transient lost, persistent gained, and persistent lost categories were evaluated for pCREB dependence (**Table S10, S11**). This was based on the following criteria:

- Transient Gained: 'pCREB dependent' if $\mu_k^C$ < 1.6 and $F_k^{BC}$ < 1/1.3
- Transient Lost: 'pCREB dependent' if $\mu_k^C$ > 1.625 and $F_k^{BC}$ >1.3
- Persistent Gained: 'pCREB dependent' if $F_k^{BC}$ < 1/1.3
- Persistent Lost: 'pCREB dependent' if and $F_k^{BC}$ > 1.3

*Defining Unlooped Regulatory Elements*
Unfiltered lists of unlooped elements were obtained by subtracting any looped promoter or enhancer, including elements on ambiguous loops, from the set of RefSeq hg38 promoters or H3K27ac enhancers, respectively.

*H3K27ac Signal at Looped Regulatory Elements:*
To validate the persistence loop class definitions, the H3K27ac CUT&Tag signals for the TTX, 2 hours post-KCl treatment, and 5 day post-KCl treatment conditions were compared at looped proximal promoters, looped distal promoters, looped enhancers. Invariant proximal promoters, invariant distal promoters, invariant enhancers, and unlooped enhancers were further stratified by H3K27ac peaks that were KCl-specific, TTX-specific, or invariant. If one of these regulatory elements intersected multiple H3K27ac peaks, then only one H3K27ac peak was assigned to that regulatory element with the priority KCl > TTX > invariant.

H3K27ac signal at regulatory elements was calculated using the computeMatrix function from deeptools (version 3.5.1) (*135*). The input-subtracted bigWig for the 5 day post-KCl treatment was used as the signal files for the -S flag, and the regulatory elements were used as region files for the -R flag. An additional set of H3K27ac signal matrices were calculated using the log2 fold change of 2 hours post-KCl to TTX, 2 hours post-KCl + Nim to TTX, and 2 hours post-KCl to 2 hours post-KCl + Nim. Log2 fold change bigWigs were generated using the bigwigCompare function from deeptools (version 3.5.1) (*135*) with the flags --operation log2 --outFileFormat "bigwig"

The reference-point argument "center" was used for proximal promoters and enhancers and the reference-point argument "TES" for distal promoters. H3K27ac scores were calculated in a 1500 bp region upstream and downstream of the reference-point using a 10 bp bin size. The regions were sorted by total 5 day post-KCl H3K27ac signal and saved in descending order. The H3K27ac signal was calculated for TTX and 2 hr post-KCl treatment using input subtracted bigwigs for each condition and the region files sorted by 5 day post-KCl treatment H3K27ac signal. Regions in the log2 fold change matrices were sorted in descending order by total signal in the log2 fold change of 2 hour post-KCl treatment to TTX.

After calculating the H3K27ac signal across three conditions for looped regulatory elements, the plotHeatmap function from deeptools (version 3.5.1) (*135*)was used to visualize the aggregate H3K27ac signal over looped regulatory elements with the flags --yMin 0 --yMax 3 --zMin 0 --zMax 3 --samplesLabel "" --regionsLabel "" --plotTitle "" --xAxisLabel "" --yAxisLabel "" --heatmapHeight 4 --whatToShow "plot and heatmap" --dpi 800 --legendLocation none. The Purples color map was used for input subtracted signal and the seismic color map was used for log2 fold change signal.

***DNA Methylation Analysis***

*Pre-processing:* The ALLC files from the YAP pipeline across all single cells were imported into ALLCools (v. 1.1.0) and then filtered and clustered according to published documentation (*65*).

***Setting parameters at every computational snm3C-seq3 step***

*Filtering bins of low quality:* 100 kb resolution bins overlapping ENCODE blacklist regions (https://github.com/Boyle-Lab/Blacklist/) were removed. Bins overlapping chromosomes X, Y, and M were also removed. Then the average coverage across cells for every bin was calculated. Low-quality and unusually high coverage bins were also removed, as they may represent low-mappability regions and alignment artifacts. Min and max cut-offs of 500 and 1400 were used based on the distribution of average coverage (**Fig. S6A**).

*Calculating methylated-cytosine fractions and cell-to-cell normalization:* With total counts and counts supporting methylation, methylation fractions were independently calculated for CpG and non-CpG methylation by dividing the number of methylated reads per cytosine by the total number of reads per cytosine, and cell-to-cell normalization was performed using the beta-binomial distribution method as described previously (*140*). Then, normalized values were calculated similar to the counts per million reads (counts) of single-cell RNA-seq analysis. Boxplots of methylated fractions are shown before (**Fig. S6A)** and after (**Fig. S6B)** cell-to-cell normalizations at 100 kb resolution.

*Calculating highly variable features (HVFs):* Selection of highly variable features has been crucial in single-cell analysis (*64-66*). In snm3C-seq3, there is a non-linear correlation between methylated fraction, average coverage, and dispersion. Since removing the non-linear correlation is important before selecting HVFs, the non-linear correlation was independently removed for CpG and non-CpG methylation using a non-linear Support Vector Machine for Regression (SVR). HVFs were independently identified for CpG and non-CpG methylation. Three options were selected based on the top 25%, 50%, and 75% of features remaining after all filters (**Fig S7A-B**).

*Principal component analysis (PCA):* PCA was independently performed for CpG and non-CpG methylation PCA as described in Xu, Chandradoss et al. 2025 (*126*). Then, the top 25 significant Principal Components of CpG and non-CpG methylation were concatenated after normalizing for standard deviation.

*Choosing the Leiden Resolution & embedding:* To choose the Leiden Resolution, a *k-nearest* neighbor (KNN) graph was built, clusters were defined using the Leiden Algorithm, and cells were embedded using UMAP. Briefly, a *k-nearest* neighbor (KNN) graph was obtained after concatenating the significant dimensions from the PCA of CpG and non-CpG methylation. Since the resolution parameter affects the Leiden Algorithm in defining the number of clusters, 3 different resolution parameters, such as 0.06, 0.125, and 0.25 at 100 kb bin size, were tried **(Fig. S7).** It was evident that increasing the Leiden Resolution increased the number of clusters defined despite the homogeneous distribution of cells.

***Differentially Methylated Loci (DML) Analysis***

Upon confirming that cells did not cluster across or within condition, our pre-identified 750 single cell ALLC files were merged into pseudo-bulk ALLC files with the 'allcools merge' command (version 1.1.0) for each condition (*64-66*). To focus our differential methylation analysis on CpG sites, the CpG sites from each condition were extracted using the 'allcools extract' command (version 1.1.0) (*64-66*).

To facilitate analysis of differentially methylated sites across our four conditions, a series of filtering steps were applied to our data. Blacklist sites and nonstandard chromosomes were removed, followed by the removal of any cytosine that did not have a coverage of at least three reads in at least one condition from all four conditions (**Fig. S13**).

To confirm that our CpG methylation matched expected patterns from literature (*112*), DNA methylation percentage at all CpG sites was assessed at High-CpG density promoters (HCPs), Low-CpG Density Promoters (LCPs), highly-conserved non-coding elements (HCNEs), long terminal repeats (LTRs), short interspersed nuclear elements (SINEs), and all other genomic features (*112*) (**Fig. S14**). Promoters were defined as HCPs if they contained a 500bp interval within -2kb to 200bp of the TSS with a (G+C) fraction of ≥ 0.55 and an observed GC content to expected GC content (O/E) ratio of ≥ 0.6 (*112, 141*). The expected GC content for a 500 bp window was defined as $(\text{GC content} / 2)^2$ (*142*). Windows were tested using a sliding interval of 100 bp. Promoters were defined as LCPs if they contained no 500 bp window with the CpG O/E ratio ≥ 0.4 (*112, 141*). HCNE annotations were created by downloading phastCons conservation score from the UCSC genome browser (hg38) and defining HCNE as CpGs not in exons with phastCons greater than 0.75. LTRs and SINEs annotations taken from RepeatMasker annotations were downloaded from the UCSC genome browser (hg38). "Other genomic features" were defined as any CpG that did not fall into these categories.

To assess our DNA methylation profiles at CpG islands, the cpgIslandExt.txt.gz file was downloaded from the UCSC genome annotation database for hg38 (https://hgdownload.soe.ucsc.edu/goldenPath/hg38/database/) on 1/16/2025. The methylation fraction of each of our four conditions was analyzed at large (-4200 bp to -200 bp from TSS) and small (-200 bp to +200 bp from TSS) promoter windows at CpGs within and not within CpG islands (**Fig. S15**). Further, the methylation fraction of each of our conditions was analyzed at large (-2000 bp to +2000 bp from H3K27ac peak center) and small (-200 bp to +200 bp from H3K27ac peak center) at CpGs within and not with CpG islands (**Fig. S16**).

To identify DMLs in our dataset, the Dispersion Shrinkage for Sequencing Data (DSS) package (v. 2.54.0) from Bioconductor was utilized (*143-145*). Our filtered, pseudo-bulk ALLC files were converted into a format compatible with the DSS package by converting each ALLC file into a file with four columns representing chromosome, CpG position, number of total reads, and number of methylated reads. The bsseq package (v. 1.42.0) was used to create a Bsobj object via the 'makeBSseqData' function. The Bsobj object was then used as an input to the 'DMLtest' function to perform a Wald test to identify Differentially Methylated Loci (DMLs) with smoothing set to TRUE, smoothing.span set to 50 bp, ncores=1, and all other parameters set to default. Our experimental group for a given comparison was set to group1, and our control group was set to group2. This comparison was run for 2-hours post KCl vs TTX and 5 days post KCl vs TTX, and the outputs were saved as TSVs. The TSVs were then converted into bed files by adding inserting a column equal to the CpG coordinate plus one as the third column. For each BED file, the raw methylation fraction for the control and experimental conditions was added in as new columns using bedtools map (v2.26.0) (*133*).

To identify DMLs, we took both the DSS-generated p-value (*143-145*), and the difference in methylation fraction between the experimental and control conditions into account to remove potential false positives. To do so, we manually identified CpGs with i) a DSS-outputted p-value of < .001 and ii) a difference in methylation fraction of at least 0.2 in magnitude between the experimental and control conditions using a custom awk command. *Hypermethylated DMLs* were identified as those with a difference in methylation fraction greater than 0.2 for the experimental condition minus the control condition, and *Hypomethylated DMLs* were identified as those with a difference in methylation fraction less than -0.2 for the experimental condition minus the control condition.

To stratify activity induced DMLs from the *2-hours post KCl condition vs TTX comparison* as Transient or Persistent DMLs, we took the presence of a CpG in the 5 days post-KCl condition vs TTX comparison into account in the following manner using bedtools intersect (v2.26.0) (*133*):

- Persistently hypermethylated DMLs were defined as CpGs that were hypermethylated in both the *2 hours post-KCl vs TTX* comparison and the *5 days post-KCl vs TTX* comparison.
- Transiently hypermethylated DMLs were defined as CpGs that were only hypermethylated in the *2 hours post-KCl vs TTX* comparison.
- Persistently hypomethylated DMLs were defined as CpGs that were hypomethylated in both the *2 hours post-KCl vs TTX* comparison and the *5 days post-KCl vs TTX* comparison.
- Transiently hypomethylated DMLs were defined as CpGs that were only hypermethylated in the *2 hours post-KCl vs TTX* comparison.

To focus our analysis only on activity-induced DMLs, DMLs that were only present in the *5 day post-KCl vs TTX* comparison were dropped from downstream analysis.

***Differential Methylation at Regulatory Elements Analysis***

Lists of distal promoters, proximal promoters, and enhancers split by looping class were analyzed for DMLs. Unlooped enhancers and invariant enhancers were stratified by whether the corresponding H3K27ac peak was TTX-specific (activity-decomissioned), 2 hours post-KCl-specific (activity-induced), or invariant. Invariant promoters were stratified by whether or not they were looped to enhancers that were TTX specific, 2 hours post-KCl specific, or invariant. All groups of regulatory elements were split by whether or not there was overlap with a CTCF peak defined as described previously (***ChIP-seq Analysis)*** and whether or not there was overlap with a

CpG island, using bedtools intersect (version 2.30.0) (*133*). Each group of promoters therefore categorized as one of four options i) +CTCF + CpG island; ii) + CTCF – CpG island; iii) – CTCF +CpG island; iv) -CTCF – CpG island; and each enhancer was categorized as one of two options i) + CTCF; ii) -CTCF.

Each list of regulatory elements was intersected by the four types of DMLs previously described (***Differentially Methylated Loci (DML) Analysis)*** in sequence such that each element could only appear in one of the four DML types using the priority ranking persistent hyper-DML > transient hyper-DML **>** persistent hypo-DML > transient hypo-DML. To plot the methylation distribution across various regulatory elements, individual CpGs in the TTX, 2 hours post-KCl, and 5 days post-KCl condition were mapped to regulatory elements using bedtools map (version 2.30.0) (*133*).

***Gene Expression Persistence Class Identification***

To identify activity-induced or activity-downregulated genes, genes that were upregulated (FC > 0, FDR < .05) or downregulated (FC < 0, FDR <.05) either 2 hours post-KCl or 5 hours post-KCl treatment were identified as described in ***Nuclear RNA Seq Computational Analysis***. Genes were then filtered to remove any gene whose promoter window intersected with a DML using bedtools intersect (*133*). Genes that were upregulated at either 2 hours or 5 hours post KCl were identified as activity-induced (Red); genes that were downregulated at either 2 hours or 5 hours post KCl were identified as activity-downregulated (Blue); all other genes with reads were identified as activity-invariant (Grey) (**Table S12**).

Upon identifying Red Blue and Grey classes, genes were further stratified by their behavior 5 days after stimulation (**Tables S12**). Genes that were Activity Induced were classified as:

- 'Red→Red' if their average counts 5 days post KCl stimulation were 1.2-fold greater than TTX
- 'Red→Grey' if their average counts 5 days post KCl stimulation were between 1/1.2 and 1.2 FC greater than TTX
- 'Red→Blue' if their average counts 5 days post KCl stimulation were less than 1/1.2.

Similarly, genes that were Invariant were classified as 'Grey→Red', 'Grey→Grey' and 'Grey→Blue', and genes that were TTX-specific were classified as 'Blue→Red' 'Blue→Grey' and 'Blue→Blue' using the same logic.

***Odds Ratio Calculation for Looping Behavior and Activity-regulated Gene Expression Outcomes***

We calculated the distribution of odd ratios (ORs) and used Fisher's exact test p-values. We created with a 2 x 2 contingency table with counts from two group of genes based on looped classes and then split the genes based on their expression changes from the TTX condition to the KCl 2 hours and 5 hours post-stimulation to the 5 days post-stimulation timepoint.

***Gene Ontology (GO) Analysis***

Gene ontology was conducted as previously described (*5, 72*) using WebGestalt (*146*) with 'Method of Interest' set to *Over-Representation Analysis*, 'Organism of Interest' set to *Homo sapiens*, 'Functional Database' set to *geneontology biological process noRedundant*, 'Select Reference Set' set to *genome*, 'Analyte Type' set to *Gene/Protein*, and 'ID Type' set to *Gene symbol* with a manually entered 'Input ID List' (**Table S13**).

***Computational Analysis of snRNA-seq Data***

Single-nucleus RNAseq libraries generated from 10X genomics v3.1 were analyzed as previously described with minor modifications (*72*). The cellranger library (version 8.0.1) (*147*) was used to incorporate the *Ngn2* transgene into the standard GRCh38 reference genome and align reads to the resultant reference genome.

First, NCBI Refseq GTF and FASTA files corresponding to the hg38 genome build were downloaded on February 6th, 2021 from the UCSC Genome Browser. (https://hgdownload.soe.ucsc.edu/goldenPath/hg38/bigZips/). The original GTF file was filtered for non-polyA transcripts using 'cellranger mkgtf' with --attribute=gene_biotype:protein_coding. The *Ngn2* sequence and annotations were appended to the original GRCh38 FASTA and filtered GTF file, respectively. The resultant merged FASTA and GTF files were used in "cellranger mkref" to generate a GRCh38 reference genome with the *Ngn2* transgene.

Next, reads for each sample were aligned to the GRCh38-*Ngn2* reference genome using cellranger count. The output of the previous mkref was used for the --transcriptome flag in cellranger count, and the --create-bam flag was set to True. The resultant nuclei x gene matrices were loaded into R (version 4.3.3) with Seurat (version 5.1.0) (*148*).

To filter for doublets and debris, barcodes that fell into the highest 99.7 percentile and lowest 0.3 percentile of counts per barcode, respectively, were removed (**Fig. S19A-B, S20A-B**). Then, nuclei with low complexity, defined as nuclei with 0s in at least 95% of genes were removed (**Fig. S19C, S20C**). Next, we filter the condition with the least nuclei remaining, from 373 to the 300 nuclei with highest genes with counts (**Fig. S19D, S20D**). Then, we used nearest neighbor matching to find nuclei from the other two conditions that match the 300 nuclei based on their number of genes with counts (**Fig. S19E, S20E**). Next, genes that were lowly expressed or detected, genes that had a count of 0 in at least 75% of cell in all conditions were excluded (**Fig. S19F, S20F**). Then, for each nucleus, counts for gene were normalized by multiplying by the 75th percentile nucleus' sum of counts in the double stimulus condition and dividing by the 75th percentile nucleus' sum of counts in the condition to which the nucleus belongs (**Fig. S19G, S20G**). Finally, to compare genes across conditions, we calculated the proportion of cells with counts > 0 in each condition.

To relate the bulk RNA-seq gene expression patterns to the single-cell RNA-seq data, we assigned genes by their behavior after initial stimulation and 5 days after stimulation as observed by bulk RNA-seq data (***Gene expression persistence class identification***) to generate nine classifications. For genes with multiple genes, we assigned genes to an expression pattern hierarchically in this order: Red→Red, Red→Grey, Red→Blue, Grey→Red, Grey→Grey, Grey→Blue, Blue→Red, Blue→Grey, and Blue→Blue. We considered genes with a difference of at least 0.15 in proportion of cells with expression as upregulation and a difference of up to -0.15 as downregulation. We labeled genes as 'no change' between the conditions, if the difference in proportion of cells was between -0.15 and 0.15 and the gene was detected in at least 5% of nuclei in each condition.

***Persistent Enhancer Memory Stratification***

We stratified distal H3K27ac peaks that were present at 2 hours post stimulation neurons but not in TTX silenced neurons (activity-induced enhancers) into varying levels of persistence in 5 days post stimulation neurons. We defined three levels of persistence based on the ratio of H3K27ac signal at 5 days post stimulation to H3K27ac signal at 2 hours post stimulation: a ratio ≥ 0.5 was high persistence; 0.5 > ratio ≥ 0.35 was medium persistence; 0.35 > ratio ≥ 0.2 was low persistence; and ratio < 0.2 was defined as 'No Memory' persistence.

***Odds Ratio Estimation through Bootstrapping for Enhancer Memory***

We estimated the distribution of odd ratios (ORs) and p-values by bootstrapping with resampling. We began with a 2 x 2 contingency table with counts from two group of enhancers split by whether they belong to one of two persistence strata with one of the strata always being the 'none' persistence strata. For each draw, we sampled from the original distribution of each group, the original number of observations per group with replacement. From the resulting 2 x 2 contingency tables, we calculated an odds ratio and Fisher's exact test p-value. If draws resulted in a 0 in the denominator of the odds ratio, we redrew that draw up to 100 times until an odds ratio, until we obtained an odds ratio with a non-zero denominator. If we did not produce an odds ratio with a non-zero denominator, we recorded that draw as such. We performed 10,000 draws per comparison and report the median ORs and p-value. We calculated an empiric 95% confidence interval by using the $2.5^{th}$ percentile OR and $97.5^{th}$ percentile OR from the 10,00 draws.
